# Leveraging eQTLs to identify individual-level tissue of interest for a complex trait

**DOI:** 10.1101/674226

**Authors:** Arunabha Majumdar, Claudia Giambartolomei, Na Cai, Tanushree Haldar, Tommer Schwarz, Michael J. Gandal, Jonathan Flint, Bogdan Pasaniuc

## Abstract

Genetic predisposition for complex traits often acts through multiple tissues at different time points during development. As a simple example, the genetic predisposition for obesity could be manifested either through inherited variants that control metabolism through regulation of genes expressed in the brain, or that control fat storage through dysregulation of genes expressed in adipose tissue, or both. Here we describe a statistical approach that leverages tissue-specific expression quantitative trait loci (eQTLs) corresponding to tissue-specific genes to prioritize a relevant tissue underlying the genetic predisposition of a given individual for a complex trait. Unlike existing approaches that prioritize relevant tissues for the trait in the population, our approach probabilistically quantifies the tissue-wise genetic contribution to the trait for a given individual. We hypothesize that for a subgroup of individuals the genetic contribution to the trait can be mediated primarily through a specific tissue. Through simulations using the UK Biobank, we show that our approach can predict the relevant tissue accurately and can cluster individuals according to their tissue-specific genetic architecture. We analyze body mass index (BMI) and waist to hip ratio adjusted for BMI (WHRadjBMI) in the UK Biobank to identify subgroups of individuals whose genetic predisposition act primarily through brain versus adipose tissue, and adipose versus muscle tissue, respectively. Notably, we find that these individuals have specific phenotypic features beyond BMI and WHRadjBMI that distinguish them from random individuals in the data, suggesting biological effects of tissue-specific genetic contribution for these traits.

## Introduction

Multiple clinical, pathologic, and molecular lines of evidence suggest that many phenotypes and diseases show heterogeneity and can be viewed as a collection of multiple traits (i.e. subtypes) in the population [1–5]. Traditional subtype identification has relied on detecting biomarkers or subphenotypes that distinguish subsets of individuals in a biologically meaningful way. For example, breast cancer has two well-known subtypes, estrogen receptor positive and negative [6–8]. Patients with psychiatric disorders can have different severities [9]. With the advent of large scale genome-wide association studies (GWAS) that have robustly identified thousands of risk variants for complex traits, multiple approaches have investigated the use of genetic risk variants to define classes of individuals that show genetic heterogeneity across subtypes [10–15]. For example, autism can be subtyped by grouping together individuals with recurrent mutations in the same autism-associated gene [10, 12]; type 2 diabetes can be subtyped using clusters of genetic variants previously associated with the disease [13]. Other examples include adiposity traits such as body mass index (BMI), waist-to-hip ratio (WHR), and WHR adjusted for BMI (WHRadjBMI), that can be subtyped based on genetic variants with distinct patterns of fat depots and metabolism [14]. Genetic subtyping offers an advantage over phenotypic subtyping in that germline genetic characteristics are more stable than phenotypic characteristics of an individual [12,13].

A significant component of the genetic susceptibility of complex traits is mediated through genetic control of gene expression in one or multiple tissues [16–18], with several studies highlighting the relevance of tissue or cell type specific biological mechanisms underlying the pathogenesis of complex traits [19–25]. Such studies rely on integration of expression quantitative loci (eQTLs) with GWAS data in a tissue or cell type specific manner to prioritize tissues and cell types that are relevant for a given complex trait. For example, a recent study [23] proposed a novel framework to identify the disease-relevant tissues by employing the stratified LD score regression to investigate whether the collection of genomic regions surrounding the set of tissue-specific expressed genes are enriched for the disease heritability. Such studies have often identified multiple tissues relevant to a given trait in the population (e.g., liver, pancreas and thyroid tissues for total cholesterol [22]; connective skeletal muscle and adipose for WHRadjBMI [23,26]; brain and adipose for BMI [21,23,27], etc.). Transcriptome-wide association studies (TWAS) have identified novel associations between the genetic component of tissue-specific expression of a gene and a complex trait [17,28–30], and have often discovered significantly associated genes for a trait in multiple tissues which demonstrates the tissue-specific genetic contribution to the trait across different tissues.

This motivates us to ask an interesting follow-up question that, if a phenotype has two or more relevant tissues in the population, how we can quantify the tissue-wise genetic contribution to the trait for a given individual. A method addressing this objective would also allow us to explore the possibility of prioritizing a relevant tissue for the phenotype of an individual. A recent study [31] has proposed to identify disease subtypes by integrating clinical features related to the disease and gene expression profiles across patients. Using a multi-view clustering algorithm they classified patients into subgroups where each subgroup has a distinct pattern of gene expression predicted based on cis-SNPs’ genotypes and various clinical features related to the disease combined together. However, their approach does not investigate tissue-specific genetic predisposition to the phenotype for a given individual. Since, tissue-specific genetics plays an important functional role in the tissue-specific biological processes, we aim to explicitly quantify tissue-wise genetic contribution to a phenotype at an individual-level, and identify subgroups of individuals who are homogeneous with respect to the effect of their tissue-specific genomic profile on the trait.

In this work, we present a statistical approach that integrates tissue-specific eQTLs (i.e., eQTLs for tissue-specific genes) with genetic association study data for a complex trait to probabilistically quantify the tissue-wise genetic contribution to the phenotype of each individual in the study. Following previous studies [20, 23] we characterize a tissue by a set of genes specifically over-expressed in it, and develop our model based on the eQTLs for such genes in the tissue. We focus on traits where multiple tissues have been implicated by previous studies (e.g. brain and adipose for BMI [21,23,27]), and hypothesize that the genetic predisposition to the trait for a subgroup of individuals can act primarily through a specific tissue. Assuming that two tissues are relevant for the trait in the population, there are three possibilities for a given individual: the genetic susceptibility of the trait is mediated primarily through first tissue, second tissue, or both tissues. That is, for one group of individuals the genetic predisposition to BMI acts through regulation in the brain, for another group of individuals the genetic predisposition acts through adipose, and for the remaining individuals it does not act through a specific tissue. In our study, we focus on the first two groups of individuals and examine the characteristics that distinguish them from the remaining population. We propose eGST (eQTL-based Genetic Sub-Typer), an approach that estimates the posterior probability that whether a relevant tissue can be prioritized for an individual’s phenotype or not, based on individual-level genotype data of tissue-specific eQTLs and marginal phenotype data. eGST implements a Bayesian framework of mixture model by employing a computationally efficient maximum a posteriori (MAP) expectation-maximization (EM) algorithm to estimate the tissue-specific posterior probabilities per individual.

We perform extensive simulations using real genotypes from the UK Biobank and show that eGST accurately infers the simulated tissue of interest for each individual. We also show that a Bayesian framework of the mixture model performs better than the corresponding frequentist framework. By integrating expression data from the GTEx consortium [16,18], we apply eGST to two obesity related measures (BMI and WHRadjBMI) in the UK Biobank [32,33]. We consider brain and adipose tissues for BMI to identify two subgroups (in aggregate 25192 individuals, 7.5% of total sample size), one with adipose and the other with brain tissue-specific genetic contribution, respectively. Similarly, we consider muscle and adipose tissues for WHRadjBMI to identify two subgroups (total 19041 individuals, 5.7% of all individuals) with muscle- or adipose-specific genetic contribution. Interestingly, the subgroups of individuals classified into each tissue show distinct genetic and phenotypic characteristics. 85 out of 106 phenotypes tested in the UK Biobank were differentially distributed between the BMI-adipose (or BMI-brain) group of individuals and the remaining population, with 72 out of 85 remaining significant after adjusting for BMI. For example, diabetes proportion, various mental health phenotypes, alcohol intake frequency, and smoking status were differentially distributed between one or both of the tissue-specific groups of BMI and the remaining population. Overall, our results suggest that tissue-specific eQTLs can be successfully utilized to quantify tissue-wise genetic contribution to a complex trait and prioritize the tissue of interest at an individual-level in the study.

## Results

### Overview of methods

We start by depicting the main intuition underlying our hypothesis and model (Figure 1). For simplicity, consider two tissues of interest and assume that gene A is only expressed in tissue 1 whereas gene B is only expressed in tissue 2. The main hypothesis underlying our model is that the genetic susceptibility of a complex trait for a given individual is mediated through regulation of either gene A in tissue 1 or gene B in tissue 2 or both. Having gene expression measurements in every individual at both genes in both tissues can be used to test this hypothesis. Unfortunately, gene expression measurements in large sample sizes such as the UK biobank are not typically available. To circumvent this, since eQTLs explain a substantial heritability of gene expression [17,34], we use eQTLs for each gene in the corresponding tissue as a proxy for the genetically regulated component of the expression. In details, eGST takes as input the phenotype values and the genotype values at a set of variants known to be eQTLs for tissue-specific genes (Figure 1). We consider a set of genes overexpressed in a tissue as the set of tissue-specific genes [20,23]. Formally, the phenotype for individual *i* under the tissue of interest *k* is modeled as 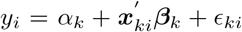, where *α_k_* is the baseline tissue-specific trait mean, *x_ki_* is the vector of normalized genotype values of individual *i* at the eQTL SNPs specific to tissue *k, β_k_* are their effects on the trait, and *ϵ_ki_* is a noise term, *i* = 1,…, *n* and *k* = 1,…, *K*. For simplicity of exposition, we introduce indicator variables for each individual *C_i_* = *k* iff the individual *i* has its tissue of interest *k*. Thus, P(*C_i_* = *k*) = *w_k_* is the prior proportion of individuals for whom the phenotype has *k^th^* tissue-specific genetic effect. We assume that the eQTL SNP sets across *k* tissues are non-overlapping and that each element in *β_k_*, the genetic effect of *k^th^* tissue-specific eQTLs (i.e., eQTLs for tissue-specific genes) on the trait, is drawn from 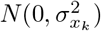. If 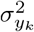 is the variance of the trait under *C_i_* = *k*, then 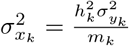, where 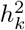 is the heritability of the trait under *C_i_* = *k* due to *k^th^* tissue-specific *m_k_* eQTLs, and is termed as *k^th^* tissue-specific subtype heritability. Under this mixture model, the likelihood of individual *i* takes the form: 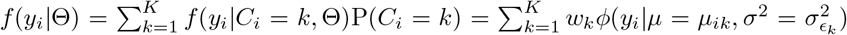, where *ϕ*(.|.) denotes the normal density, 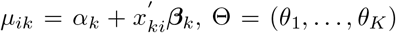 with *θ_k_* denoting *k^th^* tissue-specific set of model parameters. We propose a Bayesian inference approach based on a maximum a posteriori (MAP) expectation maximization (EM) algorithm to estimate the posterior probability that the phenotype of individual *i* is mediated through the genetic effects of eQTLs specific to tissue *k* (P(*C_i_* = *k|X,Y*)). Our main inference is based on this posterior probability. For example, while P(*C_i_* = 1|*X, Y*) > 65% indicates that tissue 1 is likely to be the tissue of interest for individual *i*, a value of the posterior probability around 50% indicates that the genetic susceptibility for the individual does not mediate through a specific tissue.

**Figure 1:**
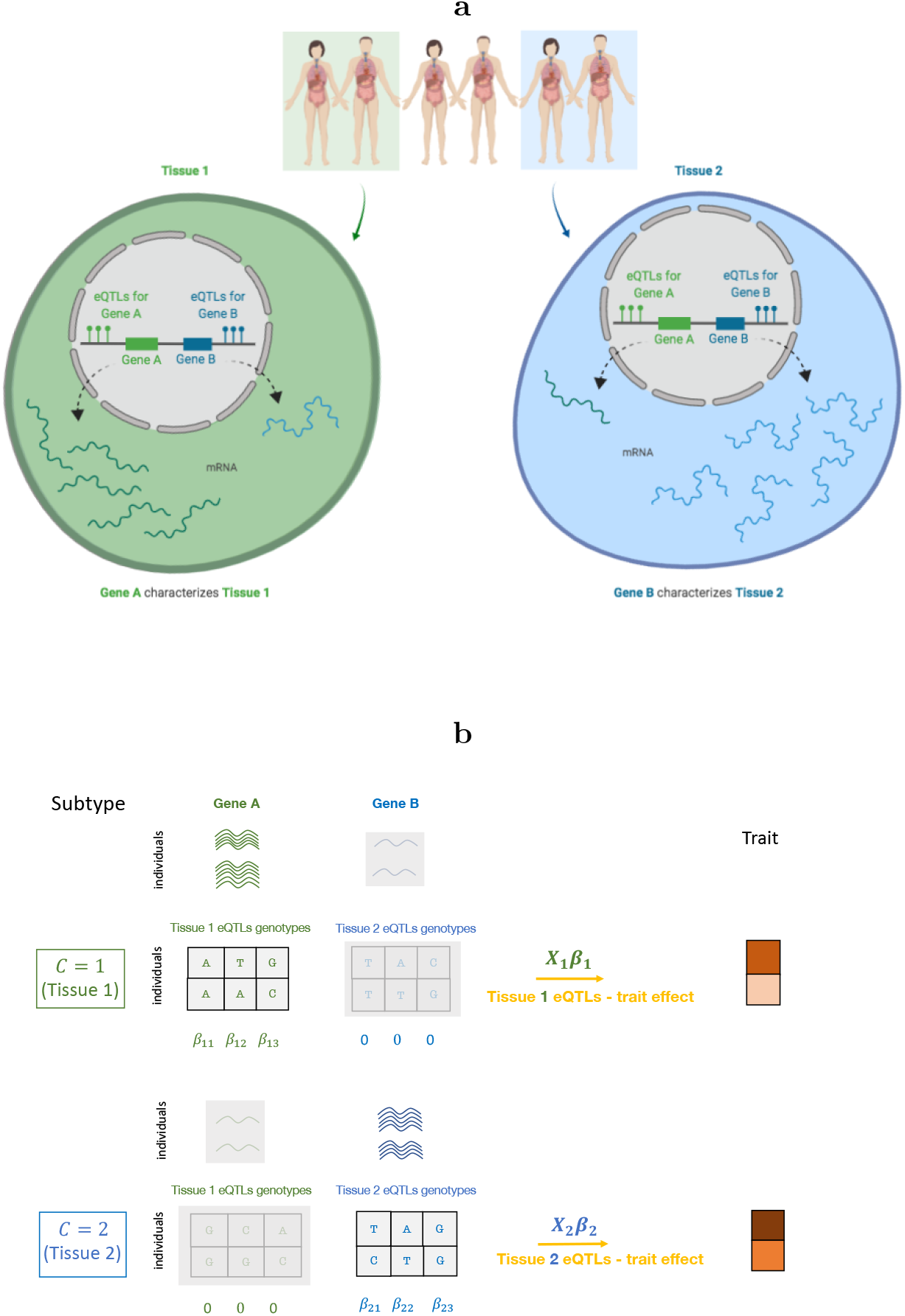
The top diagram (a) explains our main hypothesis and the bottom diagram (b) explains our model, (a): Consider two tissues of interest for a phenotype, tissue 1 and tissue 2, where gene A has higher expression but gene B has much lower expression in tissue 1, and gene B has higher expression but gene A has much lower expression in tissue 2. The key hypothesis is that the susceptibility of the phenotype for the first two individuals is mediated through the effect of gene A in tissue 1, in which case we can assign tissue 1 as the tissue of interest for these individuals (similarly tissue 2 for last two individuals). We refer to the phenotype of the first two individuals as tissue 1 specific subtype and the phenotype of last two individuals as tissue 2 specific subtype. However, two individuals in the middle are not assigned as any of the two tissue-specific subtypes and remain unclassified, (b): We use genotypes at the tissue-specific eQTLs (i.e., eQTLs for tissue-specific genes) as a proxy for the expressions of the corresponding tissue-specific genes. We consider a finite mixture model with each of its components being a linear model regressing the trait on the genotypes at each set of tissue-specific eQTLs. Our method takes as input the individual-level measurements of the phenotype and genotypes at the sets of tissue-specific eQTLs and provides per-individual tissue-specific posterior probabilities as the main output.

### Simulations

We performed simulations to assess the performance of eGST with respect to the accuracy of classifying the tissue of interest across individuals under various scenarios. We simulated phenotypes using the real genotype data from the UK Biobank, in which two tissue-specific eQTL effects generate the phenotype (see Methods). We evaluated the classification accuracy of eGST with respect to the variance explained in the trait specific to the two tissues by the two sets of tissue-specific SNPs’ effects. As expected, the average area under the curve (AUC) increases with the tissue-specific subtype heritability, ranging from AUC of 50% when 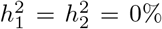 to 95% when 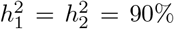 (Figure 2). This is likely due to larger tissue-specific subtype heritabilities inducing better differentiation between the tissue-specific genetic effects. Next, we assessed the performance of eGST compared to a variation of our approach that assumes the parameters of the model to be known. The performance obtained by this gold-standard strategy can be viewed as the maximum achievable under our proposed framework. We find that eGST loses 1.4% — 3.9% AUC on average compared to this strategy across all simulation scenarios considered (Figure 2, Figure S1, S2). We also considered a thresholding scheme on the tissue-specific posterior probabilities to balance total discoveries versus accuracy. As expected, the true discovery rate (TDR) of classifying the tissue of interest increases (hence FDR = 1-TDR decreases) with the posterior probability threshold but the proportion of discovery decreases (Figure 3).

**Figure 2:**
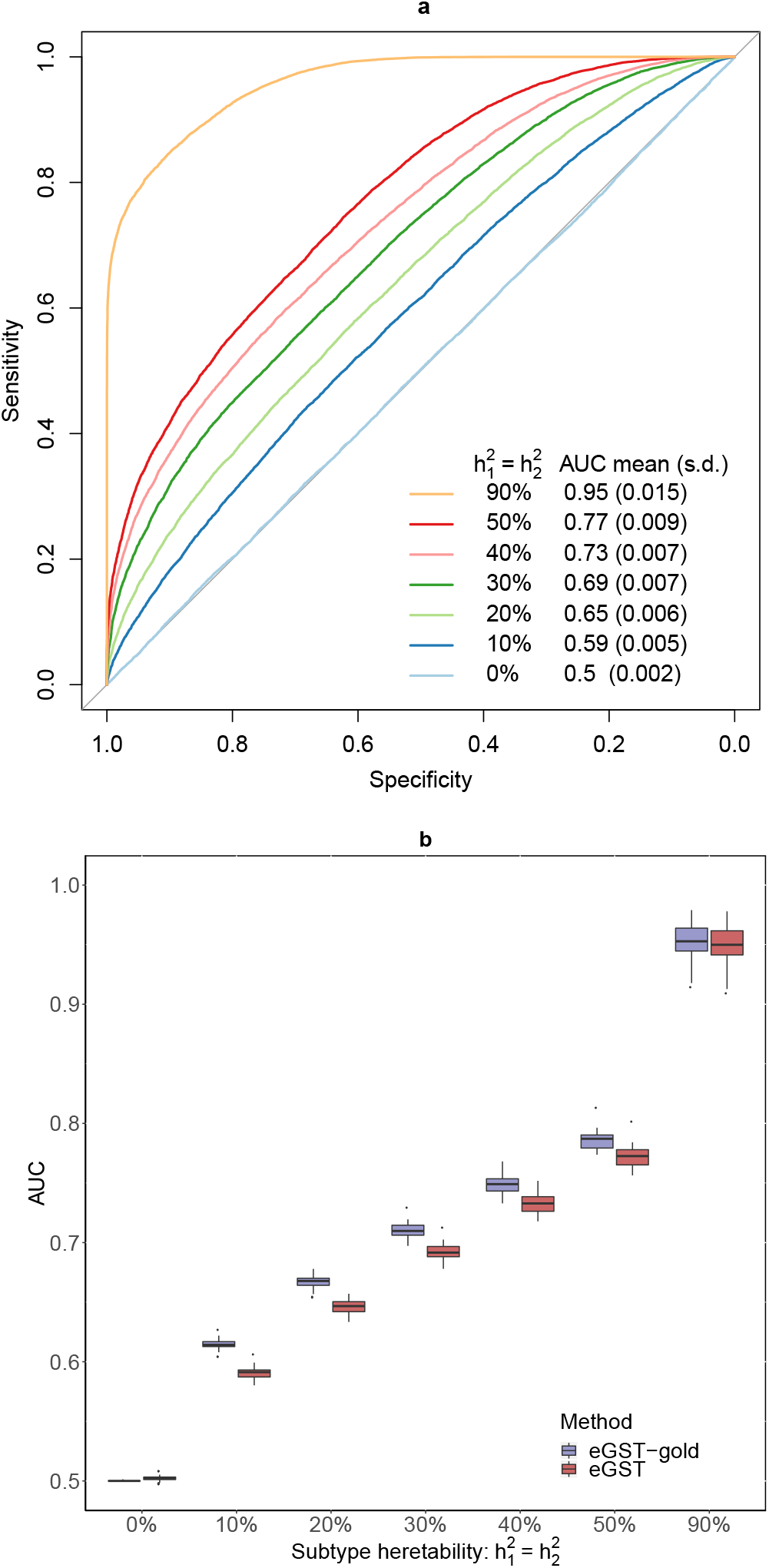
In the first diagram (a), we present the receiver operating characteristic (ROC) curve evaluating the classification accuracy of eGST for a single dataset simulated under the following scenarios: 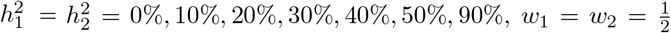, *m*_1_ = *m*_2_ = 1000, *n* = 40000. The mean (across 50 simulated datasets) area under the curve (AUC) obtained by eGST under the same scenarios are also provided. Here 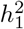 and 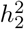 are the heritability of tissue-specific subtypes of the trait due to *m*_1_ and *m*_2_ SNPs representing two sets of tissue-specific eQTL SNPs, *w*_1_ and *w*_2_ are the proportions of individuals in the sample assigned to the two tissues, *n* is the total number of individuals. In the second diagram (b), box plots of AUCs obtained by eGST and the gold-standard strategy implementing our model in which true model parameters were assumed to be known while estimating the tissue-specific posterior probabilities are presented across the same simulation scenarios.

**Figure 3:**
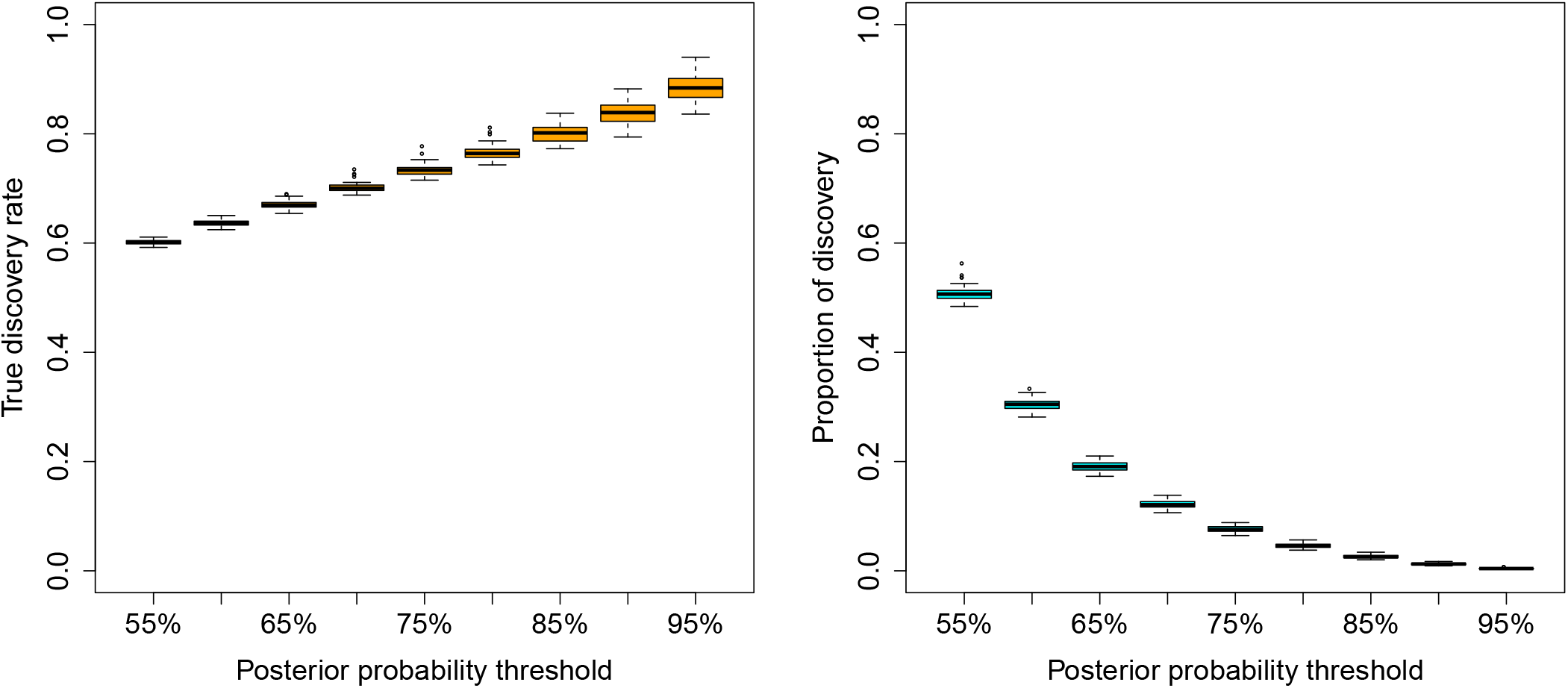
True discovery rate (TDR) and the proportion of discovery (POD) by eGST while classifying the tissue of interest across individuals at increasing thresholds of tissue-specific subtype posterior probability: 55%, 60%, 65%,…, 90%, 95%. Box plots of TDR and POD across 50 datasets simulated under 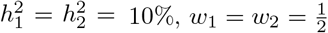, *m*_1_ = *m*_2_ = 1000, *n* = 40,000 are presented. Here 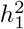 and 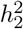 are the heritabilities of the tissue-specific subtypes of the trait due to *m*_1_ and *m*_2_ SNPs representing two sets of tissue-specific eQTL SNPs, *w*_1_ and *w*_2_ are the proportions of individuals in the sample assigned to the two tissues, n is the total number of individuals.

We then explored the effect of other parameters on the classification accuracy. First, we found that increasing sample size *n* from 40, 000 to 100, 000 marginally increases the AUC by an average of 1% across different simulation scenarios (Table S2), which indicates that increasing sample size improves the overall classification accuracy. Second, we observed that as the number of causal SNPs explaining a fixed heritability of each subtype increases, the average AUC marginally decreases. For example, for a fixed subtype heritability explained, the average AUC for 2000 causal SNPs (1000 per tissue) is 1% higher than that for 3000 causal SNPs (1500 per tissue) across different choices of other simulation parameters (Table S3). Third, as the difference between the baseline tissue-specific mean of the trait across tissues increases, the classification accuracy also increases. For example, we find that the AUC increased from 60% (for no difference in tissue-specific phenotype means, *α*_1_ = *α*_2_ = 0) to 63% when *α*_1_ = 0, *α*_2_ = 1 (Table S4). We also explored the impact of the difference between the mean of tissue-specific genetic effect size distributions and observed that the classification accuracy improves compared to zero mean of both causal effects. For example, the AUC increases from 60% to 63% if we consider *E*(*β*_1*j*_) = −0.02 and *E*(*β*_2*j*_) = 0.02, *j* = 1,…, 1000, instead of zero means of *β*_1_, *β*_2_ (Table S5).

Finally, we explored the comparative performance of the MAP-EM algorithm under Bayesian framework (Algorithm 1) and the EM algorithm under frequentist framework (Algorithm 2). Although both approaches yield similar AUC, MAP-EM performed better than EM with respect to the true discovery rate (TDR) at different posterior probability thresholds in nearly all of the simulation scenarios (Figure 4, S3, S4). MAP-EM offered an average of 0.05% — 20% higher TDR (hence lower FDR) than EM across various posterior probability thresholds (Figure 4, S3, S4).

**Figure 4:**
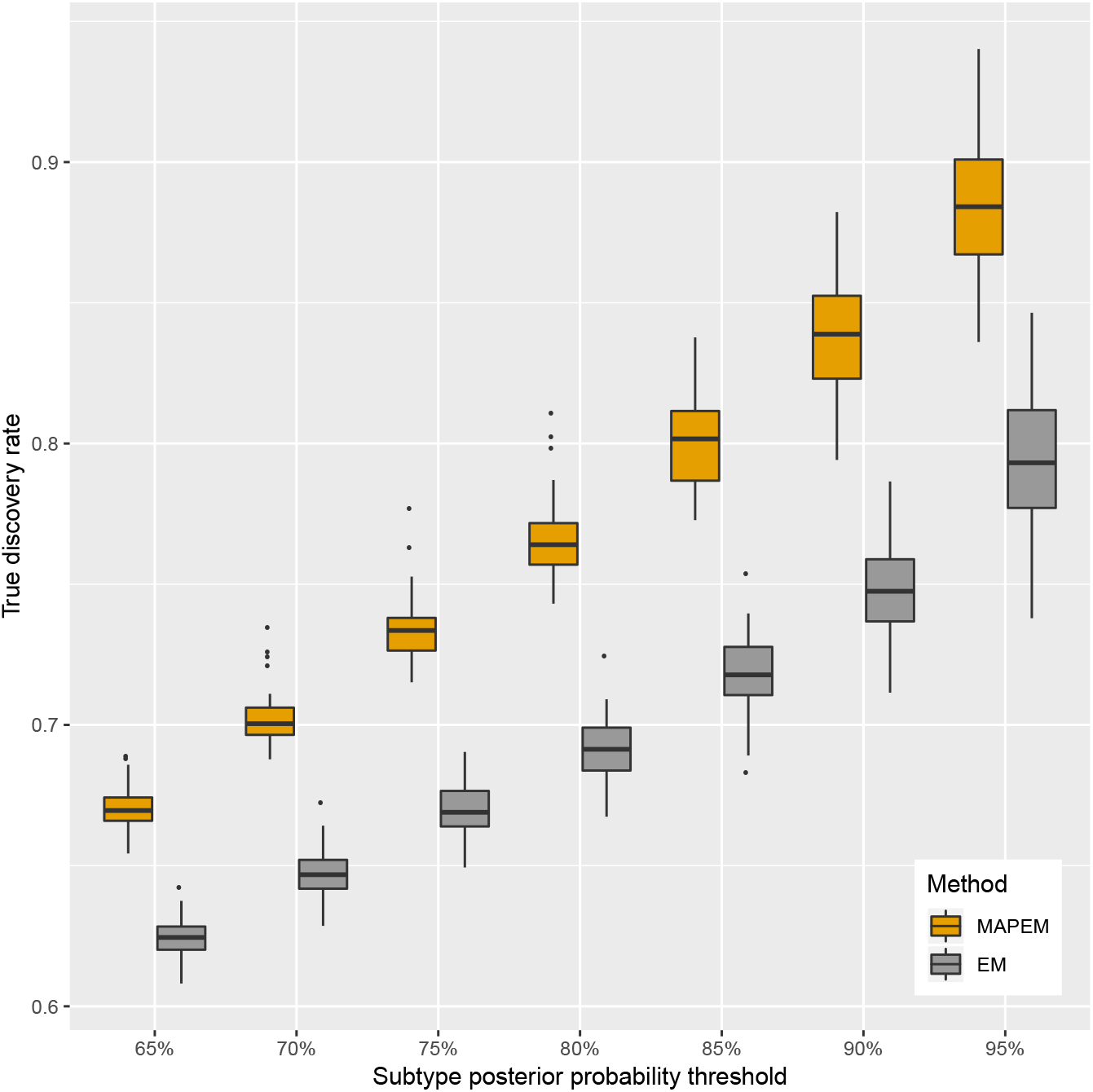
Comparison between the true discovery rate (TDR) of classifying tissue-specific subtypes by the MAP-EM algorithm (under the Bayesian framework of the mixture model which eGST employs) versus the EM algorithm (under the frequentist framework of the mixture model) based on the threshold of tissue-specific subtype posterior probability as 65%, 70%, 75%, 80%, 85%, 90%, 95%, respectively. Box plots of TDR across 50 datasets simulated under 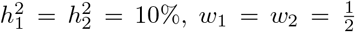, *m*_1_ = *m*_2_ = 1000, *n* = 40,000 are presented. Here 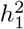 and 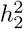 are the heritability of tissue-specific subtypes of the trait due to *m*_1_ and *m*_2_ SNPs representing two sets of tissue-specific eQTL SNPs, *w*_1_ and *w*_2_ are the proportions of individuals in the sample assigned to the two tissues, *n* is the total number of individuals.

### Inferring individual-level tissue of interest for BMI and WHRadjBMI

Having established in simulations that our approach is effective in correctly classifying the individual-level tissue of interest, we next analyzed BMI and WHRadjBMI, two phenotypes that are known to have multiple tissues of interest mediating their genetic susceptibility [21,23,26,27,35–37]. For BMI analysis, we used phenotype and genotype data for 336, 106 individuals in the UK Biobank [32, 33] at 1705 adipose specific eQTLs (i.e., eQTLs for adipose-specific genes) and 1478 brain specific eQTLs (see Methods). We considered rank-based inverse normal transformation of the BMI residual obtained after adjusting BMI for age, sex, and 20 PCs of genetic ancestry to adjust for population stratification. Each SNP eQTL is the top cis-association of a tissue-specific expressed gene [23] in the corresponding tissue [16,18] (see Methods). At 65% threshold of tissue-specific subtype posterior probability, 7.5% of all the individuals where assigned a tissue (adipose or brain) and the rest of the individuals remained unclassified; eGST classified the genetic susceptibility on the BMI of 11, 838 individuals through adipose eQTLs and for 13, 354 individuals through brain eQTLs (Table S1). Individuals classified to each of the tissues are distributed across different bins of BMI (Table S6). While the individuals classified into adipose have a higher mean of BMI (30.4) than the population, the BMI mean for brain-specific individuals (27.6) is very close to the population mean (27.4) (Table S8).

For WHRadjBMI, we included 953 adipose subcutaneous (abbreviated and referred as AS in the following) tissue-specific eQTLs (i.e., eQTLs for AS-specific genes) and 1052 muscle skeletal connective (abbreviated as MS) tissue-specific eQTLs; and inverse normal transformed WHRadjBMI residual (adjusting WHRadjBMI for age, sex, and top 20 PCs) for 336, 018 individuals. Similarly to the BMI analysis, the tissue of interest for WHRadjBMI of a small percentage (5.7%) of all individuals were classified (Table S1), and the remaining individuals were unclassified. Individuals assigned to the two tissues are both spread across different bins of WHRadjBMI (Table S7). The individuals classified into MS have a higher mean of WHRadjBMI and WHR (0.03 and 0.91) than the population, and the mean for AS-specific individuals (−0.04 and 0.85) is lower than the population mean (0 and 0.87) (Table S9).

We permuted the phenotype data across individuals while keeping the eQTL assignment to tissues fixed as it is in the original data (see Methods). For BMI, the average number of individuals classified as a tissue-specific subtype (based on 65% threshold of subtype posterior probability) across 500 random permutations of the phenotype was 7404 (s.d. 700) which is substantially smaller than 25,192 individuals classified as real adipose or brain specific subtype of BMI in the original data. For WHRadjBMI, the average number of individuals classified as a tissue-specific subtype across 500 random permutations of the phenotype was 3433 (s.d. 517) compared to 19,041 individuals classified as real AS and MS specific subtypes of WHRadjBMI. To estimate the tissue-specific subtype heritability, we employed the MCMC algorithm that implements eGST under the Bayesian framework (Algorithm S1). Through simulations we found that the MCMC estimates the tissue-specific subtype heritability more unbiasedly than the MAP-EM algorithm (results not provided for brevity), but is computationally much slower. In BMI analysis, the posterior mean of the tissue-specific subtype heritability was found to be 3.6% (posterior s.d. 0.2%) and 3.7% (posterior s.d. 0.2%) for brain and adipose, respectively. Similarly, in WHRadjBMI analysis, the posterior mean of the tissue-specific subtype heritability was 3.1% (posterior s.d. 0.2%) and 2.5% (posterior s.d. 0.2%) for adipose and muscle, respectively.

#### Genetic characteristics

To confirm that eGST identified groups of individuals with different genetic basis, we contrasted the SNP effects of the adipose and brain-specific eQTLs on the BMI in those individuals assigned to the adipose (or the brain specific subtype). As expected, we find that in the individuals classified as having the brain-specific subtype in their genetic contribution to BMI, the magnitude of the effect size of a brain eQTL SNP is larger than the corresponding effect size magnitude of an adipose eQTL SNP (Wilcoxon rank sum (WRS) right tail test p-value < 2.2 × 10^−16^ (Table 1)). The opposite is true for the individuals classified as having the adipose-specific subtype of BMI. We also find that the magnitude of effect size of the adipose eQTLs are larger in the adipose-specific individuals than that in the brain-specific individuals, and we find statistical evidence supporting the analogous hypothesis about the brain eQTLs, brain-specific, and adipose-specific individuals of BMI (Table 1). We observe the same pattern in our analogous analysis for WHRadjBMI (Table 1).

**Table 1:**
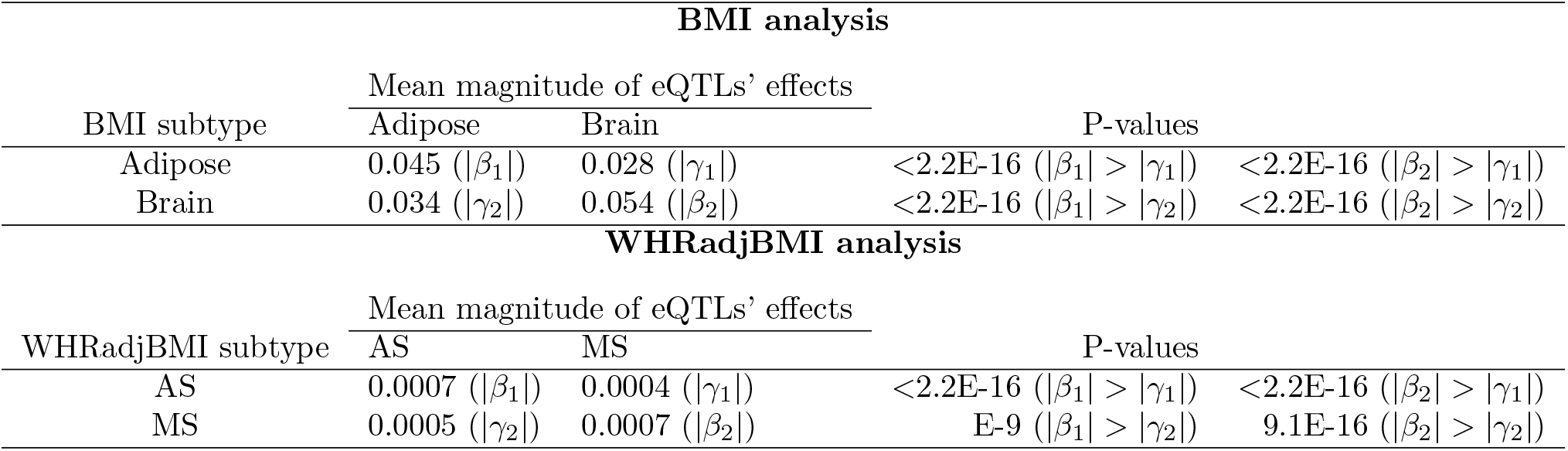
Genetic heterogeneity between groups of individuals assigned to tissue-specific subtypes of BMI (or WHRadjBMI). In the BMI analysis, *β*_1_ denotes the SNP-effect of an adipose eQTL on BMI in those individuals assigned to the adipose-specific subtype of BMI, *β*_2_ is the SNP-effect of a brain eQTL on the brain-specific subtype of BMI, *γ*_1_ is the SNP-effect of a brain eQTL on the adipose subtype and *γ*_2_ is the effect of an adipose eQTL on the brain subtype. We provide the mean magnitude of the effect sizes of a tissue-specific eQTLs on the BMI of the corresponding tissue-specific group of individuals (e.g., joint SNP-effect of the adipose eQTLs on the BMI of individuals with adipose subtype), and the p-values obtained from the Wilcoxon rank sum (WRS) right tail tests of effect heterogeneity. For each test, the alternative hypotheses are listed in parentheses, while the null hypothesis is the equality between the corresponding pair of parameters. These parameters are defined in the same way for the adipose subcutaneous (AS) and muscle skeletal (MS) tissue-specific subtype of WHRadjBMI and the same analyses are performed.

#### Phenotypic characteristics of individuals with a prioritized tissue

Next, we explored the phenotypic characteristics of the individuals assigned with a prioritized tissue. We considered 106 phenotypes in the UK Biobank and tested each one for being differentially distributed (heterogeneous) between the individuals of each tissue-specific subtype and the remaining population (see Methods). In aggregate for BMI, 45 quantitative traits and 40 qualitative traits (total 85 among 106) were significantly heterogeneous between at least one of the BMI-adipose or BMI-brain specific groups versus the remaining population (Table S8, S10, Table 2). None of these 106 traits was found to be differentially distributed between a random set of individuals from the population (with the same size as a tissue-specific subtype group) and the remaining population (see Methods). 33 quantitative and 34 categorical traits showed heterogeneity in both the adipose group versus the population, and the brain group versus population. We found 6 quantitative and 3 categorical traits heterogeneous for individuals in the adipose group but not the brain group, and found 6 quantitative and 3 categorical traits heterogeneous for individuals in the brain group but not the adipose group (Table S8, S10, Table 2).

**Table 2:**
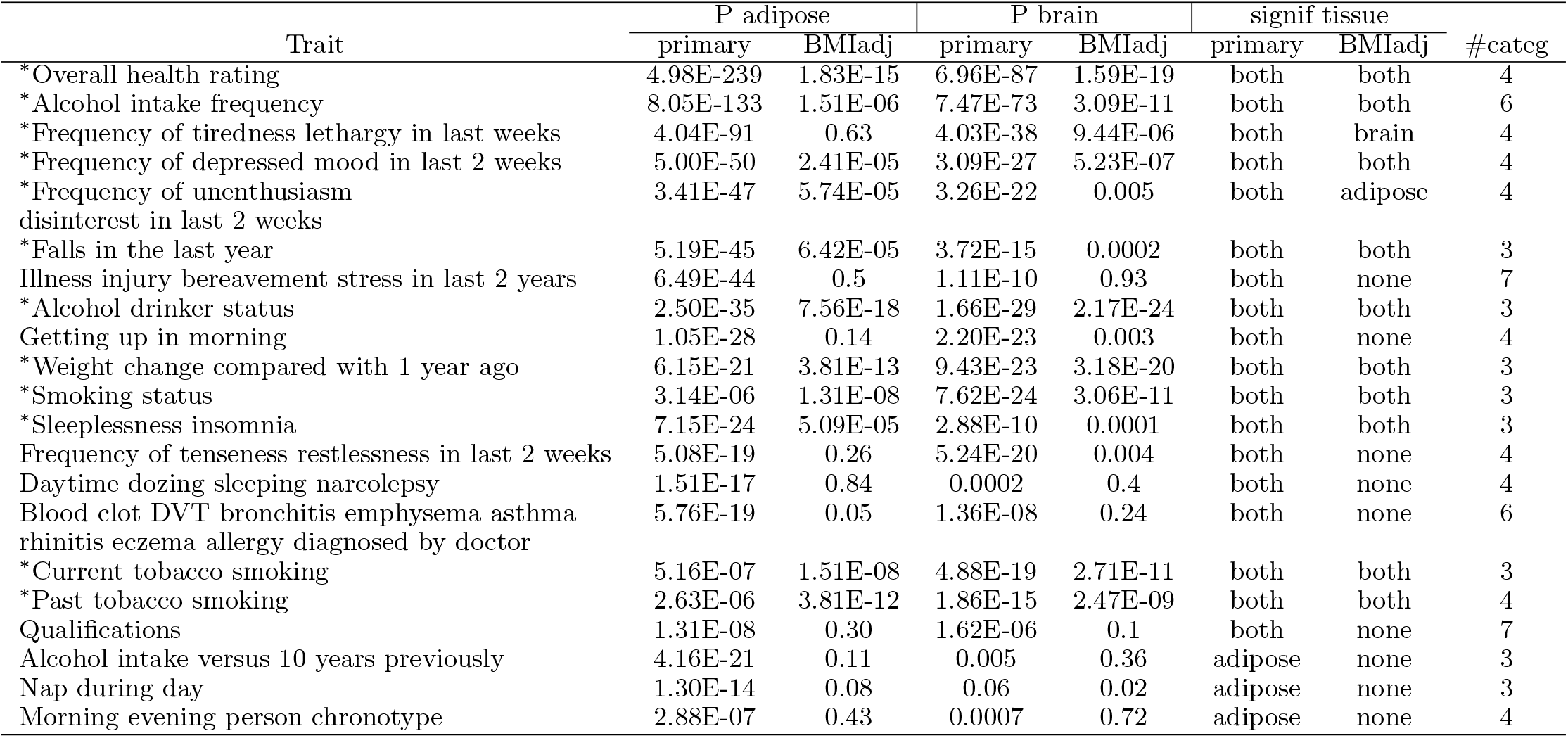
Qualitative/categorical traits with three or more categories that are differentially distributed between at least one of the adipose and brain-specific subtype groups of individuals for BMI and the remaining population. For each trait, we provide the p-values of testing heterogeneity between each tissue-specific subtype group of individuals and the remaining population before (primary) and after BMI adjustment (BMIadj). For each trait, tissue-specific groups which appear to be significantly heterogeneous (signif tissue) before (primary) and after BMI adjustment (BMIadj) are also provided. The asterisk mark attached to the traits indicate which trait remains differentially distributed between at least one of the tissue-specific groups and the remaining population after BMI adjustment. The number of categories for each trait (#categ) are also listed.

| Trait | P adipose |  | P brain |  | signif tissue |  | #categ |
| --- | --- | --- | --- | --- | --- | --- | --- |
|  | primary | BMIadj | primary | BMIadj | primary | BMIadj |  |
| *Overall health rating | 4.98E-239 | 1.83E-15 | 6.96E-87 | 1.59E-19 | both | both | 4 |
| *Alcohol intake frequency | 8.05E-133 | 1.51E-06 | 7.47E-73 | 3.09E-11 | both | both | 6 |
| *Frequency of tiredness lethargy in last weeks | 4.04E-91 | 0.63 | 4.03E-38 | 9.44E-06 | both | brain | 4 |
| *Frequency of depressed mood in last 2 weeks | 5.00E-50 | 2.41E-05 | 3.09E-27 | 5.23E-07 | both | both | 4 |
| *Frequency of unenthusiasm<br>disinterest in last 2 weeks | 3.41E-47 | 5.74E-05 | 3.26E-22 | 0.005 | both | adipose | 4 |
| *Falls in the last year | 5.19E-45 | 6.42E-05 | 3.72E-15 | 0.0002 | both | both | 3 |
| Illness injury bereavement stress in last 2 years | 6.49E-44 | 0.5 | 1.11E-10 | 0.93 | both | none | 7 |
| *Alcohol drinker status | 2.50E-35 | 7.56E-18 | 1.66E-29 | 2.17E-24 | both | both | 3 |
| Getting up in morning | 1.05E-28 | 0.14 | 2.20E-23 | 0.003 | both | none | 4 |
| *Weight change compared with 1 year ago | 6.15E-21 | 3.81E-13 | 9.43E-23 | 3.18E-20 | both | both | 3 |
| *Smoking status | 3.14E-06 | 1.31E-08 | 7.62E-24 | 3.06E-11 | both | both | 3 |
| *Sleeplessness insomnia | 7.15E-24 | 5.09E-05 | 2.88E-10 | 0.0001 | both | both | 3 |
| Frequency of tenseness restlessness in last 2 weeks | 5.08E-19 | 0.26 | 5.24E-20 | 0.004 | both | none | 4 |
| Daytime dozing sleeping narcolepsy | 1.51E-17 | 0.84 | 0.0002 | 0.4 | both | none | 4 |
| Blood clot DVT bronchitis emphysema asthma<br>rhinitis eczema allergy diagnosed by doctor | 5.76E-19 | 0.05 | 1.36E-08 | 0.24 | both | none | 6 |
| *Current tobacco smoking | 5.16E-07 | 1.51E-08 | 4.88E-19 | 2.71E-11 | both | both | 3 |
| *Past tobacco smoking | 2.63E-06 | 3.81E-12 | 1.86E-15 | 2.47E-09 | both | both | 4 |
| Qualifications | 1.31E-08 | 0.30 | 1.62E-06 | 0.1 | both | none | 7 |
| Alcohol intake versus 10 years previously | 4.16E-21 | 0.11 | 0.005 | 0.36 | adipose | none | 3 |
| Nap during day | 1.30E-14 | 0.08 | 0.06 | 0.02 | adipose | none | 3 |
| Morning evening person chronotype | 2.88E-07 | 0.43 | 0.0007 | 0.72 | adipose | none | 4 |

For example, *hemoglobin concentration* and *snoring* were heterogeneous for both the adipose and brain groups, *lymphocyte count* and *alcohol intake versus 10 years previously* were heterogeneous only for the adipose group, and *birth weight, nervous feelings* only for the brain group (Table S8, S10, Table 2). We observe that *hemoglobin concentration* was lower in individuals from both groups when compared to the population, whereas *reticulocyte percentage* was relatively higher in individuals of the adipose but lower in those with the brain tissue compared to the population (Figure 5, Table S8). Among binary traits, *snoring* was more prevalent in those from the adipose group and less prevalent in brain group compared to the population (Figure 6). We observe that for most of the case-control traits, both the tissue-specific groups of individuals had a higher risk of developing the disease compared to the population (Figure 6). Of note, when the tissue-specific relative change of the traits (see Methods) were in the same direction across tissues, they were of different magnitude for a majority of the traits (Figure 5, 6). For example, the relative change were 15% and 8% for *neutrophil count* (Table S13). Similar to BMI, we observed phenotypic heterogeneity across individuals with AS (MS) as the prioritized tissue for WHRadjBMI (Figure S5, S6, Table S9, S11, S12, Supplementary Note).

**Figure 5:**
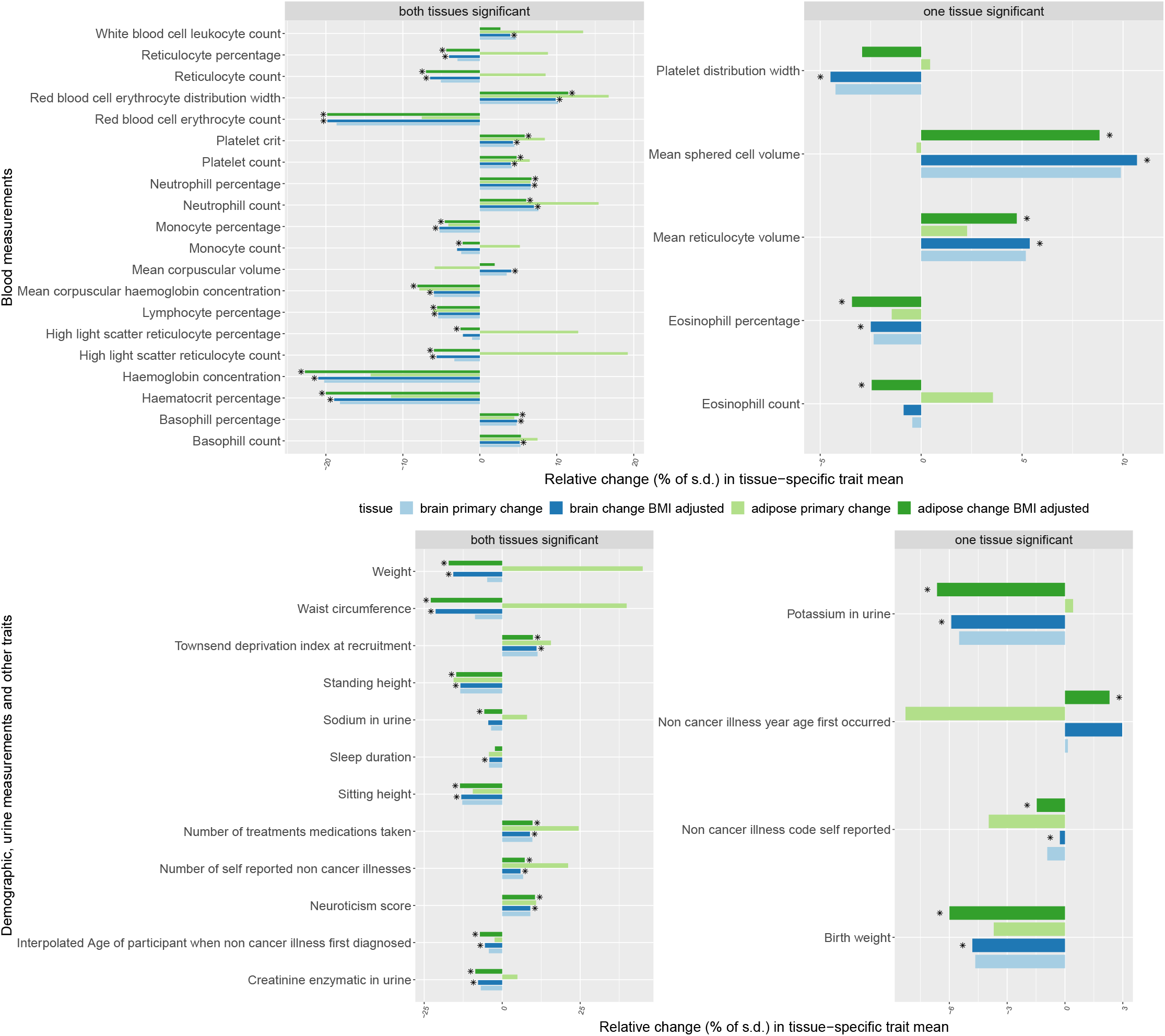
Percentage of tissue-specific relative change of the quantitative traits that were differentially distributed between the individuals assigned to a tissue-specific subtype of BMI and the remaining population. Traits in the left panels are primarily heterogeneous for both tissue-specific groups and traits in the right panels are heterogenous for one tissue-specific group. We measure the tissue-specific relative change of a trait by: 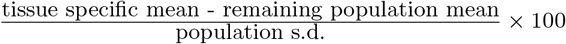, where the tissue-specific mean is computed only in the in-dividuals with the corresponding tissue-specific subtype. The same measure is calculated for a trait residual obtained after adjusting for BMI to quantify the tissue-specific relative change of the trait after BMI adjustment. The faded green (or blue) bar presents primary adipose (or brain) tissue-specific relative change of a trait compared to the remaining population. The dark green (or blue) bar presents the BMI-adjusted adipose (or brain) specific relative change of a trait. Each trait listed here was found to be differentially distributed between at least one of the adipose or brain specific groups and the remaining population after BMI adjustment. For each trait, the asterisk mark attached to the bars indicates which tissue-specific group remains significantly heterogeneous after BMI adjustment.

**Figure 6:**
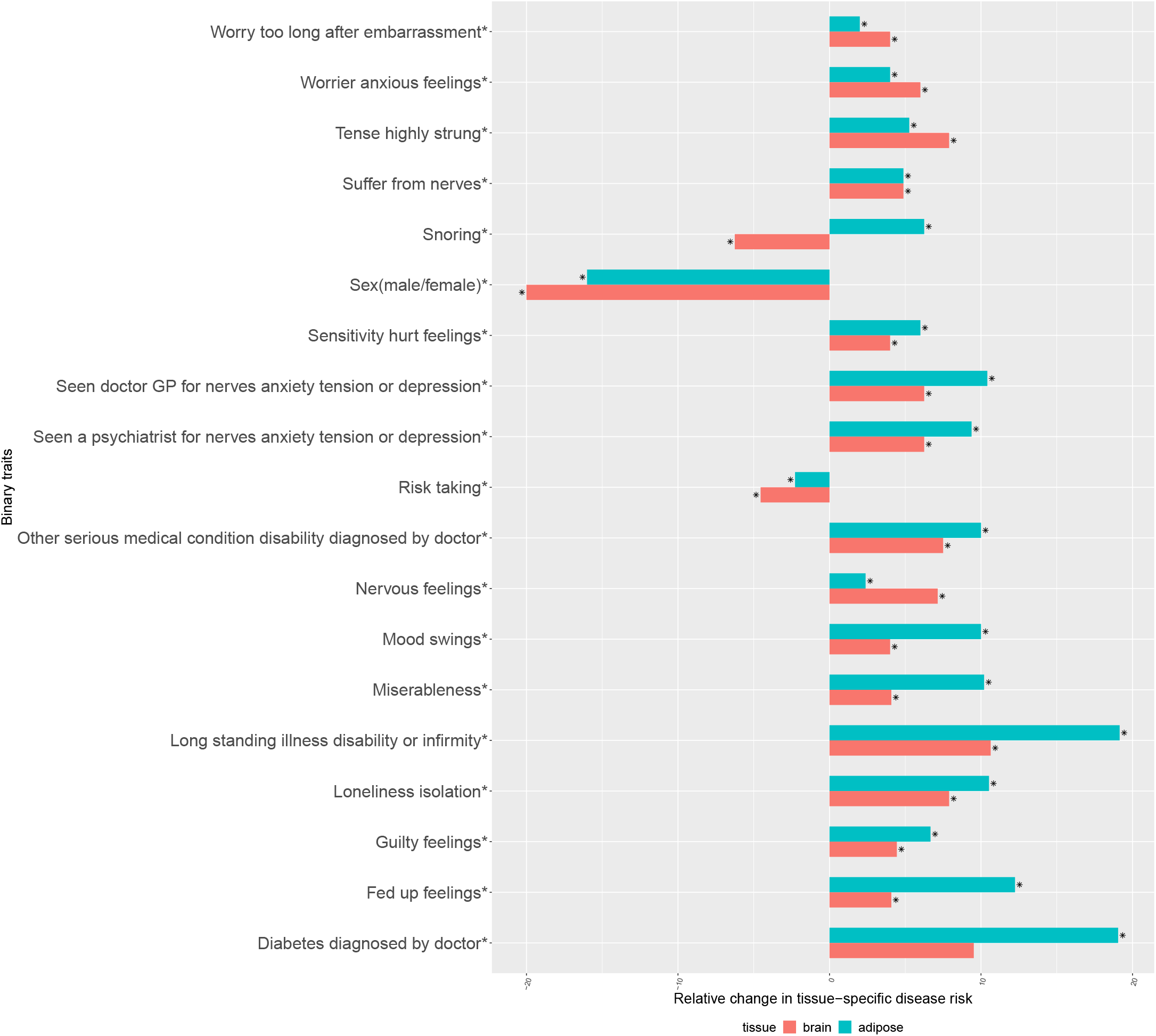
Percentage of tissue-specific relative change in the risk of case-control traits between the individuals assigned to a tissue-specific subtype of BMI and the population. The tissue-specific relative change of a disease risk is measured by: 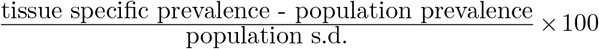. Tissue-specific prevalence of the disorder was computed only in the individuals classified as the corresponding tissue-specific subtype of BMI. The asterisk mark attached to the traits indicate which trait remains differentially distributed between at least one of the adipose and brain tissue-specific groups of individuals and the remaining population after BMI adjustment. For each trait the asterisk mark attached to the bars indicate which tissue-specific group of individuals remains significantly heterogeneous for the trait after BMI adjustment.

Since BMI itself was differentially distributed between the individuals of the adipose subtype as well as brain subtype compared to the remaining population, we investigated whether the heterogeneity of 84 non-BMI traits (Table S8 and S10, Table 2) were induced due to BMI heterogeneity (see Methods). After BMI adjustment, 72 (out of 85) traits remained heterogeneous (41 quantitative traits [Figure 5, Table S13] and 31 qualitative traits [Figure 6, Table 2]) consistent with unique phenotypic characteristics of these individuals beyond the main phenotype effect. All the quantitative traits which remained heterogeneous after BMI adjustment have the same direction in BMI-adjusted tissue-specific relative change (see Methods) in both adipose and brain compared to the population (Figure 5, Table S13). Since we used linear regression while evaluating BMI-adjusted tissue-specific relative change of heterogeneous non-BMI quantitative traits, we also investigated a model-free BMI random matching strategy. We assessed the magnitude of relative change of a trait between individuals with the brain (or adipose) subtype and a group of BMI-matched random individuals drawn from the population (see Methods). For example, the magnitude of primary brain-specific relative change (prior to BMI matching) for *hemoglobin concentration (mean reticulocyte volume)* decreased from 20% (5%) to 4% (2%) after BMI matching (Table S15). Of note, it is very difficult to exactly match BMI between a tissue-specific subtype group and the corresponding random group of individuals, because bins of BMI in the tail of its distribution contain very few individuals (Table S6), the majority of whom were assigned to a tissue-specific subtype. We observed the same pattern in the results from analogous analyses for WHRadjBMI (Table S14, S16, Supplementary Note).

To better understand the phenotypic characteristics of the individuals classified to a specific tissue, we performed the following two experiments. First, we shuffled the tissue-specific eQTL SNPs between tissues to create an artificial tissue-specific eQTL set and implemented eGST to identify groups of individuals having subtype specific to the artificial tissues (see Methods). We found that the mean of artificial tissue-specific means of a quantitative trait (found primarily heterogeneous between adipose and/or brain specific group versus the remaining population [Table S8]) across eQTL shuffles was significantly further from the original corresponding tissue-specific trait mean (Table S17 for BMI and Table S18 for WHRadjBMI). For example, for *waist circumference,* the mean of pseudo tissue-specific means over the random eQTL shuffles is 93.25 for adipose and 91.36 for brain, which are significantly different from the original adipose-specific mean 95.5 (P < 10^−100^) and brain-specific mean 89.2 (P < 10^−100^), respectively (Table S17). The same pattern was observed for the primary phenotypes BMI (Table S17) and WHRadjBMI (S18) themselves. Second, we permuted the phenotype data across individuals while keeping the eQTL assignment to tissues fixed as it is in the original data. As before, we also observed that the mean of tissue-specific means of a quantitative trait (found primarily heterogeneous between a tissue-specific group versus the remaining population [Table S8]) across random phenotype permutations was significantly further from the original corresponding tissue-specific trait mean (Table S19 for BMI and Table S20 for WHRadjBMI).

### Computational efficiency

The MAP-EM algorithm underlying eGST is computationally efficient. 70 MAP-EM iterations in the BMI analysis (336*K* individuals with 1705 adipose-specific eQTLs and 1478 brain-specific eQTLs) took a runtime of 1.75 hours and yielded a log likelihood improvement of 2 × 10^−8^ in the final iteration. Though we ran eGST for a pair of tissues only considering the top eQTL per gene, it is computationally feasible to analyze larger datasets considering more eQTLs and multiple tissues simultaneously.

## Discussion

We proposed a novel approach to quantify tissue-wise genetic contribution to a complex trait and prioritize a relevant tissue for every individual in the study, integrating genotype and phenotype data and an external expression panel data. We applied our method to infer individual-level tissue of interest for BMI and WHRadjBMI in the UK Biobank, integrating expression data in brain, adipose, and muscle tissues from the GTEx consortium, previously shown to be enriched in heritability for these phenotypes [21,23,26,27].

Our approach identified subgroups of individuals with their genetic susceptibility to the trait mediated in a tissue-specific manner. Interestingly, multiple metabolic traits, neuropsychiatric traits, and other traits attained significant differences between the tissue-specific groups of individuals and the remaining population, suggesting a biologically meaningful interpretation for these subgroups of individuals. Even after adjusting the traits for the primary phenotype (BMI or WHRadjBMI), a majority of the traits remained differentially distributed between a tissue-specific group and the remaining population.

We note that for some complex traits multiple (more than two) tissues can be biologically relevant. Although in this work we demonstrated the utility of eGST for a pair of tissues for BMI and WHRadjBMI, the MAP-EM algorithm underlying eGST is general and can be applied to any number (≥ 2) of tissues. We note that our model can alternatively be viewed as an approach to assign individuals’ phenotypes to a collection of tissues that are biologically important for the trait based on tissue-specific polygenic risk score [38]. We did not explicitly model for individuals who have their genetic contribution to the trait mediated through both tissues. We anticipate that such individuals would be assigned posterior probabilities equally distributed across the tissues and hence not appear in the tails of the tissue-specific subtype posterior probability distribution. We developed our model under minimal assumptions on tissue-specific genetics. Following previous studies [20, 23] we characterized a tissue by a set of genes specifically over-expressed in it. However, the best possible strategies of choosing an optimal subset of genes (e.g. combination of both over-expressed and low-expressed genes) to efficiently characterize a tissue need to be further investigated. In real data application, we considered the top eQTL of each gene in a tissue mainly for computational convenience. However, it is straightforward to include more eQTLs in the analysis.

The estimated tissue-specific subtype heritability across tissues appeared to be small (3% – 4%) for BMI and WHRadjBMI. These estimates should increase upon inclusion of more eQTLs in the analyses. However, we observed that the estimates are comparable to the average estimated heritability of six complex traits (3.4%) due to the effects of imputed expression of genes (in blood/adipose tissue) on the trait (provided in [17]). Also, not all of a complex trait’s heritability may mediate through gene expression. As mentioned in the introduction, a recent study [31] has proposed to integrate clinical features related to a disease and imputed gene expression profiles to identify subtypes of the disease. We note that the objective of our study is distinct, and the two approaches are not comparable. Because, we aim to explicitly quantify tissue-wise genetic contribution to the trait at an individual-level and prioritize a relevant tissue for each individual. From a methodological perspective, we proposed an explicitly likelihood-based classification framework in contrast to their multi-view clustering algorithm.

We conclude with several caveats and limitations of our work and opportunities for future improvement. We investigated the utility of eGST using adipose and brain tissues for BMI [21,23,27,35–37], and using adipose and muscle tissues for WHRadjBMI [23,26]. However, the true tissues of interest could be different due to limitations in the existing studies. Since eGST depends on the choice of relevant tissues for a trait, a possible generalization of the model could include an additional mixture component which does not associate to any of the tissues considered (a null component) and represents individuals for whom none of the tissues is relevant. In BMI analysis, a preliminary experimentation with this model indicates that the subgroup of individuals assigned to the null component remained unclassified (to any of the tissues) by the primary 2-component model. That said, we emphasize that eGST is a general analytic framework that can be applied to a collection of tissues for any complex trait. Even though eGST identified subgroups of individuals having their genetic contribution to the trait mediated in a tissue-specific manner, a major proportion of individuals remained unclassified. Few possible reasons are that we considered two tissues in the analysis, but multiple tissues can be relevant for the trait; we considered the top eQTL for each tissue-specific gene. We note that other types of tissue-specific QTLs (e.g., methylation QTLs, histone QTLs, splicing QTLs, etc. [39]) can also be combined with eQTLs to create a set of SNPs that better represent a tissue-specific genetic architecture.

We developed the model for continuous traits, meaning that to extend the method for case-control data, we would need to use a logistic regression likelihood. Another future methodological investigation is to extend the model under penalized regression framework; if the number of SNPs characterizing the genetic architecture of a tissue becomes large and the ratio between the number of individuals and number of SNPs in the data decreases, model fitting issues can arise. Finally, Figure 1b motivates that if gene expression data across tissues are available, it is possible to use the expression data itself to identify expression subtypes of the trait. However, since expression data is not available in most GWAS cohorts, an alternative avenue will be to impute the genetically regulated component of gene expression, e.g., using PrediXcan [34], EpiXcan [40] and identify tissue of interest based on imputed gene expression. While a possible advantage of such approach will be that all cis eQTLs can be unified to impute tissue-specific expression (instead of top few eQTLs only), a significant noise in the predicted expression due to limited sample size of expression panel data can also trim the improvement in performance. Another limitation of our analysis was that we focused on the GTEx data which does not have a large sample size across tissues. In future work, we plan to apply eGST on other complex traits integrating expression datasets of larger sample size. We provide a user-friendly R-software package ‘eGST’ for general use of our approach.

## Methods

### Model

For simplicity, we describe the model assuming two (*K* = 2) tissues of interest. Suppose, for *n* unrelated individuals, we have phenotype data *Y* = (*y*_1_, …,*y_n_*) and expression data for two sets of tissue-specific expressed genes *E*^(1)^, *E*^(2)^ characterizing the two tissues. We define an indicator variable *C* such that for an individual, *C* = *k* iff the genetic susceptibility of the phenotype of the individual is mediated through tissue *k, k* = 1, 2 (Figure 1). We model the phenotype of individual *i* based on the tissue-specific expression of the two sets of tissue-specific genes as:

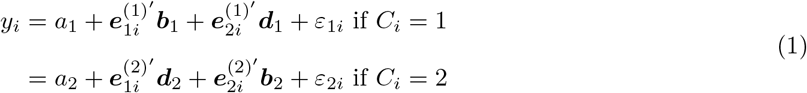

Here, *α*_1_ and *α*_2_ represent the baseline tissue-specific trait means. 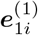 and 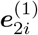 denote the vector of expression values of the first and second tissue-specific set of genes for individual i in the first tissue, respec-tively; 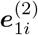 and 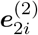 denote the vector of expression values of the first and second tissue-specific set of genes for individual *i* in the second tissue. Under *C_i_* = 1, ***b***_1_ and ***d***_1_ denote the effects of expression of the first and second tissue-specific set of genes in the first tissue on the trait, respectively. Similarly, when *C_i_* = 2, *b*_2_ and *d*_2_ denote the effects of expression of the two gene sets in the second tissue on the trait. If *C_i_* = 1, we assume that the expression of second tissue-specific genes (much low expressed in first tissue) in the first tissue have no effect (***d***_1_ = **0**) on the phenotype of the individual. Similarly, when *C_i_* = 2, we assume that the expression of first tissue-specific genes (much low expressed in second tissue) in the second tissue have no effect (***d***_2_ = **0**) on the phenotype. Thus, we obtain the following model under these assumptions:

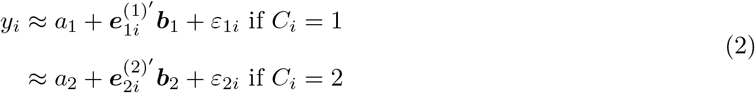

Expression datasets in general have limited sample size and are not available in large GWAS cohorts. Therefore, we consider genetically regulated component of a tissue-specific gene’s expression. However, genetic component of expression predicted by integrating an external panel of expression data [17, 34] can have substantial noise, which is mainly due to limited sample size of expression panel (e.g., GTEx). Since eQTLs explain a substantial heritability of gene expression, we use genotypes of tissue-specific eQTLs (i.e., eQTLs for tissue-specific genes) as a proxy for the genetically regulated component of the expressions of the corresponding tissue-specific genes. Suppose, in a GWAS cohort, we have phenotype data, and genotype data for the two sets of tissue-specific eQTL SNPs corresponding to the two sets of tissue-specific expressed genes, one comprising m1 SNPs and the other comprising m2 SNPs. Then, we consider the following model for the phenotype of individual *i*:

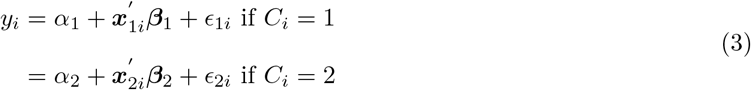

So, the phenotype of individual *i* under the tissue of interest *k* is modeled as 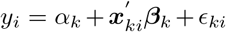, where *α_k_* is the baseline tissue-specific trait mean, *x_ki_* is the vector of normalized genotype values of individual *i* at the eQTL SNPs specific to tissue *k, β_k_* = (*β*_*k*1_, *β*_*k*2_ …, *β_km_k__*) are their effects on the trait under *C_i_* = *k*, and *ϵ_ki_* is a noise term, *i* = 1,…, *n* and *k* =1, 2. The random errors are distributed as: 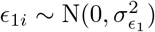 and 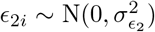. The above mixture model can be viewed as a variant of finite mixture of regression models where each component is a linear model with a distinct set of predictors. Of note, the mixture model in our context is identifiable because the mean parameter in each component is a function of the genotype vector of the set of tissue-specific eQTLs, which is distinct across tissues [41].

#### Prior distributions

P(*C_i_ = k*) = *w_k_* is the prior proportion of individuals for whom the phenotype has *k^th^* tissue-specific genetic effect. We assume that the eQTL SNP sets across *k* tissues are non-overlapping and that each element in *β_k_*, the genetic effect of *k^th^* tissue-specific eQTLs on the trait, is independently drawn from 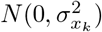. If 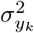 is the variance of the trait under *C_i_ = k*, then 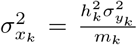, where 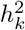 is the heritability of the trait under *C_i_ = k* due to *k^th^* tissue-specific *m_k_* eQTLs, and is termed as *k^th^* tissue-specific subtype heritability. We also assume that 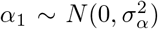 and 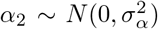, with fixed 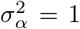. For *K* = 2, we assume that *w*_1_ ~ Beta(*s*_1_,*s*_2_) [*w*_2_ = 1 — *w*_1_], which will be a Dirichlet distribution for more than two tissues. We consider fixed values of *s*_1_ = *s*_2_ = 1. Next, we assume: 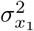 and 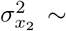 Inverse-Gamma(*α_x_, b_x_*); 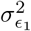 and 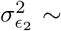 Inverse-Gamma(*α_ϵ_, b_ϵ_*). We choose fixed values of *α_x_, b_x_, α_ϵ_, b_ϵ_* such that in the prior expectation, 5% of the total variance of each tissue-specific subtype (under *C_i_* = 1 or 2) of the trait is explained by the corresponding set of tissue-specific eQTL SNPs and 95% of the variance remains unexplained.

### Inference procedure

Under this Bayesian framework, we implemented the maximum a posteriori (MAP) expectation-maximization (EM) algorithm (Algorithm 1) to estimate the posterior probability that the phenotype of individual i is mediated through the genetic effects of eQTLs specific to tissue *k* (P(*C_i_* = *k*|*X, Y*)). We note that it is also possible to consider a frequentist framework of the mixture model, i.e., instead of having a distribution, 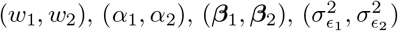 can be assumed to have a fixed unknown true value. We implemented an EM algorithm to estimate the tissue-specific posterior probability across individuals under the frequentist framework. Next, for a general *K* (≥ 2) number of tissues, we outline the MAP-EM algorithm that implements the Bayesian framework, and the EM algorithm that implements the frequentist framework of the mixture model [42,43].

For individual *i* and tissue *k, i* = 1,…, *n* and *k* = 1,…, *K, P*(*C_i_* = *k*) = *w_k_* Σ_*k*_ *w_k_* = 1. Denote *k^th^* tissue-specific set of parameters by 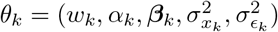 and full set of parameters by Θ = (*θ*_1_,…, *θ_K_*). Under the mixture model, the likelihood of individual *i* takes the following form:

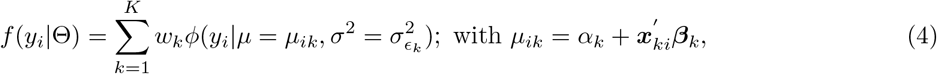

where *ϕ*(.|.) denotes the normal density. Thus, the full data log-likelihood conditioned on Θ is given by: 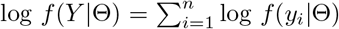. The prior log-likelihood of (*C*_1_ …, *C_n_*) is given by 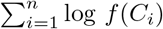, where *f* (*C_i_*) is: *P*(*C_i_* = *k*) = *w_k_*; (*w*_1_,…, *w_K_*) ~ Dirichlet(*s*_1_,…, *s_K_*). The prior of *k^th^* tissue-specific parameters *θ_k_* has the following hierarchical structure: 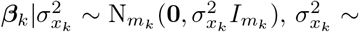 Inverse-Gamma(*a_k_, b_k_*), 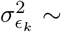 Inverse-Gamma(*a_ϵ_, b_ϵ_*), 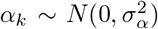. In the prior, *θ*_1_,…,*θ_K_* are independently distributed. Define the posterior probability that the phenotype of individual *i* be assigned tissue *k* as:

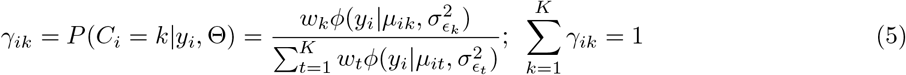

Thus, the choice of the tissue of interest across individuals is quantified by Γ = {*γ_ik_; i* = 1,…, *n*; *k* = 1,…,*K*}. Next, we define the total membership weight of *k^th^* tissue-specific subtype: 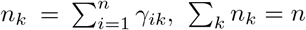. In the expectation-maximization algorithm, the main component which we maximize is given by: 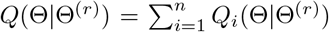, where the conditional expectation 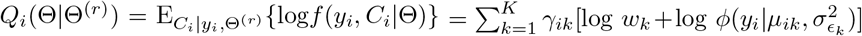. To obtain Θ^(*r*+1)^, we maximize {*Q*(Θ|Θ^(*r*)^)+log *f*(Θ)} in the MAP-EM algorithm implementing the Bayesian framework, and maximize only *Q*(Θ|Θ^(*r*)^) in the EM algorithm implementing the frequentist framework [42,43]. The steps of the MAP-EM and EM algorithm are provided in Algorithm 1 and 2, respectively. Our main inference is based on the posterior probability matrix Γ = {*γ_ik_; i* = 1,…, *n; k* = 1,…, *K*}. For example, *γ*_*i*1_ > 65% indicates that tissue 1 is likely to be the tissue of interest for individual *i*. We also designed a MCMC algorithm to implement the eGST model which we describe in Algorithm S1.

**Algorithm 1.**
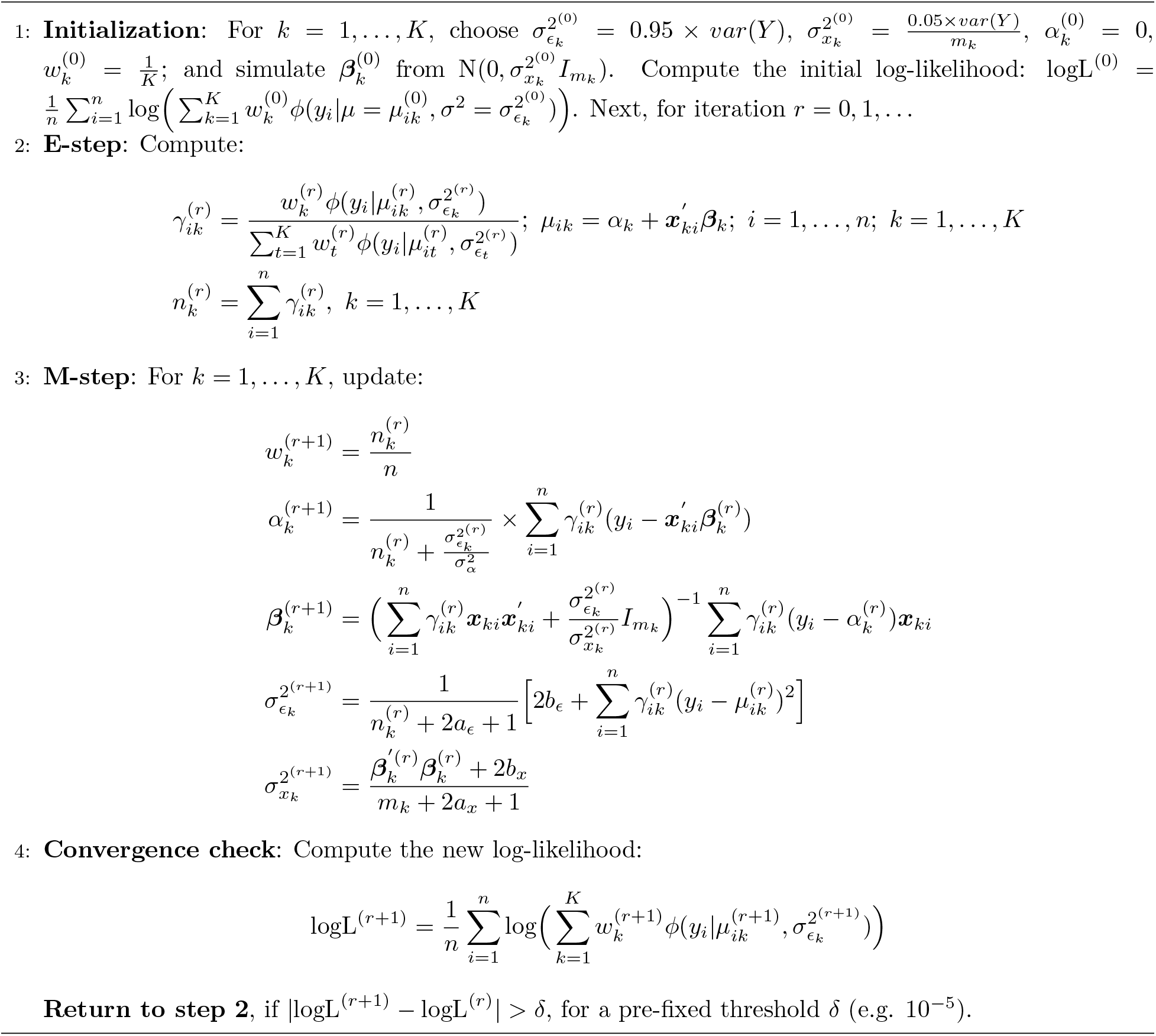
Maximum a posteriori (MAP) expectation maximization (EM) algorithm under Bayesian framework of the mixture model

**Algorithm 2.**
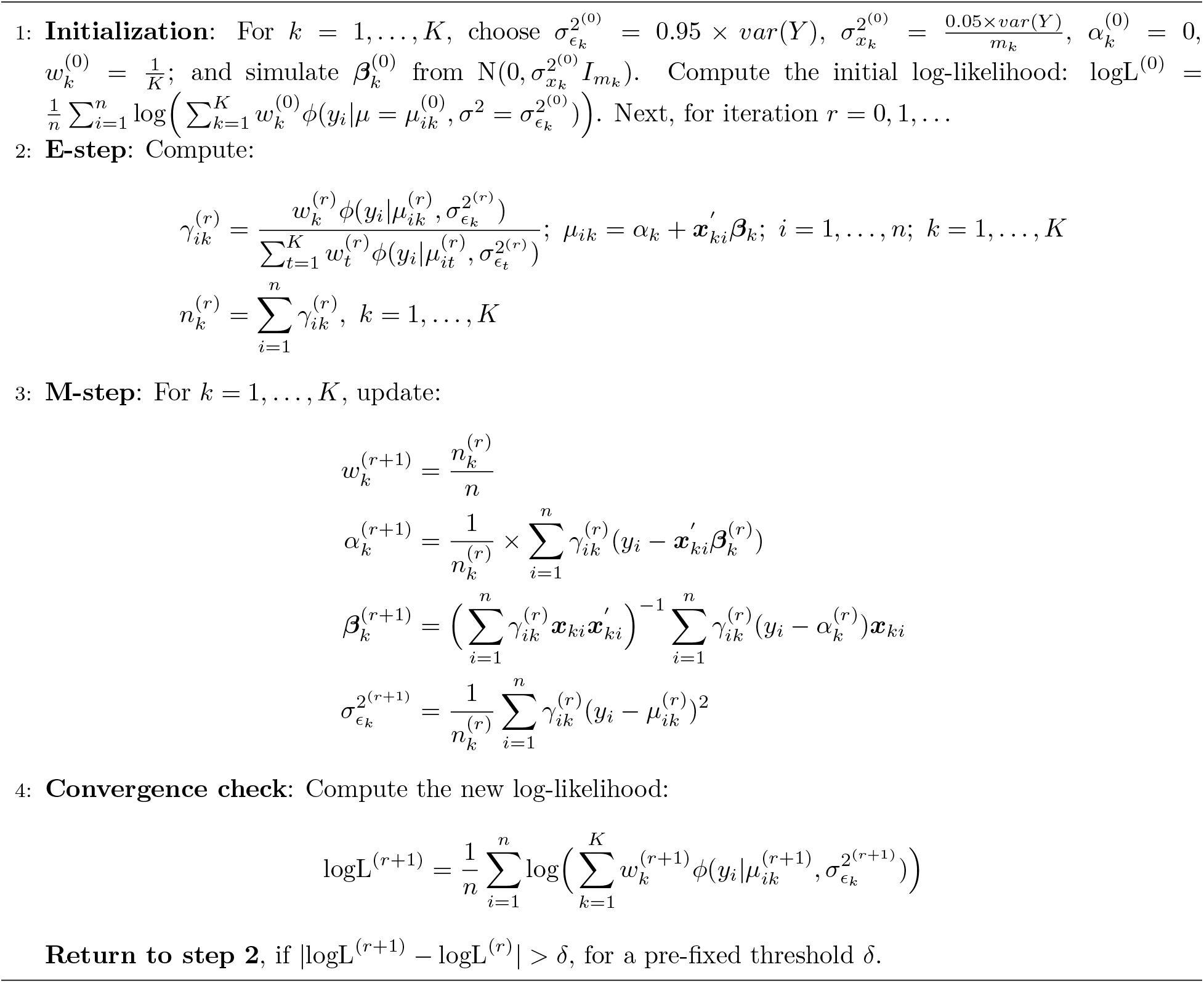
Expectation Maximization (EM) algorithm under frequentist framework of the mixture model

### Simulation design and choice of parameters

Consider *n* individuals and two non-overlapping sets of *m*_1_ SNPs and *m*_2_ SNPs representing eQTL SNP sets specific to two tissues. We chose the SNPs on chromosome 8 — 17 from the array SNPs in the UK Biobank (UKB). We pruned for LD between the SNPs such that two consecutive SNPs (on a chromosome) included in a SNP set had *r*^2^ < 0.25 (based on UKB in-sample LD). Each SNP had MAF > 1% and satisfied Hardy Weinberg Equilibrium (HWE). We collected genotype data at both sets of SNPs for *n* individuals that were randomly selected from 337,205 white-British individuals in the UKB.

Let *w* = (*w*_1_, *w*_2_) denote the proportions of individuals in the sample assigned to the two tissues where (100 × *w*_1_)% individuals are assigned the first tissue-specific subtype and (100 × *w*_2_)% individuals are assigned the second tissue-specific subtype. We assume that *m_k_* SNPs explain 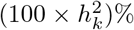 of the total variance of *k^th^* tissue-specific subtype, *k* = 1, 2. So 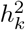 is the heritability of *k^th^* tissue-specific subtype of the trait due to *m_k_* SNPs representing *k^th^* tissue-specific eQTLs, *k* = 1, 2. Thus, if first subtype of *Y* has a total variance 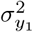, we draw each element of *β*_1_ as: 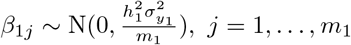. Similarly we simulate *β*_2_, the genetic effect of second set of *m*_2_ SNPs on the second subtype of *Y*. For simplicity, we assume 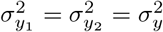, but the performance of eGST remains similar for other choices of this parameter. If the genetic susceptibility of an individual’s phenotype was assigned to be mediated through first tissue, we simulated the phenotype as: 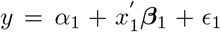, where *x*_1_ is the normalized genotype values of the individual at the first set of SNPs. While simulating the phenotype, we normalized the genotypes at each of first tissue-specific *m*_1_ SNPs only based on the individuals assigned to the first tissue-specific subtype. However, when applying eGST on a simulated dataset, we normalized the genotypes at each SNP based on all *n* individuals in the sample, because the tissue of interest across individuals are unknown. The random error components have the following distribution: 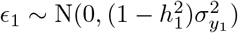 and 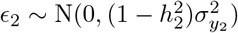.

We varied the choice of parameters to evaluate eGST in various simulation scenarios. We chose 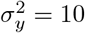, and initially assumed *α*_1_ = *α*_2_ =0 and simulate *β*_1_, *β*_2_ from zero-mean normal distributions. We considered all possible combinations of (*w*_1_, *w*_2_) where 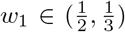 and 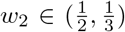, and all possible combinations of 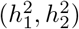, where 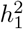 and 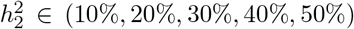. We also considered two unrealistic scenarios of null and high subtype heritability: 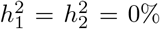 and 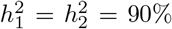 to evaluate if eGST is performing as expected in these extreme scenarios. We chose (*m*_1_,*m*_2_) with *m*_1_ = 1000,1500 and *m*_2_ = 1000,1500. Initially we chose *n* = 40,000, and later *n* = 100, 000 to explore the effects of an increased sample size. For each choice of the complete set of simulation parameters, we summarized the results of eGST across 50 simulated datasets. We also performed simulations for *α*_1_ ≠ *α*_2_ and different non-zero mean of *β*_1_, *β*_2_ distributions.

### BMI and WHRadjBMI analysis in the UK Biobank integrating GTEx data

We implemented eGST to infer the individual-level tissue of interest for two obesity related measures, BMI and WHRadjBMI, in the UK Biobank [32,33], integrating expression data from the Genotype-Tissue Expression (GTEx) project [16,18]. Sets of tissue-specific expressed genes were obtained from Finucane et al. [23] who considered a gene to be specifically expressed in a tissue of interest if the gene’s mean expression in the tissue is substantially higher than its mean expression in other tissues combined, and calculated a t-statistic to rank the genes with respect to higher expression in a specific tissue. Similar to their work [23], we considered the top 10% of all genes (2485 such genes) in a tissue, ranked according to descending value of the t-statistic, as the set of genes specifically expressed in the tissue.

We focused on the adipose and brain tissue for BMI, and the adipose and muscle tissue for WHRadjBMI. We took the union of the sets of genes specifically expressed in adipose subcutaneous and adipose visceral tissues, and considered it as the adipose-specific gene set. Similarly, we took the union of sets of genes specifically expressed in the brain cerebellum and brain cortex regions (these two had maximum sample size among different brain regions) to create a brain-specific set of genes. We excluded the genes overlapping between these two sets to consider non-overlapping sets of adipose and brain specific genes. For WHRadjBMI, we considered adipose subcutaneous and muscle skeletal connective tissue, and excluded the genes overlapping between the two sets of top 10% expressed genes within the tissues. We considered genes on the autosomal chromosomes 1-22. In BMI analysis, the main reason behind merging two type of adipose (or brain) tissues together to represent adipose (or brain) was to increase the number of tissue-specific eGenes per tissue. For WHRadjBMI analysis, we considered the adipose subcutaneous and muscle skeletal tissues to find different possible patterns in the performance of eGST that might be missed in BMI analysis due to merging tissues.

The subsets of primary sets of tissue-specific genes that were found to be eGenes in GTEx were included in subsequent analyses. For WHRadjBMI analysis, among the initially selected 2228 adipose subcutaneous tissue-specific genes, 1152 genes were found to be eGenes for which at least one bi-allelic SNP was reported to be an eQTL in the GTEx summary-level data (version v7). Similarly, we had 1272 eGenes for muscle skeletal tissue. In BMI analysis, we had 1887 eGenes for adipose and 1653 eGenes for brain. For each gene in a tissue, we took the top bi-allelic eQTL SNP (smallest SNP-expression association p-value) with MAF > 1%. In BMI analysis, while creating an adipose-specific set of eQTLs, if a gene was both adipose subcutaneous and visceral tissue-specific gene, we included the top eQTL of the gene in both tissues, one in subcutaneous and one in visceral. We implemented the same strategy for brain tissue, as well.

Next, we obtained the subset of SNPs from each set of tissue-specific eQTL SNPs (obtained from GTEx), which were genotyped or imputed in UKB (imputation accuracy score > 0.9). The SNPs were also screened for HWE (p-value > 10^−6^) in UKB. We LD-pruned each set of tissue-specific eQTL SNPs based on *r*^2^ threshold 0.25 using UKB in-sample LD. In a tissue-specific set, if two eQTL SNPs had *r*^2^ > 0.25, we excluded the one for which the minimum of SNP-expression association p-value (in GTEx) across the genes (for which it was found to be the top eQTL) was larger. Finally, after LD pruning, we had 1705 eQTL SNPs specific to adipose and 1478 eQTL SNPs specific to brain for BMI analysis. We obtained 953 eQTL SNPs specific to adipose subcutaneous tissue and 1052 eQTL SNPs specific to muscle skeletal tissue for WHRadjBMI analysis. We used individual-level genotype data for the tissue-specific SNP sets in UKB to infer tissue of interest across individuals. Before running eGST, we normalized genotypes at each SNP in the two tissue-specific sets based on the whole sample of individuals.

### Phenotype data

We considered the BMI of 337,205 unrelated white-British individuals in the UK Biobank (full release) and excluded individuals for whom BMI or relevant covariates (age, sex, etc.) were missing. We then adjusted BMI for age, sex, and the top 20 principal components (PCs) of genetic ancestry by linear regression and obtained the BMI residuals. We initially developed eGST assuming that each tissue-specific subtype of the trait follows a normal distribution. Since the BMI residuals obtained after the adjustment of covariates deviated substantially from the normal distribution (p-value of Kolmogorov Smirnov (KS) test for deviation from normal distribution < 2.2 × 10^−16^), we applied the rank-based inverse normal transformation on the BMI residuals, and implemented eGST for the transformed phenotype data. We adjusted WHR for BMI to obtain WHRadjBMI. We then adjusted WHRadjBMI for age, sex, and top 20 PCs of genetic ancestry. Since the WHRadjBMI residuals significantly deviated from the normal distribution, we applied the inverse normal transformation on the residuals.

#### Genetic characteristics

We contrasted the genetic basis of the groups of individuals assigned to the adipose and brain-specific subtypes of BMI. Let *β*_1_ denote the joint SNP-effects of the adipose-specific eQTLs on the BMI of the individuals classified as the adipose-specific subtype for whom the adipose-specific posterior probability obtained by eGST was > 50%. We chose a relaxed threshold of posterior probability because we used multiple linear regression (MLR) to estimate the joint SNP effects of a set of tissue-specific eQTLs on the BMI of a tissue-specific group of individuals, and MLR requires sufficiently large number of individuals (assigned to the corresponding tissue-specific subtype) in the sample for efficient estimation of the model parameters. Thus, if *Y*_1_ = {*Y*: *C_i_* = 1} and *X*_1_ is the genotype matrix of adipose eQTLs for these adipose-specific individuals, we fit the linear model: *E*(*Y*_1_) = *X*_1_*β*_1_. Let *γ*_1_ be the joint SNP effects of the brain eQTLs on BMI of individuals assigned to the adipose subtype. Since the BMI of individuals assigned to the adipose subtype should have larger effects from adipose-specific eQTLs than from brain-specific eQTLs, we should expect that the magnitude of a general element in *β*_1_ would be larger than the magnitude of a general element in *γ*_1_. Based on the individuals assigned to the adipose subtype, we estimated *β*_1_ and *γ*_1_ using multiple linear regression of BMI residual on the genotypes of adipose eQTLs and brain eQTLs in UKB, respectively. Based on the estimated *β*_1_ and *γ*_1_ vectors, we performed the non-parametric Wilcoxon rank sum (WRS) test to evaluate *H*_0_: |*β*_1_| = |*γ*_1_| versus *H*_1_: |*β*_1_| > |*γ*_1_|, where *β*_1_ and *γ*_1_ represent a general element in *β*_1_ and *γ*_1_ vectors, respectively. Similarly, we tested if the magnitude of the effect of a brain eQTL SNP on the BMI of individuals assigned to brain-specific subtype (*β*_2_) was larger than the corresponding effect magnitude of an adipose eQTL SNP (*γ*_2_). We also tested whether the adipose eQTLs had a larger SNP effect on the adipose subtype of BMI than on the brain subtype, and whether brain eQTLs had a larger effect on the brain subtype than on the adipose subtype. We performed the analogous experiments for the groups of individuals assigned to AS and MS tissue-specific subtype of WHRadjBMI.

#### Phenotypic characteristics

We explored if the group of individuals whose BMI were classified as a tissue-specific genetic subtype is phenotypically distinct from the rest of the population, with respect to various other phenotypes collected in the UK Biobank. We considered 106 such phenotypes and individually tested each trait for being differentially distributed between individuals of each tissue-specific subtype and the remaining population (for BMI, 11,838 individuals assigned to adipose subtype and 13,354 individuals assigned to brain subtype based on 65% threshold of subtype posterior probability [Table S1]). We performed the Wilcoxon rank sum (WRS) test for a quantitative trait and *χ*^2^ test based on the contingency table for a qualitative/categorical trait. We corrected the p-values for multiple testing across traits using the Bonferroni correction procedure. The same approach was adopted to find the traits differentially distributed between individuals classified as a tissue-specific subtype of WHRadjBMI and the remaining population (for WHRadjBMI, 11,803 individuals with AS subtype and 7,238 individuals of MS subtype [Table S1]). For a binary/case-control trait, we term the percentage of individuals (among those assigned to the tissue) who had the disorder as tissue-specific risk of the disease.

#### A random group of individuals is phenotypically homogeneous with the remaining population

For BMI, we randomly selected two groups of individuals from the population with the same size as the groups of tissue-specific BMI subtype (11,838 and 13,354) and evaluated phenotypic heterogeneity across 106 traits between each of the two random groups and the rest of the population using WRS test for a continuous trait and contingency table *χ*^2^ test for a qualitative trait (as before). We repeated the random selection of individuals from the population to replicate the experiment. We did the same experiment for WHRadjBMI.

### BMI (or WHRadjBMI) adjusted phenotypic heterogeneity

In the above analysis for BMI, BMI itself was found to be differentially distributed between the individuals with the adipose (as well as brain) specific subtype and the remaining population. Therefore, we further investigated whether the heterogeneity of non-BMI traits between a subtype group and the remaining population were induced due to BMI heterogeneity. For each quantitative trait initially found to be heterogeneous between individuals assigned to one of the subtype groups and the remaining population (Table S8), we first adjusted the trait for BMI in the whole population and obtained the trait residuals. We then tested for heterogeneity between the trait residual in the adipose (or the brain) subtype group and the remaining population using WRS test. Similarly for qualitative/categorical traits that were initially heterogeneous (Fig. 6, Table 2, Table S10), we performed a binomial or multinomial (depending on the number of categories of the trait) logistic regression adjusting for BMI in the population. We adopted the same strategy for WHRadjBMI (which itself was found to be differentially distributed between AS as well as MS group and the remaining population) to find which among the non-WHR traits remain heterogeneous after WHRadjBMI adjustment.

#### Tissue-specific relative change

For each quantitative trait that was differentially distributed between the individuals of a tissue-specific subtype and the remaining population, we measured the relative change (or difference) of the trait between the tissue-specific subtype group and the remaining population as: 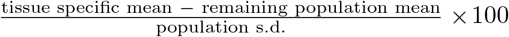, where the tissue-specific mean of the trait is calculated only in the individuals classified as the corre-sponding tissue-specific subtype of BMI (or WHRadjBMI). To quantify BMI-adjusted tissue-specific relative change of a primarily heterogeneous quantitative trait, we computed the same measure for BMI-adjusted trait residual (instead of the trait itself). To evaluate the tissue-specific relative change in the risk of a binary/case-control trait, we calculated 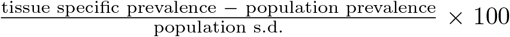, where the tissue-specific prevalence of a disease is computed only in the individuals assigned to the corresponding tissue-specific subtype.

#### BMI (or WHRadjBMI) matched tissue-specific relative change

In order to further investigate the role of tissue-specific genetics (uncoupled from the role of BMI heterogeneity) underlying the phenotypic characteristics of the individuals assigned to a tissue-specific subtype of BMI, we performed the following experiment. We split the range of BMI of the individuals assigned to the adipose subtype (11,838 individuals [Table S1]) into 30 consecutive non-overlapping bins. In each BMI bin, we counted the number of individuals assigned to the adipose subtype, and randomly sampled the same number of individuals from all of the individuals contained in the bin. In this way, we randomly selected a pool of individuals (with the same size as the adipose-specific group) from the population, who are matched with the BMI of the adipose subtype individuals. Next, for each non-BMI quantitative trait which was found to be heterogeneous between the adipose group and the remaining population after BMI adjustment (Table S13), we computed: 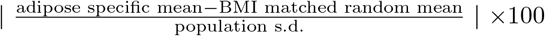, where the adipose specific mean of the trait is calculated only in the individuals of the adipose subtype and the BMI matched random mean is the trait mean calculated only in the BMI matched (with adipose group) random pool of individuals. This measure quantifies the relative change/difference of the trait between the individuals assigned to the adipose subtype and the corresponding BMI-matched random individuals selected from the population. This should provide insights into the phenotypic characteristics of the individuals with the adipose subtype, which is solely mediated through adipose-specific genetics (uncoupled from the corresponding effect of BMI heterogeneity between the adipose group and the remaining population). We repeated the random selection of BMI-matched individuals 500 times and computed the mean and s.d. of the above measure of BMI-matched tissue-specific relative change of a quantitative trait across random selections. We replicated the same experiment for individuals with brain subtype of BMI. For WHRadjBMI, we performed the same experiment to characterize the phenotypic characteristics of AS (or MS) subtype group induced due to AS-(or MS-) specific genetics only.

#### Tissue-specificity of phenotypic characteristics

To investigate tissue-specificity of the phenotypic characteristics of the individuals assigned to adipose and brain specific subtype of BMI, we randomly shuffled/exchanged 739 (half of the minimum of number of adipose and brain specific 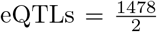) eQTLs between the set of adipose and brain specific eQTLs to create artificial tissue-specific eQTL sets. We considered 500 such random shuffles. Keeping the phenotype data fixed, for the genotype data at each set of artificial tissue-specific eQTLs, we ran eGST to identify the groups of individuals with the BMI subtype specific to the artificial adipose and brain tissues (based on the posterior probability threshold of 65%). Next, for each quantitative trait that was found to be primarily heterogeneous between the individuals assigned to the original adipose (or brain) subtype of BMI and the remaining population (Table S8), we computed the artificial adipose and brain tissue-specific trait mean only in the individuals classified into the corresponding artificial tissue-specific subtype of BMI. Then for each trait, we computed central tendency measures of the artificial tissue-specific trait means across 500 sets of artificial tissue-specific eQTLs. For each trait, we also tested whether the overall mean of the artificial tissue-specific trait means is significantly different from the corresponding original (adipose or brain) tissue-specific trait mean. We performed the same experiment for WHRadjBMI.

#### Permuting phenotype data across individuals

Next, we performed a similar experiment for permuted phenotype (BMI or WHRadjBMI) data while keeping the eQTL assignment to tissues fixed as it is in the original data. We consider 500 random permutations of BMI across individuals. Keeping the genotype data fixed, we ran eGST for each permuted phenotype data and classified the tissue of interest across individuals based on 65% threshold of subtype posterior probability. As before, in each of these 500 pairs of subtype groups of individuals thus obtained, subtype-specific means were computed for each quantitative trait that was found to be primarily heterogeneous between the individuals of the original adipose (or brain) subtype of BMI and the remaining population (Table S8). For each trait, we then computed central tendency measures of the tissue-specific means across 500 random BMI permutations. For each trait, we tested whether the overall mean of the tissue-specific trait means obtained across random permutations was significantly different from the corresponding original (adipose or brain) tissue-specific trait means. We conducted the same experiment for WHRadjBMI.

### URLs

eGST R-package: **https://cran.r-project.org/web/packages/eGST/index.html**

Tissue-specific expressed genes: **https://data.broadinstitute.org/alkesgroup/LDSCORE/LDSC_SEG_ldscores/**

GTEx portal: **https://gtexportal.org/home/**

UK Biobank: **https://www.ukbiobank.ac.uk**

PLINK: **https://www.cog-genomics.org/plink2**

## Acknowledgments

This research was conducted using the UK Biobank Resource under applications 24129 and 33297. We thank the participants of UK Biobank for making this work possible. We sincerely thank Malika Kumar Freund for her help with graphical presentation in the paper. We thank Huwenbo Shi, Kathryn Burch, Nicholas Mancuso, Hilary Finucane, Eran Halperin, Michael Gandal, Andy Dahl and Noah Zaitlen for helpful discussions relating to this work. This work was funded in part by the National Institutes of Health (NIH) under awards R01HG009120, R01MH115676, R01HG006399, U01CA194393, T32NS048004, T32MH073526 and T32HG002536.

## Supplementary materials for

### Markov Chain Monte Carlo (MCMC) algorithm for eGST

Our model for the phenotype of individual *i* is:

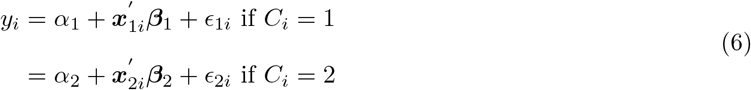

Here, *α_k_* is the baseline tissue-specific trait mean, *x_ki_* is the vector of normalized genotype values of individual *i* at the eQTL SNPs specific to tissue *k*, *β_k_* = (*β*_*k*1_, *β*_*k*2_ …, *β_km_k__*) are their effects on the trait under *C_i_* = *k*, and *ϵ_kj_* is a noise term, *i* = 1,…, *n* and *k* =1, 2. The random errors are distributed as: 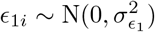 and 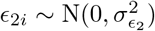.

Given that *i* = 1, we note that: 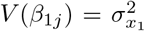 and 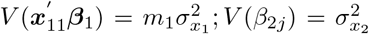 and 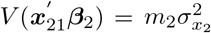. We define *τ*_1_ and *τ*_2_ such that: 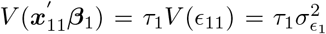 and 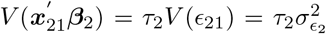. Thus, under *C* =1, total variance of the trait is: 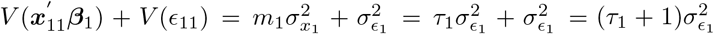. Hence, when *C* =1, the heritability of the trait is: 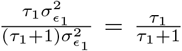. So, *k^th^* tissue-specific subtype heritability of the trait is 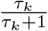. Since, specifying 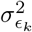 and *τ_k_* determine 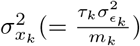, we place prior distributions on 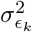 and *τ_k_* and update these parameters in the MCMC. From the MCMC sample of *τ_k_*, we can estimate the *k^th^* tissue-specific subtype heritability of the trait. The prior distributions of the *k^th^* tissue-specific set of parameters are given by: for *k* = 1, 2,

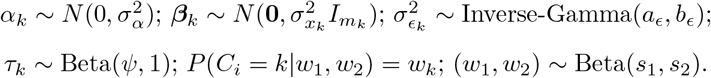

For ease in presentation of the MCMC algorithm, we define the following terms:

1. *m_k_* = number of *k^th^* tissue-specific eQTLs.
2. *n_k_* = #{*i: C_i_* = *k*} = number of individuals assigned to *k^th^* tissue-specific subtype.
3. 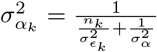
4. *G_k_* is a (*m_k_* × *n_k_*) matrix which is a sub-matrix of *X_k_* of order (*m_k_ × n*), the columns of *G_k_* are selected from the columns of *X_k_* corresponding to the individuals such that *C_i_* = *k*. Of note, *G_k_* and *X_k_* both have *m_k_* rows. ***y**_k_* = {*y_i_* — *α_k_*: *C_i_* = *k*}, i.e., it is a sub-vector of (***y*** — *α_k_*) corresponding to the individuals such that *C_i_ = k*. So, the sub-vector has length *n_k_*.

**Algorithm S1.**
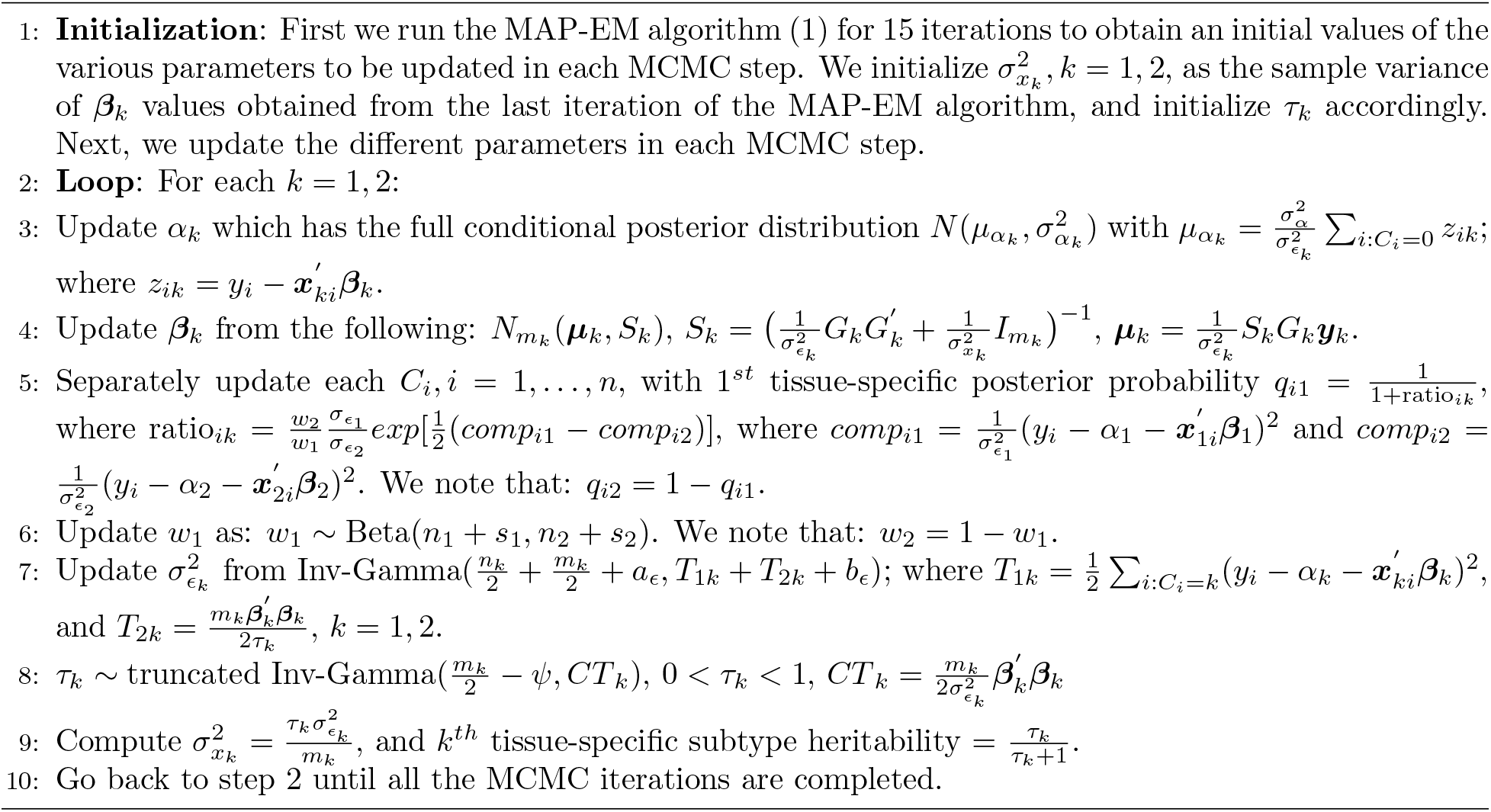
MCMC algorithm: Gibbs sampling to implement eGST

### Phenotypic characteristics of AS and MS tissue-specific groups of individuals for WHRadjBMI

We explored the phenotypic characteristics of the individuals assigned with a prioritized tissue for WHRadjBMI [11,803 AS (adipose subcutaneous) specific subtype individuals and 7,238 MS (muscle skeletal) specific subtype individuals (Table S1)]. We considered 106 phenotypes in the UK Biobank and tested each one for being differentially distributed (heterogeneous) between individuals of each tissue-specific subtype and the remaining population (see Methods). In aggregate, 37 quantitative traits and 37 qualitative traits (total 74 among 106) were significantly heterogeneous between at least one of the WHRadjBMI-AS or WHRadjBMI-MS specific groups versus the remaining population. For example, various *blood counts, body size measures, medical conditions, urine assays, townsend deprivation index, age at which full time education completed* (Table S9), *sex (male/female), diabetes proportion,* various mental health phenotypes (Table S11), *alcohol intake frequency, sleeplessness insomnia* (Table S12), etc., were differentially distributed between one or both tissue-specific subtype groups of individuals versus the remaining population. But, none of these 106 traits was found to be differentially distributed between a random set of individuals from the population (with the same size as a tissue-specific subtype group) and the remaining population (see Methods).

23 quantitative and 28 categorical traits showed heterogeneity in both the AS group versus the population, and the MS group versus population. We found 6 quantitative and 5 qualitative traits heterogeneous for individuals in the AS group but not the MS group, and found 8 quantitative and 4 qualitative traits heterogeneous for individuals in the MS group but not in the AS group (Table S9, S11, S12). We can interpret the following finding as subtype specificity: while *townsend deprivation index at recruitment* and *neuroticism score* were heterogeneous for both AS and MS tissue-specific subtypes; *creatinine enzymatic in urine* was heterogeneous only for AS and *basophil count* only for MS (Table S9). Similarly, *sex* (male/female) and *diabetes diagnosed by doctor* were heterogeneous for both tissue-specific subtypes but *risk taking* was heterogeneous only for AS and *pregnant* was heterogeneous for MS (Table S11).

For a majority of the quantitative traits (except *reticulocyte count, monocyte count, non cancer illness code selfreported*), the relative difference of the trait between individuals of a tissue-specific subtype and the remaining population (tissue-specific relative change, see Methods) are in the same direction across tissues (Fig. S5). Among binary traits, *ever smoked* was more prevalent in MS-specific group and less prevalent in AS-specific group when compared to the population (Fig. S6). We observe that for most of the case-control traits, both the tissue-specific groups of individuals had a higher risk of developing the disease compared to the population (Fig. S6). When the tissue-specific relative change of the traits (see Methods) were in the same direction across tissues, they were of different magnitude for a majority of the traits (Figure S5, S6).

Since WHRadjBMI itself was differentially distributed between the individuals of AS (as well as MS) specific subtype compared to the remaining population, we investigated whether the heterogeneity of 72 non-WHR traits (Table S9, S11, S12) were induced due to WHRadjBMI heterogeneity (see Methods). After WHRadjBMI adjustment, 32 quantitative traits (Fig. S5) among 37 primarily heterogeneous quantitative traits (Table S9) remained heterogeneous between at least one of AS and MS specific groups and the remaining population. Similarly, out of 37 qualitative traits, 27 traits (Fig. S6, Table S12) remained heterogeneous for at least one of the tissue specific groups after WHRadjBMI adjustment. These results are consistent with unique phenotypic characteristics of these individuals beyond the main phenotype effect. The WHRadjBMI-adjusted tissue-specific relative change of a quantitative trait compared to the population was computed based on the trait residuals (obtained after WHRadjBMI adjustment in the population) instead of the trait itself (see Methods). For some phenotypes, WHRadjBMI-adjusted tissue-specific relative change was in opposite direction across tissues, but in the same direction for other phenotypes (Fig. S5). Since we used linear regression while evaluating WHRadjBMI-adjusted tissue-specific relative change of heterogeneous non-WHR quantitative traits (Fig. S5, Table S14), we also investigated a model-free WHRadjBMI random matching strategy. We assessed the magnitude of relative change of a trait between individuals with the AS (or MS) subtype and a group of WHRadjBMI-matched random individuals drawn from the population (see Methods). For example, the magnitude of primary AS-specific relative change (prior to WHRadjBMI matching) for *sitting height* and *standing height* decreased from 16% and 22% to 4% and 3% after WHRadjBMI matching, respectively (Table S16).

**Figure S1:**
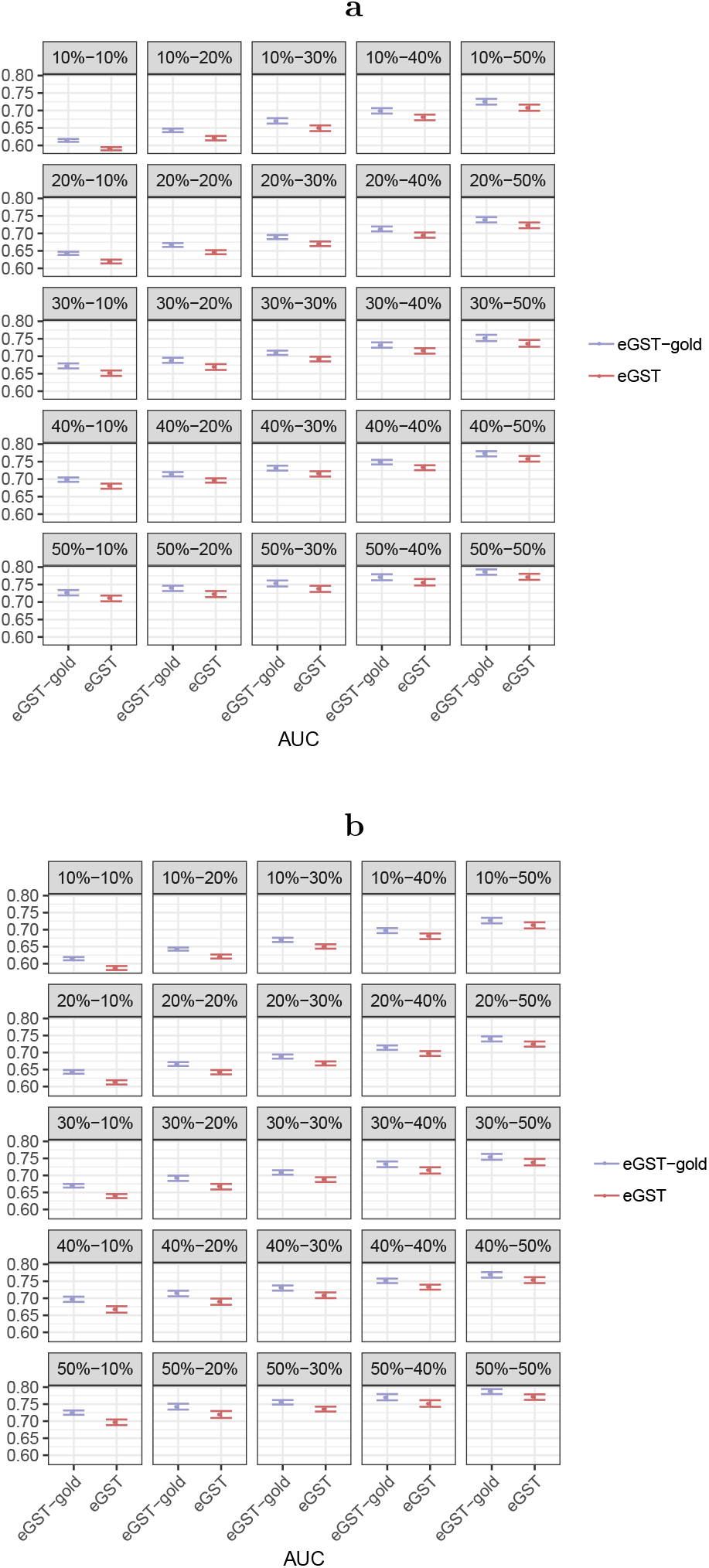
Box plots of AUCs comparing the classification accuracy of eGST and the gold-standard strategy implementing our model under the following simulation scenarios: ***m***_1_ = ***m***_2_ = **1000**, all possible combinations of 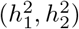 with 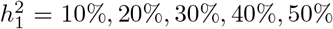 and 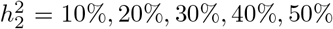 *n* = 40000; (a) 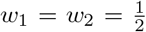 and (b) 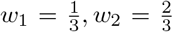. Here 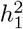 and 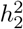 are the heritabilities of tissue-specific subtypes of the trait due to *m*_1_ and *m*_2_ SNPs representing two sets of tissue-specific eQTL SNPs, *w*_1_ and *w*_2_ are the proportions of individuals in the sample assigned to the two tissues, n is the total number of individuals. The box plots of AUC were constructed based on 50 datasets simulated under each scenario. In the gold standard strategy, true model parameters were assumed to be known while estimating the tissue-specific posterior probabilities.

**Figure S2:**
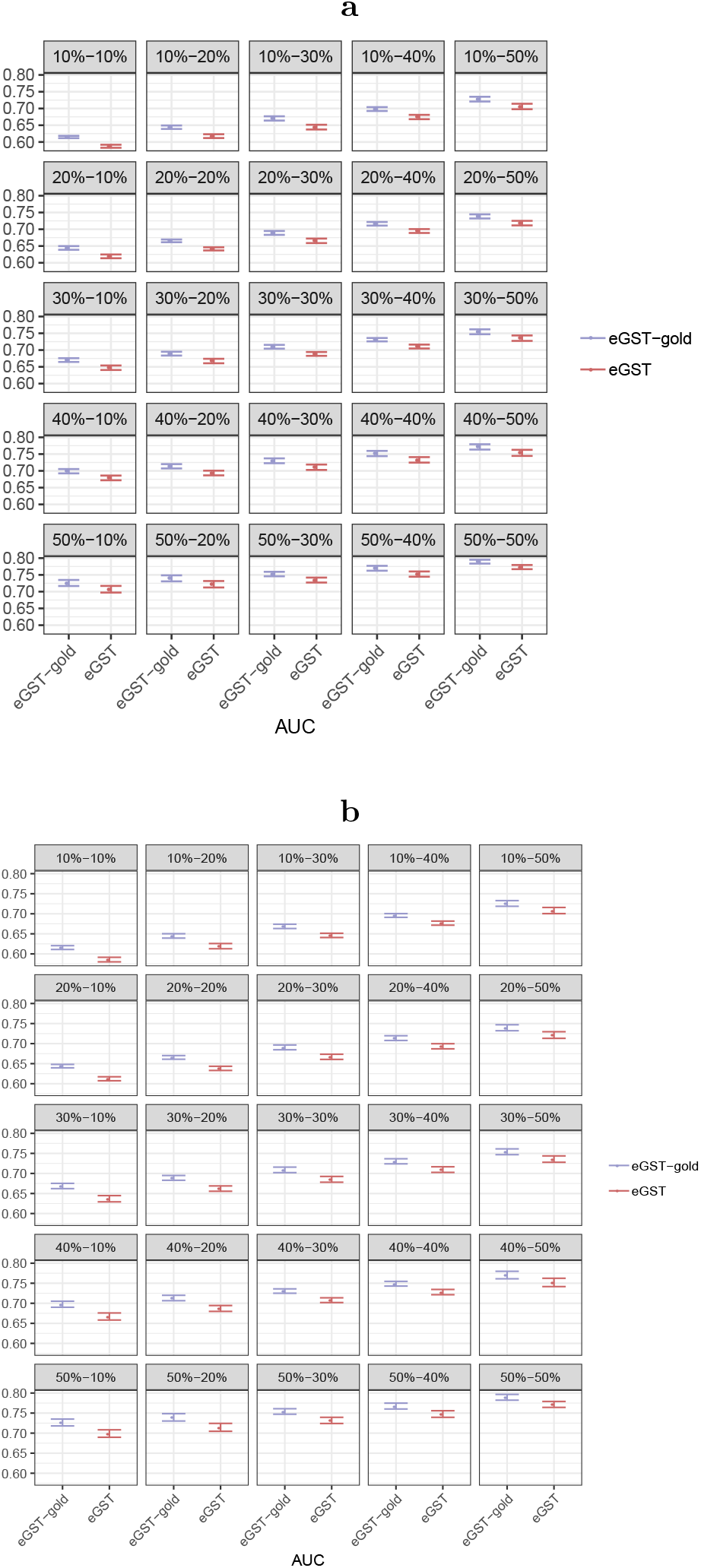
Box plots of AUCs comparing the classification accuracy of eGST and the gold-standard strategy implementing our model under the following simulation scenarios: ***m***_1_ = **1000**, ***m***_2_ = **1500**, all possible combinations of 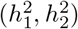 with 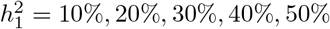 and 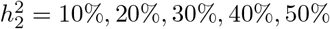; *n* = 40000; (a) 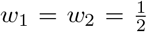 and (b) 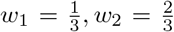. Here 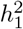 and 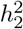 are the heritability of tissue-specific subtypes of the trait due to *m*_1_ and *m*_2_ SNPs representing two sets of tissue-specific eQTL SNPs, *w*_1_ and *w*_2_ are the proportion of individuals in the sample assigned to two tissues, n is the total number of individuals. The box plots of AUC were constructed based on 50 datasets simulated under each scenario. In the gold standard strategy, true model parameters were assumed to be known while estimating the tissue-specific posterior probabilities.

**Figure S3:**
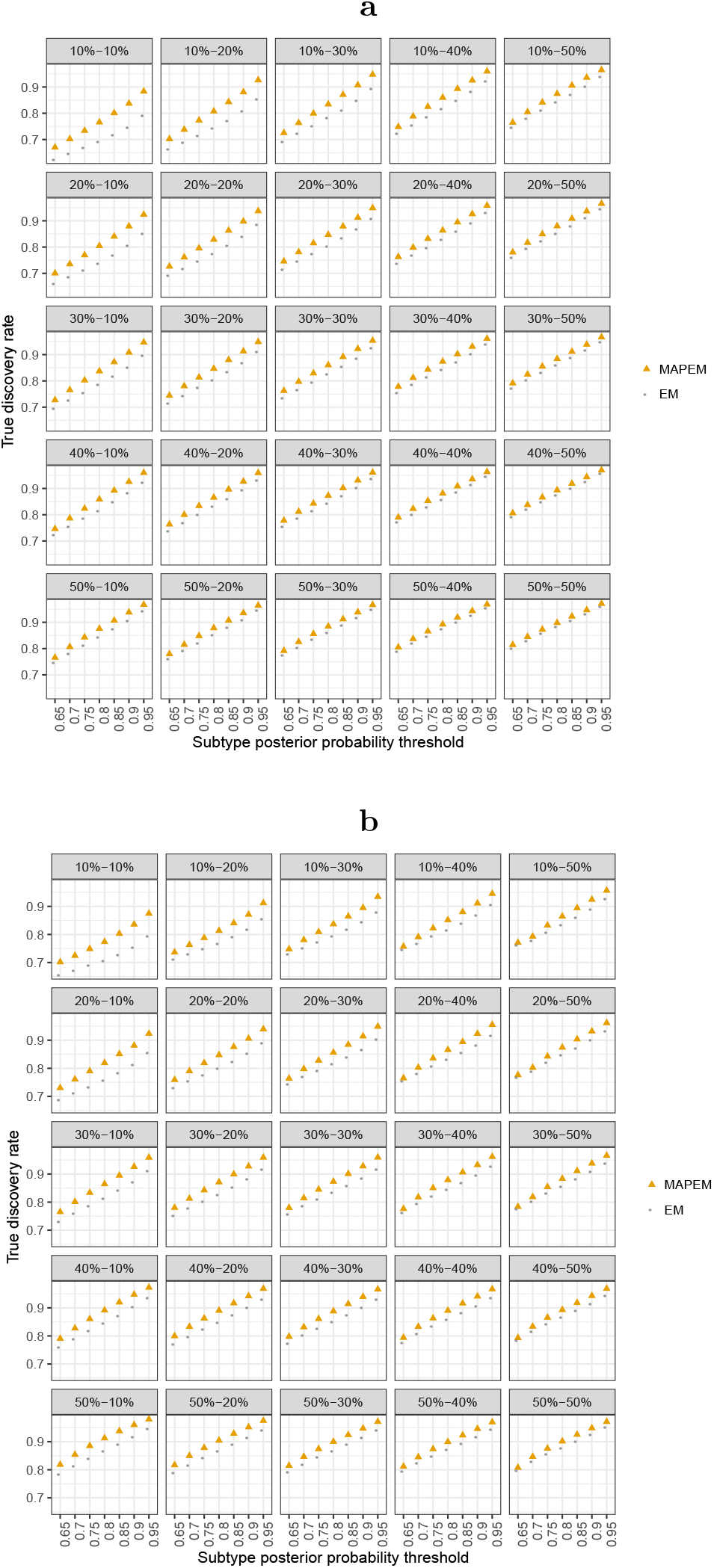
Comparison between the true discovery rate (TDR) of classifying tissue-specific subtypes by the MAP-EM algorithm (under the Bayesian framework of the mixture model which eGST employs) versus the EM algorithm (under the frequentist framework of the mixture model) based on the threshold of tissue-specific subtype posterior probability as 65%, 70%, 75%, 80%, 85%, 90%, 95%, respectively. Box plots of TDR across 50 datasets simulated under ***m***_1_ = ***m***_2_ = **1000**, combinations of 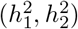 with 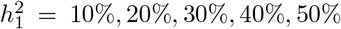 and 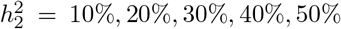; *n* = 40000; (a) 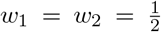 and (b) 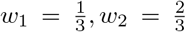, are presented. Here 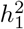 and 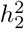 are the heritabilities of tissue-specific subtypes of the trait due to *m*_1_ and *m*_2_ SNPs representing two sets of tissue-specific eQTL SNPs, *w*_1_ and *w*_2_ are the proportions of individuals in the sample assigned to the two tissues, n is the total number of individuals.

**Figure S4:**
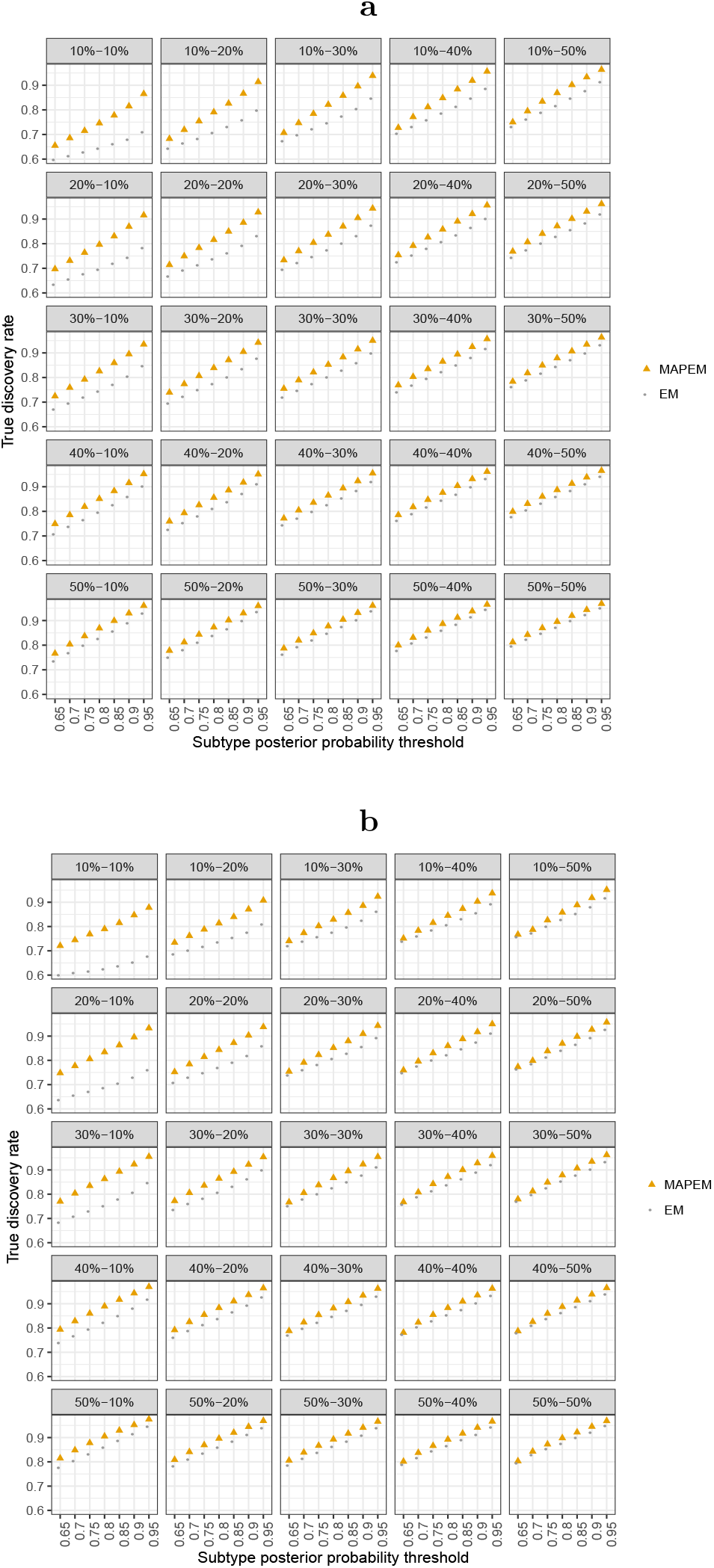
Comparison between the true discovery rate (TDR) of classifying tissue-specific subtypes by the MAP-EM algorithm (under the Bayesian framework of the mixture model which eGST employs) versus the EM algorithm (under the frequentist framework of the mixture model) based on the threshold of tissue-specific subtype posterior probability as 65%, 70%, 75%, 80%, 85%, 90%, 95%, respectively. Box plots of TDR across 50 datasets simulated under ***m***_1_ = **1000**, ***m***_2_ = **1500**, combinations of 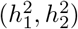 with 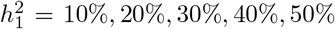 and 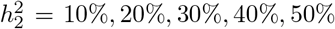; *n* = 40000; (a) 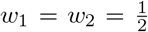 and (b) 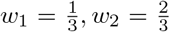, are presented. Here 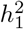 and 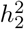 are the heritabilities of tissue-specific subtypes of the trait due to *m*_1_ and *m*_2_ SNPs representing two sets of tissue-specific eQTL SNPs, *w*_1_ and *w*_2_ are the proportions of individuals in the sample assigned to two tissues, *n* is the total number of individuals.

**Figure S5:**
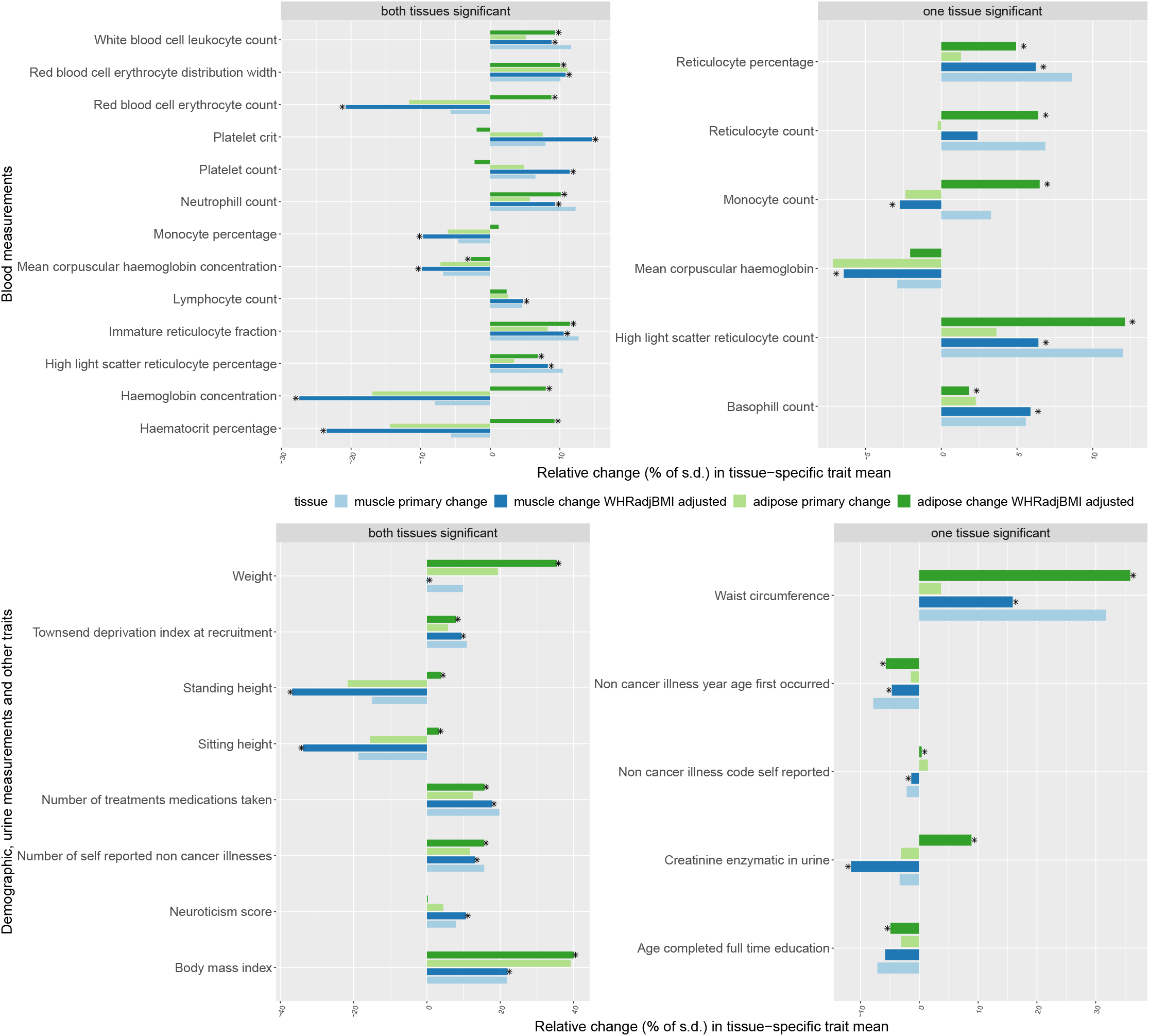
Percentage of tissue-specific relative change of the quantitative traits differentially distributed between the individuals assigned to a tissue-specific subtype of WHRadjBMI and the remaining population. Traits in the left panels are primarily heterogeneous for both tissue-specific groups and traits in the right panels are heterogenous for one tissue-specific group. We measure the tissue-specific relative change of a trait by: 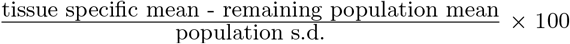, where the tissue-specific mean is computed only in the individuals with the corresponding tissue-specific subtype. The same measure is calculated for a trait residual obtained after adjusting for WHRadjBMI to quantify the tissue-specific relative change of the trait after WHRadjBMI adjustment. The faded green (or blue) bar presents primary adipose (or muscle) tissue-specific relative change of a trait compared to the remaining population. The dark green (or blue) bar presents the WHRadjBMI-adjusted adipose (or muscle) specific relative change of a trait. Each trait listed here was found differentially distributed between at least one of adipose or muscle specific group and the remaining population after WHRadjBMI adjustment. For each trait the asterisk mark attached to the bars indicates which tissue-specific group remains significantly heterogeneous after WHRadjBMI adjustment.

**Figure S6:**
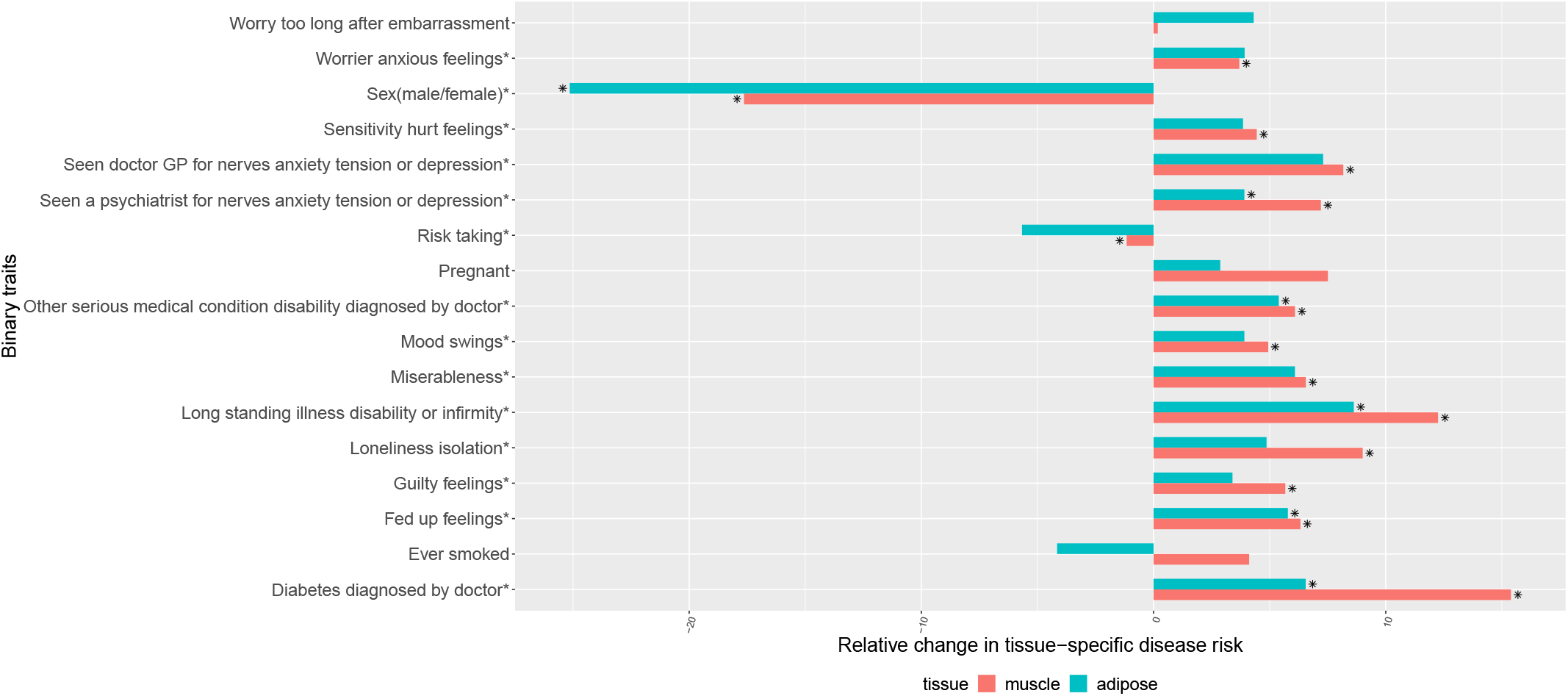
Percentage of tissue-specific relative change in the risk of case-control traits between the individuals assigned to a tissue-specific subtype of WHRadjBMI and the population. The tissue-specific relative change of a disease risk is measured by: 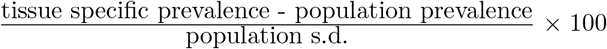. Tissue specific prevalence of the disorder was computed only in the individuals classified as the corresponding tissue-specific subtype of WHRadjBMI. The asterisk mark attached to the traits indicate which trait remains differentially distributed between at least one of adipose and muscle tissue-specific group of individuals and the remaining population after WHRadjBMI adjustment. For each trait the asterisk mark attached to the bars indicate which tissue-specific group of individuals remains significantly heterogeneous for the trait after WHRadjBMI adjustment.

**Table S1:** Number of individuals classified by eGST as a tissue-specific subtype of BMI and WHRadjBMI based on tissue-specific posterior probability threshold of 65% and 70%.

| post prob cutoff | BMI (total #individuals: 336,106) |  | WHRadjBMI (total #individuals: 336,018) |  |
| --- | --- | --- | --- | --- |
|  | Adipose | Brain | Adipose subcutaneous | Muscle skeletal |
| 65% | 11,838 | 13,354 | 11,803 | 7,238 |
| 70% | 5,397 | 6,465 | 5,319 | 2,787 |

**Table S2:** Simulation results: effect of increasing sample size on classification accuracy of eGST.

| simulation scenario | mean AUC |  |
| --- | --- | --- |
| | $n = 40,000$ | $n = 100,000$ |
| $w_1 = w_2 = 0.5, m_1 = m_2 = 1000, h_1^2 = 10\%, h_2^2 = 10\%$ | 0.59 | 0.60 |
| $w_1 = w_2 = 0.5, m_1 = m_2 = 1000, h_1^2 = 20\%, h_2^2 = 20\%$ | 0.65 | 0.66 |

**Table S3:** Simulation results: effect of increasing number of tissue-specific SNPs with fixed subtype heritability per tissue on classification accuracy of eGST.

| simulation scenario ($n = 40,000$) | mean AUC | |
| --- | --- | --- |
| | $(m_1, m_2) = (1000, 1000)$ | $(m_1, m_2) = (1500, 1500)$ |
| $w_1 = w_2 = 0.5, h_1^2 = 10\%, h_2^2 = 10\%$ | 0.59 | 0.58 |
| $w_1 = w_2 = 0.5, h_1^2 = 20\%, h_2^2 = 20\%$ | 0.65 | 0.64 |

**Table S4:**
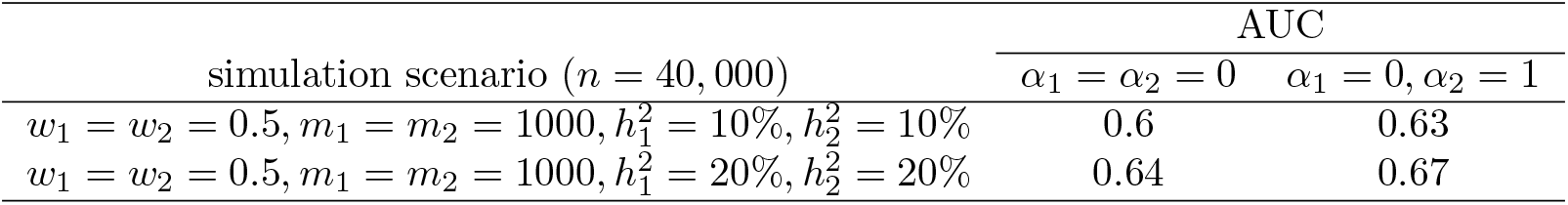
Simulation results: effect of difference between baseline tissue-specific means of the phenotype (*α*_1_, *α*_2_) on the classification accuracy of eGST.

**Table S5:** Simulation results: effect of difference between the mean of tissue-specific genetic effect size distribution on the classification accuracy of eGST. Here *j* = 1,…, *m*_1_ for *β*_1_ (tissue 1) and *j* = 1,…, *m*_2_ for *β*_2_ (tissue 2).

| simulation scenario ($n = 40,000$) | AUC | |
| --- | --- | --- |
| | $E(\beta_{1j}) = E(\beta_{2j}) = 0$ | $E(\beta_{1j}) = -0.2, E(\beta_{2j}) = 0.2$ |
| $w_1 = w_2 = 0.5, m_1 = m_2 = 1000, h_1^2 = 10\%, h_2^2 = 10\%$ | 0.6 | 0.63 |
| $w_1 = w_2 = 0.5, m_1 = m_2 = 1000, h_1^2 = 20\%, h_2^2 = 20\%$ | 0.64 | 0.68 |

**Table S6:**
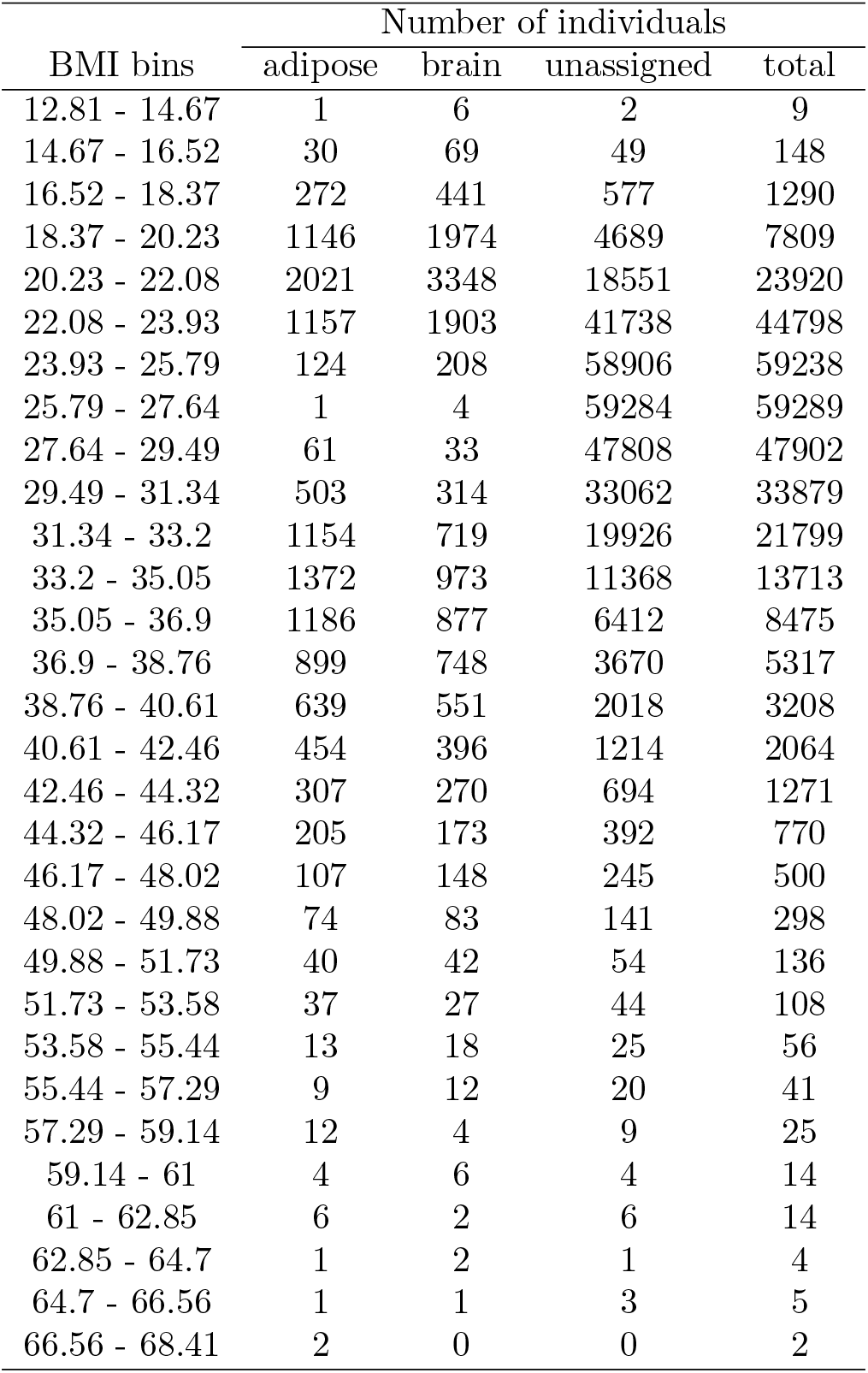
Number of individuals in consecutive non-overlapping bins of BMI who were assigned to adipose and brain specific subtype of BMI, and the number of individuals that remained unassigned by eGST based on 65% threshold of tissue-specific posterior probability.

**Table S7:** Number of individuals in consecutive non-overlapping bins of WHRadjBMI who were assigned to adipose and muscle specific subtype of WHRadjBMI, and the number of individuals that remained unassigned by eGST based on 65% threshold of tissue-specific posterior probability.

| WHRadjBMI bins | Number of individuals |  |  |  |
| --- | --- | --- | --- | --- |
|  | adipose | muscle | unassigned | total |
| -0.44 - -0.4 | 3 | 2 | 0 | 5 |
| -0.4 - -0.36 | 4 | 2 | 2 | 8 |
| -0.36 - -0.32 | 18 | 5 | 11 | 34 |
| -0.32 - -0.28 | 50 | 14 | 28 | 92 |
| -0.28 - -0.24 | 116 | 33 | 128 | 277 |
| -0.24 - -0.21 | 369 | 107 | 473 | 949 |
| -0.21 - -0.17 | 1134 | 301 | 2318 | 3753 |
| -0.17 - -0.13 | 2201 | 660 | 10377 | 13238 |
| -0.13 - -0.09 | 1311 | 410 | 29627 | 31348 |
| -0.09 - -0.05 | 524 | 172 | 47369 | 48065 |
| -0.05 - -0.01 | 1265 | 444 | 49017 | 50726 |
| -0.01 - 0.03 | 1257 | 994 | 51773 | 54024 |
| 0.03 - 0.07 | 1077 | 1169 | 56769 | 59015 |
| 0.07 - 0.11 | 712 | 762 | 43180 | 44654 |
| 0.11 - 0.14 | 671 | 948 | 19153 | 20772 |
| 0.14 - 0.18 | 647 | 818 | 5285 | 6750 |
| 0.18 - 0.22 | 298 | 289 | 1162 | 1749 |
| 0.22 - 0.26 | 102 | 71 | 216 | 389 |
| 0.26 - 0.3 | 27 | 24 | 66 | 117 |
| 0.3 - 0.34 | 11 | 9 | 11 | 31 |
| 0.34 - 0.38 | 2 | 3 | 4 | 9 |
| 0.38 - 0.42 | 1 | 0 | 1 | 2 |
| 0.42 - 0.46 | 1 | 0 | 0 | 1 |
| 0.46 - 0.49 | 0 | 1 | 2 | 3 |
| 0.49 - 0.53 | 0 | 0 | 1 | 1 |
| 0.53 - 0.57 | 0 | 0 | 0 | 0 |
| 0.57 - 0.61 | 0 | 0 | 1 | 1 |
| 0.61 - 0.65 | 0 | 0 | 0 | 0 |
| 0.65 - 0.69 | 0 | 0 | 0 | 0 |
| 0.69 - 0.73 | 1 | 0 | 0 | 1 |

**Table S8:** Quantitative traits among 106 phenotypes in UK Biobank which are differentially distributed between the adipose (and/or brain) specific subtype group of individuals for BMI and the remaining population. We provide the p-values of testing heterogeneity of each trait between each tissue-specific subtype group of individuals and the remaining population. For each trait, the tissue-specific (adipose and brain) mean which is calculated only in the individuals classified as the corresponding tissue-specific subtype of BMI are provided. We also provide the trait means computed in whole sample (popln). For each trait, we list the tissues for which the trait was differentially distributed between the corresponding tissue-specific subtype group of individuals and the remaining population (signif tissue).

| Category | Trait | adipose P | brain P | Means |  |  | signif tissue |
| --- | --- | --- | --- | --- | --- | --- | --- |
|  |  |  |  | adipose | brain | popln |  |
| Body size measures | Body mass index | 1.84E-227 | 3.01E-230 | 30.39 | 27.60 | 27.40 | both |
| Body size measures | Waist circumference | 1.06E-183 | 1.46E-108 | 95.50 | 89.20 | 90.33 | both |
| Body size measures | Weight | 8.66E-175 | 2.82E-140 | 85.20 | 77.58 | 78.31 | both |
| Body size measures | Standing height | 4.32E-64 | 2.49E-56 | 167.45 | 167.65 | 168.84 | both |
| Body size measures | Sitting height | 9.28E-28 | 2.14E-53 | 88.97 | 88.82 | 89.41 | both |
| Blood count | Haemoglobin concentration | 1.79E-48 | 3.94E-122 | 14.04 | 13.97 | 14.21 | both |
| Blood count | Haematocrit percentage | 3.60E-33 | 4.67E-98 | 40.76 | 40.54 | 41.16 | both |
| Blood count | Red blood cell erythrocyte count | 6.27E-14 | 2.90E-96 | 4.48 | 4.44 | 4.51 | both |
| Blood count | High light scatter reticulocyte percentage | 3.58E-67 | 1.54E-21 | 0.44 | 0.40 | 0.40 | both |
| Blood count | Red blood cell erythrocyte distribution width | 9.82E-62 | 1.26E-24 | 13.62 | 13.56 | 13.47 | both |
| Blood count | White blood cell leukocyte count | 1.84E-58 | 6.44E-06 | 7.15 | 6.98 | 6.89 | both |
| Blood count | High light scatter reticulocyte count | 1.09E-56 | 4.95E-37 | 0.02 | 0.02 | 0.02 | both |
| Blood count | Neutrophill count | 1.34E-55 | 1.47E-10 | 4.45 | 4.35 | 4.24 | both |
| Blood count | Reticulocyte count | 1.92E-32 | 3.69E-54 | 0.06 | 0.06 | 0.06 | both |
| Blood count | Reticulocyte percentage | 8.06E-44 | 2.39E-30 | 1.42 | 1.32 | 1.35 | both |
| Blood count | Mean corpuscular haemoglobin concentration | 2.91E-22 | 1.78E-15 | 34.46 | 34.48 | 34.54 | both |
| Blood count | Platelet crit | 1.92E-16 | 5.08E-07 | 0.24 | 0.23 | 0.23 | both |
| Blood count | Basophill count | 1.31E-13 | 7.98E-06 | 0.04 | 0.04 | 0.03 | both |
| Blood count | Monocyte percentage | 1.77E-13 | 2.29E-13 | 6.99 | 6.97 | 7.10 | both |
| Blood count | Neutrophill percentage | 1.18E-12 | 2.01E-12 | 61.67 | 61.67 | 61.14 | both |
| Blood count | Mean corpuscular volume | 2.42E-11 | 2.74E-05 | 91.09 | 91.49 | 91.34 | both |
| Blood count | Platelet count | 8.36E-10 | 1.94E-05 | 256.95 | 255.58 | 253.21 | both |
| Blood count | Lymphocyte percentage | 1.46E-09 | 1.56E-08 | 28.20 | 28.25 | 28.63 | both |
| Blood count | Monocyte count | 7.85E-07 | 0.0002 | 0.49 | 0.47 | 0.48 | both |
| Blood count | Basophill percentage | 1.38E-05 | 1.86E-05 | 0.59 | 0.60 | 0.57 | both |
| Medications | Number of treatments medications taken | 6.17E-102 | 7.84E-11 | 3.08 | 2.70 | 2.45 | both |
| Medical conditions | Number of self reported non cancer illnesses | 1.32E-84 | 6.03E-06 | 2.24 | 1.98 | 1.87 | both |
| Medical conditions | Interpolated Age of participant when<br>non cancer illness first diagnosed | 3.77E-05 | 6.52E-09 | 41.89 | 41.56 | 42.32 | both |
| Baseline characteristics | Townsend deprivation index at recruitment | 1.26E-53 | 6.69E-28 | -1.13 | -1.26 | -1.58 | both |
| Mental health | Neuroticism score | 1.41E-24 | 5.79E-19 | 4.44 | 4.38 | 4.10 | both |
| Urine assays | Creatinine enzymatic in urine | 1.51E-05 | 2.80E-24 | 9077.69 | 8429.45 | 8806.93 | both |
| Urine assays | Sodium in urine | 6.09E-13 | 3.67E-08 | 79.56 | 74.72 | 76.21 | both |
| Sleep | Sleep duration | 0.0001 | 1.92E-06 | 7.08 | 7.08 | 7.13 | both |
| Blood count | Immature reticulocyte fraction | 2.46E-69 | 0.13 | 0.30 | 0.29 | 0.29 | adipose |
| Blood count | Mean corpuscular haemoglobin | 5.73E-25 | 0.96 | 31.39 | 31.55 | 31.55 | adipose |
| Blood count | Lymphocyte count | 2.47E-15 | 0.26 | 1.98 | 1.93 | 1.95 | adipose |
| Blood count | Eosinophill count | 8.62E-07 | 0.02 | 0.18 | 0.17 | 0.17 | adipose |
| Medical conditions | Non cancer illness code self reported | 2.02E-35 | 0.05 | 2405.55 | 2779.19 | 2891.97 | adipose |
| Medical conditions | Non cancer illness year age first occurred | 1.25E-15 | 0.002 | 628.73 | 703.82 | 702.47 | adipose |
| Blood count | Mean sphered cell volume | 0.05 | 9.57E-21 | 82.86 | 83.37 | 82.87 | brain |
| Blood count | Mean reticulocyte volume | 0.09 | 2.40E-08 | 106.00 | 106.22 | 105.83 | brain |
| Blood count | Eosinophill percentage | 0.37 | 3.26E-07 | 2.53 | 2.52 | 2.56 | brain |
| Blood count | Platelet distribution width | 0.79 | 3.69E-06 | 16.49 | 16.47 | 16.49 | brain |
| Urine assays | Potassium in urine | 0.18 | 8.26E-12 | 63.61 | 61.68 | 63.47 | brain |
| Early life factors | Birth weight | 0.001 | 4.19E-06 | 3.30 | 3.30 | 3.33 | brain |

**Table S9:** Quantitative traits among 106 phenotypes in UK Biobank which are differentially distributed between the adipose subcutaneous (AS) (and/or muscle skeletal (MS)) specific subtype group of individuals for WHRadjBMI and the remaining population. We provide the p-values of testing heterogeneity of each trait between each tissue-specific subtype group of individuals and the remaining population. For each trait, the tissue-specific (AS and MS) mean which is calculated only in the individuals classified as the corresponding tissue-specific subtype of WHRadjBMI are provided. We also provide the trait means computed in whole sample (popln). For each trait, we list the tissues for which the trait was differentially distributed between the corresponding tissue-specific subtype group of individuals and the remaining population (signif tissue).

| Category | Trait | AS P | MS P | Means |  |  | signif tissue |
| --- | --- | --- | --- | --- | --- | --- | --- |
|  |  |  |  | AS | MS | popln |  |
| Body size measure | WHRadjBMI | 0 | 2.37E-192 | -0.04 | 0.03 | 0.00 | both |
| Body size measure | WHR | 3.68E-232 | 3.71E-201 | 0.85 | 0.91 | 0.87 | both |
| Body size measures | Body mass index | 6.76E-147 | 4.07E-42 | 29.19 | 28.41 | 27.39 | both |
| Body size measures | Standing height | 2.86E-124 | 1.38E-37 | 166.91 | 167.48 | 168.84 | both |
| Body size measures | Sitting height | 1.93E-70 | 1.18E-55 | 88.68 | 88.53 | 89.41 | both |
| Body size measures | Weight | 1.13E-36 | 9.02E-07 | 81.27 | 79.83 | 78.31 | both |
| Blood count | Haemoglobin concentration | 2.27E-81 | 6.20E-11 | 14.01 | 14.12 | 14.21 | both |
| Blood count | Haematocrit percentage | 1.08E-59 | 1.03E-06 | 40.67 | 40.96 | 41.16 | both |
| Blood count | Red blood cell erythrocyte distribution width | 6.69E-40 | 3.06E-14 | 13.57 | 13.56 | 13.47 | both |
| Blood count | Red blood cell erythrocyte count | 8.69E-38 | 4.27E-06 | 4.47 | 4.49 | 4.51 | both |
| Blood count | High light scatter reticulocyte percentage | 5.46E-06 | 7.63E-31 | 0.41 | 0.43 | 0.40 | both |
| Blood count | White blood cell leukocyte count | 3.41E-06 | 1.64E-28 | 6.99 | 7.12 | 6.89 | both |
| Blood count | Immature reticulocyte fraction | 3.27E-17 | 1.74E-23 | 0.29 | 0.30 | 0.29 | both |
| Blood count | Neutrophil count | 1.01E-06 | 3.91E-22 | 4.32 | 4.41 | 4.24 | both |
| Blood count | Mean corpuscular haemoglobin concentration | 1.44E-21 | 2.67E-08 | 34.47 | 34.47 | 34.54 | both |
| Blood count | Monocyte percentage | 2.51E-21 | 6.52E-10 | 6.94 | 6.98 | 7.10 | both |
| Blood count | Platelet crit | 5.69E-18 | 1.54E-11 | 0.24 | 0.24 | 0.23 | both |
| Blood count | Lymphocyte count | 0.0003 | 1.30E-15 | 1.98 | 2.00 | 1.95 | both |
| Blood count | Platelet count | 8.60E-08 | 4.87E-08 | 256.00 | 257.00 | 253.21 | both |
| Medications | Number of treatments medications taken | 9.24E-30 | 4.17E-50 | 2.77 | 2.97 | 2.45 | both |
| Medical conditions | Number of self reported non cancer illnesses | 5.95E-26 | 1.19E-34 | 2.08 | 2.15 | 1.86 | both |
| Baseline characteristics | Townsend deprivation index at recruitment | 3.34E-10 | 8.09E-17 | -1.41 | -1.26 | -1.58 | both |
| Mental health | Neuroticism score | 1.72E-06 | 2.04E-08 | 4.24 | 4.35 | 4.10 | both |
| Blood count | Mean corpuscular haemoglobin | 1.64E-18 | 0.03 | 31.43 | 31.50 | 31.55 | adipose |
| Blood count | Monocyte count | 4.67E-08 | 0.04 | 0.47 | 0.49 | 0.48 | adipose |
| Blood count | Mean corpuscular volume | 9.40E-06 | 0.74 | 91.18 | 91.38 | 91.34 | adipose |
| Blood count | Mean platelet thrombocyte volume | 9.74E-05 | 0.48 | 9.36 | 9.33 | 9.32 | adipose |
| Blood count | Eosinophil percentage | 0.0004 | 0.54 | 2.49 | 2.53 | 2.56 | adipose |
| Urine assays | Creatinine enzymatic in urine | 0.0001 | 0.02 | 8633.55 | 8618.76 | 8806.64 | adipose |
| Body size measures | Waist circumference | 0.20 | 3.17E-127 | 90.80 | 94.50 | 90.33 | muscle |
| Blood count | High light scatter reticulocyte count | 0.01 | 2.98E-25 | 0.02 | 0.02 | 0.02 | muscle |
| Blood count | Reticulocyte percentage | 0.14 | 2.25E-23 | 1.36 | 1.42 | 1.35 | muscle |
| Blood count | Reticulocyte count | 0.22 | 2.65E-17 | 0.06 | 0.06 | 0.06 | muscle |
| Blood count | Basophil count | 0.27 | 4.63E-10 | 0.03 | 0.04 | 0.03 | muscle |
| Medical conditions | Non cancer illness code self reported | 0.0007 | 1.21E-10 | 3071.24 | 2623.36 | 2891.68 | muscle |
| Medical conditions | Non cancer illness year age first occurred | 0.06 | 4.69E-06 | 689.59 | 631.65 | 702.50 | muscle |
| Education | Age completed full time education | 0.12 | 8.65E-06 | 16.36 | 16.25 | 16.45 | muscle |

**Table S10:**
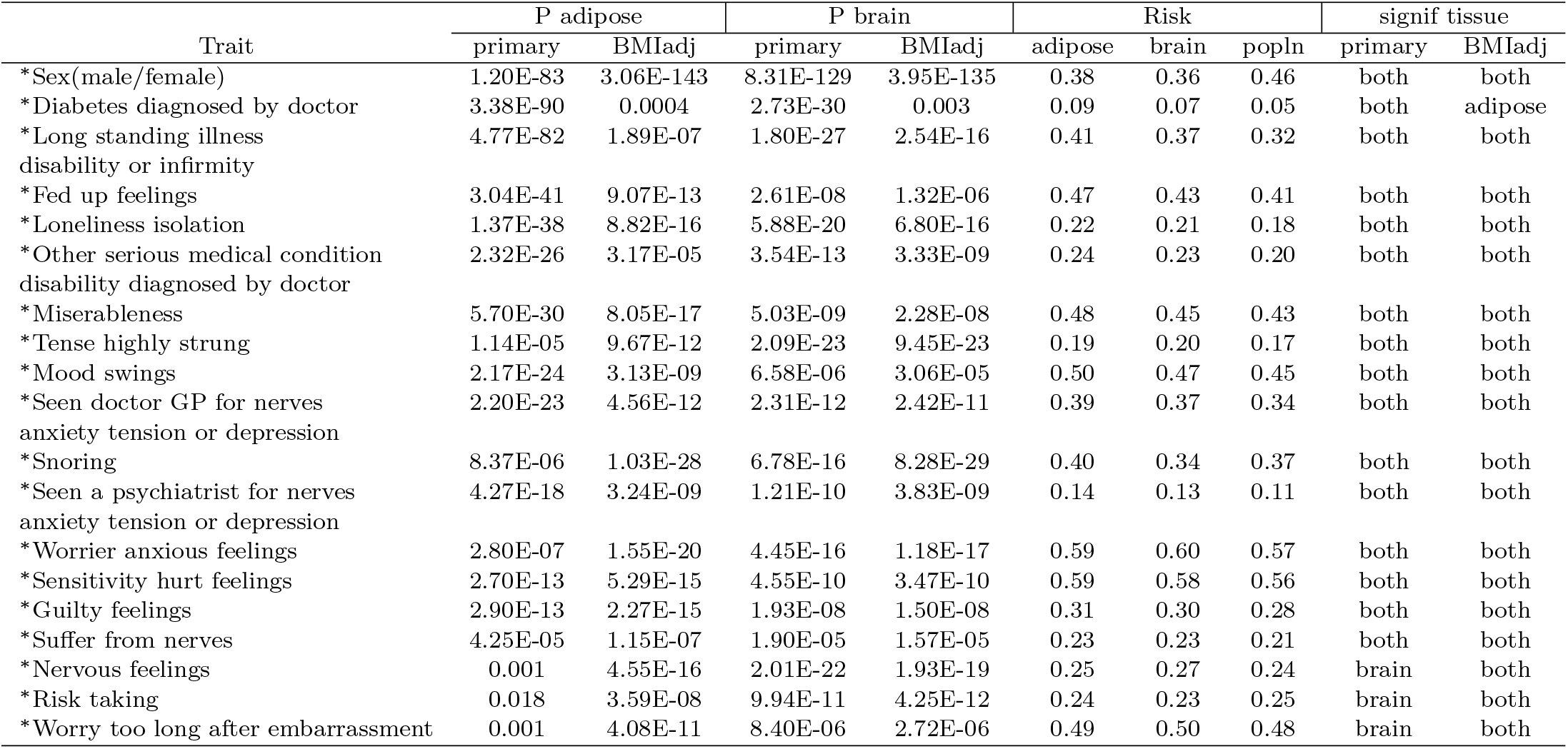
Case-control traits among 106 phenotypes considered in the UK biobank which are differentially distributed between the adipose (and/or brain) specific subtype group of individuals for BMI and the remaining population. For each trait, we provide the p-value of testing heterogeneity between each tissue-specific subtype group of individuals and the remaining population before (primary) and after BMI adjustment (BMIadj). For each trait, tissue-specific groups which appear to be significantly heterogeneous (signif tissue) before (primary) and after BMI adjustment (BMIadj) are provided. The asterisk mark attached to the traits indicate which trait remains differentially distributed between at least one of the tissue-specific groups and the remaining population after BMI adjustment. Tissue-specific risk of the disorders along with population-level risk are also provided.

**Table S11:** Case-control traits among 106 phenotypes considered in the UK biobank which are differentially distributed between the adipose subcutaneous (AS) (and/or muscle skeletal (MS)) specific subtype group of individuals for WHRadjBMI and the remaining population. For each trait, we provide the p-value of testing heterogeneity between each tissue-specific subtype group of individuals and the remaining population before (primary) and after WHRadjBMI adjustment (adjusted). For each trait, tissue-specific groups which appear to be significantly heterogeneous (signif tissue) before (primary) and after WHRadjBMI adjustment (adjusted) are provided. The asterisk mark attached to the traits indicate which trait remains differentially distributed between at least one of the tissue-specific groups and the remaining population after WHRadjBMI adjustment. Tissue-specific risk of the disorders along with population-level risk are also provided.

| Trait | P AS |  | P MS |  | Risk |  |  | signif tissue |  |
| --- | --- | --- | --- | --- | --- | --- | --- | --- | --- |
|  | primary | adjusted | primary | adjusted | AS | MS | popln | primary | adjusted |
| *Sex(male/female) | 4.37E-170 | 1.14E-08 | 6.69E-52 | 0 | 0.34 | 0.37 | 0.46 | both | both |
| *Diabetes diagnosed by doctor | 5.42E-13 | 2.68E-22 | 8.91E-40 | 4.26E-11 | 0.06 | 0.08 | 0.05 | both | both |
| *Long standing illness<br>disability or infirmity | 4.69E-21 | 3.51E-50 | 2.92E-25 | 6.61E-12 | 0.36 | 0.38 | 0.32 | both | both |
| *Seen doctor GP for nerves<br>anxiety tension or depression | 8.80E-16 | 0.01 | 2.73E-12 | 5.68E-22 | 0.38 | 0.38 | 0.34 | both | muscle |
| *Loneliness isolation | 1.02E-07 | 0.002 | 1.81E-14 | 2.61E-18 | 0.20 | 0.21 | 0.18 | both | muscle |
| *Miserableness | 2.54E-11 | 0.09 | 2.37E-08 | 3.04E-16 | 0.46 | 0.46 | 0.43 | both | muscle |
| *Fed up feelings | 2.59E-10 | 1.90E-07 | 8.04E-08 | 2.24E-09 | 0.43 | 0.44 | 0.41 | both | both |
| *Other serious medical condition<br>disability diagnosed by doctor | 3.83E-09 | 2.61E-16 | 2.36E-07 | 0.0002 | 0.23 | 0.23 | 0.20 | both | both |
| *Seen a psychiatrist for nerves<br>anxiety tension or depression | 1.70E-05 | 6.28E-06 | 6.95E-10 | 1.21E-09 | 0.13 | 0.14 | 0.11 | both | both |
| Ever smoked | 4.74E-06 | 0.004 | 0.0004 | 0.88 | 0.58 | 0.62 | 0.60 | both | none |
| *Guilty feelings | 0.0002 | 0.05 | 1.57E-06 | 5.14E-13 | 0.30 | 0.31 | 0.28 | both | muscle |
| *Sensitivity hurt feelings | 2.69E-05 | 0.02 | 0.0002 | 2.37E-14 | 0.58 | 0.58 | 0.56 | both | muscle |
| *Mood swings | 2.01E-05 | 0.002 | 2.90E-05 | 1.65E-06 | 0.47 | 0.48 | 0.45 | both | muscle |
| *Risk taking | 8.17E-10 | 0.30 | 0.34 | 1.90E-07 | 0.23 | 0.25 | 0.25 | adipose | muscle |
| Worry too long<br>after embarrassment | 3.26E-06 | 0.48 | 0.89 | 0.003 | 0.50 | 0.48 | 0.48 | adipose | none |
| *Worrier anxious feelings | 1.94E-05 | 0.38 | 0.002 | 7.28E-10 | 0.59 | 0.59 | 0.57 | adipose | muscle |
| Pregnant | 0.02 | 0.13 | 2.43E-06 | 0.01 | 0.001 | 0.002 | 0.0004 | muscle | none |

**Table S12:** Qualitative/categorical traits with three or more categories that are differentially distributed between at least one of adipose subcutaneous (AS) and muscle skeletal (MS) specific subtype groups of individuals for WHRadjBMI and the remaining population. For each trait we provide the p-values of testing heterogeneity between each tissue-specific subtype group of individuals and the remaining population before (primary) and after WHRadjBMI adjustment (adjusted). For each trait, tissue-specific groups which appear to be significantly heterogeneous (signif tissue) before (primary) and after WHRadjBMI adjustment (adjusted) are also provided. The asterisk mark attached to the traits indicate which trait remains differentially distributed between at least one of the tissue-specific groups and the remaining population after WHRadjBMI adjustment. The number of categories for each trait (#categ) are also listed.

| Trait | P AS |  | P MS |  | signif tissue |  | #categ |
| --- | --- | --- | --- | --- | --- | --- | --- |
|  | primary | adjusted | primary | adjusted | primary | adjusted |  |
| *Alcohol intake frequency | 8.90E-74 | 1.41E-07 | 3.66E-24 | 6.98E-11 | both | both | 6 |
| *Overall health rating | 2.35E-46 | 2.11E-23 | 6.34E-49 | 3.35E-11 | both | both | 4 |
| *Weight change compared with 1 year ago | 8.51E-37 | 6.67E-19 | 2.89E-17 | 5.46E-27 | both | both | 3 |
| *Alcohol drinker status | 2.93E-23 | 4.56E-14 | 7.33E-06 | 9.74E-09 | both | both | 3 |
| *Falls in the last year | 1.43E-23 | 4.22E-12 | 4.68E-14 | 1.95E-10 | both | both | 3 |
| Frequency of tiredness lethargy in last weeks | 3.69E-22 | 0.002 | 3.00E-17 | 0.12 | both | none | 4 |
| *Current tobacco smoking | 0.0004 | 0.35 | 9.55E-19 | 4.74E-05 | both | muscle | 3 |
| Qualifications | 0.0002 | 0.1 | 5.74E-17 | 1 | both | none | 7 |
| *Smoking status | 1.46E-05 | 0.35 | 1.29E-16 | 4.50E-05 | both | muscle | 3 |
| *Sleeplessness insomnia | 7.08E-13 | 0.02 | 1.02E-09 | 3.51E-10 | both | muscle | 3 |
| *Frequency of depressed mood in last 2 weeks | 2.94E-08 | 0.0002 | 2.60E-05 | 0.009 | both | adipose | 4 |
| *Blood clot DVT bronchitis emphysema asthma rhinitis eczema allergy diagnosed by doctor | 3.07E-06 | 0.07 | 1.12E-10 | 1.57E-06 | both | muscle | 6 |
| *Getting up in morning | 2.40E-06 | 0.0002 | 2.97E-11 | 0.54 | both | adipose | 4 |
| *Frequency of unenthusiasm disinterest in last 2 weeks | 2.23E-05 | 2.54E-05 | 5.29E-07 | 0.0009 | both | adipose | 4 |
| Daytime dozing sleeping narcolepsy | 3.88E-06 | 0.16 | 0.0003 | 0.46 | both | none | 4 |
| Illness injury bereavement stress in last 2 years | 1.55E-09 | 0.06 | 0.003 | 0.42 | adipose | none | 7 |
| Past tobacco smoking | 8.52E-06 | 0.02 | 0.004 | 0.24 | adipose | none | 4 |
| *Nap during day | 0.12 | 1.77E-18 | 2.65E-10 | 0.14 | muscle | adipose | 3 |
| Frequency of tenseness restlessness in last 2 weeks | 0.02 | 0.01 | 3.04E-06 | 0.05 | muscle | none | 4 |
| Morning evening person chronotype | 0.001 | 0.03 | 0.0001 | 0.43 | muscle | none | 4 |

**Table S13:** Heterogeneity of non-BMI quantitative traits between the adipose (brain) specific subtype group of individuals for BMI and the remaining population after BMI adjustment of the traits in the population using linear regression. For each trait, we provide the p-values of testing heterogeneity between each tissue-specific subtype group of individuals and the remaining population after BMI adjustment. For each trait, tissue-specific groups which appear to be significantly heterogeneous (signif tissue) after BMI adjustment are provided. To measure the primary tissue-specific relative change of a trait we calculate the following: 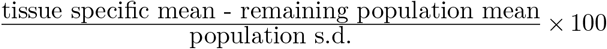, where the tissue-specific mean is computed only in the individuals with the corresponding tissue-specific subtype of BMI. The same measure is calculated for a trait residual obtained after adjusting for BMI in the population to quantify the tissue-specific relative change of the trait after BMI adjustment (BMIadj).

| Trait | adipose P | brain P | signif tissue | Tissue-specific relative change |  |  |  |
| --- | --- | --- | --- | --- | --- | --- | --- |
|  |  |  |  | adipose BMIadj | adipose primary | brain BMIadj | brain primary |
| Non cancer illness code self reported | 0 | 3.37E-06 | both | -1.46 | -3.95 | -0.26 | -0.92 |
| Interpolated Age of participant when non cancer illness first diagnosed | 3.05E-22 | 4.31E-14 | both | -7.14 | -2.43 | -5.53 | -4.28 |
| Number of self reported non cancer illnesses | 3.01E-13 | 1.60E-21 | both | 7.22 | 21.14 | 5.87 | 6.71 |
| Haemoglobin concentration | 8.50E-118 | 4.50E-125 | both | -22.72 | -14.19 | -20.99 | -20.23 |
| Red blood cell erythrocyte count | 2.98E-93 | 8.65E-108 | both | -19.83 | -7.54 | -19.81 | -18.61 |
| Haematocrit percentage | 1.52E-94 | 1.57E-102 | both | -20.01 | -11.54 | -18.92 | -18.18 |
| Reticulocyte count | 1.70E-31 | 1.24E-42 | both | -7.00 | 8.54 | -6.46 | -5.06 |
| Mean sphered cell volume | 3.28E-16 | 2.44E-27 | both | 8.84 | -0.23 | 10.69 | 9.89 |
| Red blood cell erythrocyte distribution width | 1.17E-22 | 3.01E-25 | both | 11.49 | 16.74 | 9.86 | 10.18 |
| Mean corpuscular haemoglobin concentration | 1.39E-23 | 5.61E-15 | both | -8.11 | -7.86 | -5.97 | -5.94 |
| Reticulocyte percentage | 5.39E-15 | 1.46E-23 | both | -4.37 | 8.85 | -4.00 | -2.90 |
| Monocyte percentage | 1.31E-15 | 2.68E-13 | both | -4.55 | -4.09 | -5.25 | -5.23 |
| Neutrophil percentage | 4.68E-13 | 3.96E-12 | both | 6.72 | 6.58 | 6.62 | 6.61 |
| High light scatter reticulocyte count | 1.16E-10 | 3.16E-08 | both | -5.96 | 19.22 | -5.62 | -3.31 |
| Neutrophil count | 2.40E-08 | 2.43E-10 | both | 6.02 | 15.43 | 7.03 | 7.61 |
| Basophil percentage | 1.31E-09 | 6.19E-05 | both | 5.06 | 4.48 | 4.85 | 4.77 |
| Mean reticulocyte volume | 6.71E-06 | 4.40E-09 | both | 4.74 | 2.28 | 5.38 | 5.18 |
| Lymphocyte percentage | 3.06E-08 | 2.12E-08 | both | -5.54 | -6.01 | -5.40 | -5.43 |
| Platelet crit | 2.29E-08 | 7.27E-07 | both | 5.83 | 8.44 | 4.31 | 4.50 |
| Eosinophil percentage | 5.59E-05 | 5.25E-08 | both | -3.43 | -1.46 | -2.49 | -2.35 |
| Platelet count | 1.22E-05 | 2.66E-05 | both | 4.78 | 6.48 | 4.01 | 4.13 |
| Waist circumference | 6.43E-106 | 1.15E-118 | both | -22.89 | 39.90 | -21.34 | -8.71 |
| Weight | 7.13E-71 | 1.60E-66 | both | -17.15 | 45.05 | -15.66 | -4.81 |
| Standing height | 6.79E-58 | 6.24E-63 | both | -14.74 | -15.57 | -13.37 | -13.43 |
| Sitting height | 2.15E-46 | 8.76E-48 | both | -13.52 | -9.42 | -13.10 | -12.79 |
| Number of treatments medications taken | 6.14E-17 | 9.61E-38 | both | 9.72 | 24.52 | 8.89 | 9.68 |
| Neuroticism score | 6.29E-14 | 4.23E-31 | both | 10.52 | 10.86 | 9.01 | 9.02 |
| Townsend deprivation index at recruitment | 7.86E-20 | 7.03E-29 | both | 9.85 | 15.63 | 11.01 | 11.34 |
| Creatinine enzymatic in urine | 1.36E-20 | 1.07E-13 | both | -8.62 | 4.89 | -7.73 | -6.85 |
| Potassium in urine | 3.55E-10 | 1.11E-11 | both | -6.64 | 0.42 | -5.90 | -5.49 |
| Birth weight | 3.63E-07 | 5.43E-05 | both | -5.99 | -3.70 | -4.81 | -4.65 |
| Non cancer illness year age first occurred | 1.67E-90 | 0.25 | adipose | 2.31 | -8.27 | 2.97 | 0.15 |
| Monocyte count | 1.82E-07 | 0.003 | adipose | -2.23 | 5.21 | -2.95 | -2.42 |
| Eosinophil count | 3.15E-05 | 0.18 | adipose | -2.45 | 3.56 | -0.87 | -0.44 |
| High light scatter reticulocyte percentage | 6.41E-05 | 0.003 | adipose | -2.51 | 12.78 | -2.19 | -1.00 |
| Sodium in urine | 1.45E-08 | 0.0005 | adipose | -5.73 | 7.97 | -4.45 | -3.55 |
| Basophil count | 0.004 | 1.61E-22 | brain | 5.34 | 7.50 | 5.18 | 5.30 |
| Platelet distribution width | 0.002 | 2.59E-06 | brain | -2.92 | 0.44 | -4.49 | -4.24 |
| Mean corpuscular volume | 0.07 | 4.19E-06 | brain | 1.93 | -5.86 | 4.09 | 3.49 |
| White blood cell leukocyte count | 0.001 | 1.56E-05 | brain | 2.67 | 13.43 | 3.95 | 4.67 |
| Sleep duration | 0.01 | 1.55E-54 | brain | -2.33 | -4.26 | -4.08 | -4.20 |

**Table S14:** Phenotypic heterogeneity of non-WHR quantitative traits between adipose subcutaneous (AS) (muscle skeletal (MS)) specific group of individuals for WHRadjBMI and the remaining population after WHRadjBMI adjustment of the traits in the population using linear regression. For each trait we provide the p-values of testing heterogeneity for the two tissues after WHRadjBMI adjustment. For each trait, tissues which appear to be significant (signif tissue) after WHRadjBMI adjustment are provided. To measure the primary tissue-specific relative change of a trait we calculate the following: 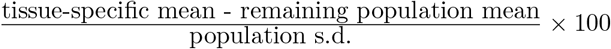, where the tissue-specific mean is computed based only on the individuals classified as the corresponding tissue-specific subtype of WHRadjBMI. The same measure is calculated for the trait residual obtained after adjusting for WHRadjBMI in the population to quantify the tissue-specific relative change of the trait after WHRadjBMI adjustment (adjusted).

| Trait | AS P | MS P | signif tissue | Tissue-specific relative change |  |  |  |
| --- | --- | --- | --- | --- | --- | --- | --- |
|  |  |  |  | AS |  | MS |  |
|  |  |  |  | adjusted | primary | adjusted | primary |
| Non cancer illness code self reported | 1.66E-231 | 7.82E-110 | both | 0.41 | 1.46 | -1.36 | -2.15 |
| Non cancer illness year age first occurred | 5.06E-93 | 1.00E-26 | both | -5.69 | -1.45 | -4.67 | -7.83 |
| Number of self reported non cancer illnesses | 1.94E-87 | 6.28E-13 | both | 15.63 | 11.79 | 13.14 | 15.63 |
| Standing height | 1.81E-08 | 5.60E-178 | both | 3.89 | -21.63 | -36.75 | -14.98 |
| Sitting height | 8.24E-06 | 1.99E-160 | both | 3.22 | -15.58 | -33.73 | -18.68 |
| Body mass index | 6.65E-147 | 4.13E-42 | both | 40.01 | 39.22 | 21.98 | 21.86 |
| Weight | 1.09E-132 | 9.16E-08 | both | 35.38 | 19.36 | 0.19 | 9.81 |
| Waist circumference | 6.93E-113 | 2.32E-18 | both | 35.87 | 3.68 | 15.89 | 31.76 |
| Haemoglobin concentration | 1.06E-22 | 1.61E-98 | both | 7.97 | -16.95 | -27.44 | -7.94 |
| Haematocrit percentage | 3.37E-27 | 7.59E-75 | both | 9.23 | -14.40 | -23.49 | -5.65 |
| Red blood cell erythrocyte count | 1.14E-23 | 6.46E-59 | both | 8.78 | -11.66 | -20.76 | -5.71 |
| High light scatter reticulocyte count | 2.01E-38 | 5.32E-07 | both | 12.12 | 3.64 | 6.41 | 11.99 |
| Red blood cell erythrocyte distribution width | 1.28E-31 | 5.61E-17 | both | 10.05 | 11.09 | 10.83 | 10.08 |
| Immature reticulocyte fraction | 3.84E-31 | 1.12E-16 | both | 11.45 | 8.25 | 10.55 | 12.68 |
| High light scatter reticulocyte percentage | 5.11E-30 | 5.07E-16 | both | 6.84 | 3.42 | 8.27 | 10.43 |
| White blood cell leukocyte count | 7.27E-24 | 3.21E-17 | both | 9.32 | 5.10 | 8.81 | 11.61 |
| Neutrophill count | 7.85E-24 | 1.22E-12 | both | 10.13 | 5.68 | 9.32 | 12.25 |
| Basophill count | 3.48E-11 | 1.37E-23 | both | 1.84 | 2.28 | 5.89 | 5.58 |
| Monocyte count | 4.62E-18 | 1.92E-06 | both | 6.49 | -2.36 | -2.72 | 3.28 |
| Mean corpuscular haemoglobin concentration | 3.22E-05 | 2.08E-17 | both | -2.73 | -7.16 | -9.84 | -6.78 |
| Reticulocyte percentage | 4.03E-17 | 2.22E-10 | both | 4.94 | 1.30 | 6.24 | 8.64 |
| Number of treatments medications taken | 3.89E-81 | 2.33E-27 | both | 15.67 | 12.53 | 17.78 | 19.77 |
| Townsend deprivation index at recruitment | 1.43E-19 | 2.17E-12 | both | 7.92 | 5.76 | 9.42 | 10.87 |
| Creatinine enzymatic in urine | 1.27E-27 | 1.42E-27 | both | 8.87 | -3.13 | -11.60 | -3.35 |
| Reticulocyte count | 1.24E-24 | 0.02 | adipose | 6.39 | -0.24 | 2.40 | 6.86 |
| Age completed full time education | 3.06E-20 | 0.04 | adipose | -4.91 | -3.10 | -5.79 | -7.13 |
| Platelet crit | 0.01 | 3.06E-35 | muscle | -1.96 | 7.53 | 14.62 | 7.93 |
| Monocyte percentage | 0.29 | 5.34E-32 | muscle | 1.21 | -6.11 | -9.67 | -4.58 |
| Platelet count | 0.002 | 1.37E-21 | muscle | -2.26 | 4.84 | 11.42 | 6.49 |
| Lymphocyte count | 0.006 | 6.57E-17 | muscle | 2.35 | 2.59 | 4.72 | 4.58 |
| Mean corpuscular haemoglobin | 0.02 | 1.24E-07 | muscle | -2.05 | -7.16 | -6.43 | -2.91 |
| Neuroticism score | 0.90 | 5.28E-15 | muscle | 0.24 | 4.48 | 10.63 | 7.91 |

**Table S15:** Magnitude of relative change of non-BMI quantitative traits between a tissue-specific subtype group of individuals for BMI and the corresponding group of BMI-matched individuals randomly selected from the population. The magnitude of BMI-matched tissue-specific relative change of a trait is calculated as the following: 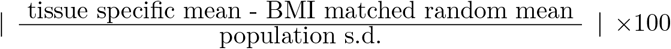, where BMI-matched random mean is the trait mean computed only in the BMI-matched (with the individuals belonging to the corresponding tissue-specific subtype group of BMI) random individuals selected from the population. The tissue-specific mean is computed only in the individuals with the corresponding tissue-specific subtype of BMI. We also provide the magnitude of primary tissue-specific relative change (before BMI-matching) of each trait which is measured by: 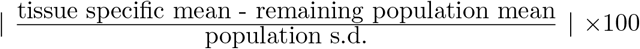. These traits were found to be differentially distributed between at least one of adipose and brain specific group of individuals and the remaining population after BMI adjustment (Table S13).

| Trait | BMI matched adip change |  | adip change | BMI matched brain change |  | brain change |
| --- | --- | --- | --- | --- | --- | --- |
|  | mean | sd | primary | mean | sd | primary |
| Haemoglobin concentration | 0.78 | 0.58 | 14.19 | 3.50 | 0.77 | 20.23 |
| Mean sphered cell volume | 1.68 | 0.87 | 0.23 | 2.44 | 0.79 | 9.89 |
| Mean reticulocyte volume | 1.93 | 0.89 | 2.28 | 2.04 | 0.79 | 5.18 |
| Number of self reported non cancer illnesses | 2.91 | 0.92 | 21.14 | 0.90 | 0.64 | 6.71 |
| Mean corpuscular volume | 1.05 | 0.72 | 5.86 | 2.69 | 0.83 | 3.49 |
| Interpolated Age of participant when non cancer illness first diagnosed | 0.81 | 0.57 | 2.43 | 2.89 | 0.92 | 4.28 |
| Haematocrit percentage | 0.71 | 0.53 | 11.54 | 2.83 | 0.78 | 18.18 |
| Standing height | 0.72 | 0.52 | 15.57 | 2.74 | 0.74 | 13.43 |
| Number of treatments medications taken | 2.60 | 0.92 | 24.52 | 0.81 | 0.60 | 9.68 |
| Mean corpuscular haemoglobin concentration | 0.87 | 0.68 | 7.86 | 2.49 | 0.82 | 5.94 |
| Red blood cell erythrocyte distribution width | 2.31 | 0.92 | 16.74 | 0.91 | 0.66 | 10.18 |
| Neuroticism score | 1.73 | 0.89 | 10.86 | 1.24 | 0.74 | 9.02 |
| Non cancer illness year age first occurred | 1.44 | 0.79 | 8.27 | 1.39 | 0.80 | 0.15 |
| Non cancer illness code self reported | 1.69 | 0.85 | 3.95 | 0.91 | 0.64 | 0.92 |
| Red blood cell erythrocyte count | 0.86 | 0.61 | 7.54 | 1.43 | 0.74 | 18.61 |
| Birth weight | 1.43 | 0.95 | 3.70 | 0.84 | 0.62 | 4.65 |
| Lymphocyte percentage | 1.38 | 0.82 | 6.01 | 0.82 | 0.58 | 5.43 |
| Potassium in urine | 0.71 | 0.53 | 0.42 | 1.45 | 0.78 | 5.49 |
| Sleep duration | 0.75 | 0.58 | 4.26 | 1.39 | 0.75 | 4.20 |
| Eosinophill count | 0.89 | 0.68 | 3.56 | 1.22 | 0.76 | 0.44 |
| Platelet crit | 1.10 | 0.75 | 8.44 | 1.00 | 0.68 | 4.50 |
| Neutrophill count | 0.98 | 0.72 | 15.43 | 1.10 | 0.73 | 7.61 |
| Sitting height | 0.86 | 0.63 | 9.42 | 1.10 | 0.67 | 12.79 |
| Waist circumference | 0.49 | 0.35 | 39.90 | 1.46 | 0.49 | 8.71 |
| Eosinophill percentage | 0.72 | 0.55 | 1.46 | 1.19 | 0.74 | 2.35 |
| High light scatter reticulocyte percentage | 1.10 | 1.06 | 12.78 | 0.76 | 0.72 | 1.00 |
| Sodium in urine | 0.79 | 0.58 | 7.97 | 1.06 | 0.66 | 3.55 |
| Reticulocyte count | 0.73 | 0.55 | 8.54 | 1.09 | 0.63 | 5.06 |
| Monocyte count | 1.03 | 0.71 | 5.21 | 0.77 | 0.57 | 2.42 |
| Neutrophill percentage | 1.07 | 0.75 | 6.58 | 0.70 | 0.51 | 6.61 |
| Platelet count | 1.04 | 0.71 | 6.48 | 0.70 | 0.52 | 4.13 |
| Creatinine enzymatic in urine | 0.91 | 0.68 | 4.89 | 0.84 | 0.60 | 6.85 |
| Monocyte percentage | 0.97 | 0.70 | 4.09 | 0.76 | 0.55 | 5.23 |
| Platelet distribution width | 0.95 | 0.66 | 0.44 | 0.75 | 0.54 | 4.24 |
| Reticulocyte percentage | 0.80 | 0.61 | 8.85 | 0.85 | 0.57 | 2.90 |
| Basophill count | 0.83 | 0.64 | 7.50 | 0.82 | 0.60 | 5.30 |
| Basophill percentage | 0.84 | 0.65 | 4.48 | 0.79 | 0.59 | 4.77 |
| Townsend deprivation index at recruitment | 0.95 | 0.67 | 15.63 | 0.67 | 0.52 | 11.34 |
| Weight | 1.23 | 0.54 | 45.05 | 0.37 | 0.28 | 4.81 |
| White blood cell leukocyte count | 0.69 | 0.52 | 13.43 | 0.81 | 0.61 | 4.67 |
| High light scatter reticulocyte count | 0.81 | 0.61 | 19.22 | 0.62 | 0.48 | 3.31 |

**Table S16:** Magnitude of relative change of non-WHR quantitative traits between a tissue-specific subtype group of individuals for WHRadjBMI and the corresponding group of WHRadjBMI-matched individuals randomly selected from the population. The magnitude of WHRadjBMI-matched tissue-specific relative change of a trait is calculated as the following: 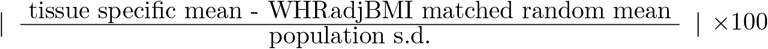, where WHRadjBMI-matched random mean is the trait mean computed only in the WHRadjBMI-matched (with the individuals belonging to the corresponding tissue-specific subtype group of WHRadjBMI) random individuals selected from the population. The tissue-specific mean is computed only in the individuals with the corresponding tissue-specific subtype of WHRadjBMI. We also provide the magnitude of primary tissue-specific relative change (before WHRadjBMI-matching) of each trait which is measured by: 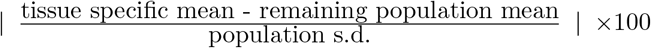. These traits were found to be differentially distributed between at least one of adipose subcutaneous (AS) and muscle skeletal (MS) specific group of individuals and the remaining population after WHRadjBMI adjustment (Table S14).

| Trait | WHRadjBMI matched AS change |  | AS change<br>primary | WHRadjBMI matched MS change |  | MS change<br>primary |
| --- | --- | --- | --- | --- | --- | --- |
|  | mean | sd |  | mean | sd |  |
| Sitting height | 3.56 | 0.82 | 15.58 | 30.18 | 1.03 | 18.68 |
| Standing height | 2.85 | 0.73 | 21.63 | 30.51 | 0.94 | 14.98 |
| Haemoglobin concentration | 0.89 | 0.61 | 16.95 | 24.21 | 1.06 | 7.94 |
| Haematocrit percentage | 0.61 | 0.46 | 14.40 | 21.90 | 1.08 | 5.65 |
| Red blood cell erythrocyte count | 0.76 | 0.54 | 11.66 | 20.14 | 1.11 | 5.71 |
| Weight | 3.38 | 0.91 | 19.36 | 16.41 | 1.10 | 9.81 |
| Creatinine enzymatic in urine | 0.66 | 0.50 | 3.13 | 14.27 | 1.16 | 3.35 |
| Platelet crit | 0.92 | 0.64 | 7.53 | 12.96 | 1.12 | 7.93 |
| Number of treatments<br>medications taken | 5.19 | 0.92 | 12.53 | 8.31 | 1.16 | 19.77 |
| Platelet count | 0.74 | 0.55 | 4.84 | 10.78 | 1.14 | 6.49 |
| Number of self reported<br>non cancer illnesses | 6.12 | 0.91 | 11.79 | 5.16 | 1.20 | 15.63 |
| Monocyte percentage | 1.23 | 0.77 | 6.11 | 9.91 | 1.19 | 4.58 |
| Neuroticism score | 0.78 | 0.58 | 4.48 | 8.73 | 1.26 | 7.91 |
| Mean corpuscular<br>haemoglobin concentration | 1.07 | 0.75 | 7.16 | 7.36 | 1.15 | 6.78 |
| Red blood cell erythrocyte<br>distribution width | 3.14 | 0.92 | 11.09 | 4.83 | 1.15 | 10.08 |
| Monocyte count | 0.72 | 0.52 | 2.36 | 6.96 | 1.24 | 3.28 |
| Body mass index | 6.07 | 1.00 | 39.22 | 0.97 | 0.72 | 21.86 |
| Townsend deprivation<br>index at recruitment | 2.32 | 0.84 | 5.76 | 4.52 | 1.24 | 10.87 |
| Immature reticulocyte fraction | 2.44 | 0.92 | 8.25 | 3.81 | 1.19 | 12.68 |
| Waist circumference | 4.10 | 0.85 | 3.68 | 1.86 | 0.94 | 31.76 |
| Lymphocyte count | 1.81 | 0.78 | 2.59 | 3.96 | 1.25 | 4.58 |
| Age completed full time education | 2.17 | 1.06 | 3.10 | 3.35 | 1.35 | 7.13 |
| Mean corpuscular haemoglobin | 0.72 | 0.57 | 7.16 | 4.76 | 1.17 | 2.91 |
| High light scatter<br>reticulocyte percentage | 1.21 | 0.61 | 3.42 | 4.03 | 1.07 | 10.43 |
| White blood cell leukocyte count | 1.84 | 0.81 | 5.10 | 2.83 | 1.19 | 11.61 |
| Basophill count | 0.75 | 0.56 | 2.28 | 3.71 | 1.21 | 5.58 |
| Reticulocyte percentage | 0.76 | 0.55 | 1.30 | 3.25 | 1.12 | 8.64 |
| Neutrophill count | 1.39 | 0.79 | 5.68 | 2.31 | 1.12 | 12.25 |
| Non cancer illness code self reported | 1.57 | 0.88 | 1.46 | 1.07 | 0.83 | 2.15 |
| High light scatter reticulocyte count | 1.60 | 0.82 | 3.64 | 0.97 | 0.69 | 11.99 |
| Non cancer illness year<br>age first occurred | 0.82 | 0.59 | 1.45 | 1.24 | 0.90 | 7.83 |
| Reticulocyte count | 0.72 | 0.54 | 0.24 | 1.16 | 0.82 | 6.86 |

**Table S17:** Summary of results from the analyses performed to characterize the tissue-specificity of phenotypic characteristics of individuals assigned to a tissue-specific subtype of BMI. Central tendency measures of the artificial tissue-specific mean of a quantitative trait (computed only in the individuals classified into artificial tissue-specific subtype of BMI) across 500 artificial tissue-specific subtype groups identified based on 500 sets of artificial tissue-specific eQTLs (obtained by random exchange of eQTLs between the sets of adipose and brain specific eQTLs) are provided. These quantitative traits were found primarily heterogeneous between at least one of real adipose and brain specific subtype groups of individuals and the remaining population (Table S8). Here adipose P denotes the p-value of testing whether the artificial adipose tissue-specific trait mean across the artificial adipose tissue-specific subtype groups is different from the original adipose tissue-specific trait mean. Similarly Brain P is defined. The real adipose and brain tissue-specific mean and population mean of the traits are also provided.

| Trait | real<br>adip mean | shuffled<br>adip mean sd | adipose P | real<br>brain mean | shuffled<br>brain mean sd | brain P | popln<br>mean |
| --- | --- | --- | --- | --- | --- | --- | --- |
| Body mass index | 30.39 | 29.4(1.102) | 6.25E-142 | 27.60 | 28.58(1.133) | 9.76E-130 | 27.40 |
| Waist circumference | 95.50 | 93.25(2.455) | 1.00E-146 | 89.20 | 91.36(2.51) | 2.53E-128 | 90.33 |
| Weight | 85.20 | 82.53(2.982) | 4.36E-140 | 77.58 | 80.27(3.059) | 2.32E-135 | 78.31 |
| Standing height | 167.45 | 167.57(0.144) | 1.81E-127 | 167.65 | 167.62(0.152) | 1.41E-08 | 168.84 |
| Sitting height | 88.97 | 88.95(0.054) | 1.58E-32 | 88.82 | 88.89(0.054) | 0 | 89.41 |
| Haemoglobin concentration | 14.04 | 14.03(0.029) | 1.86E-62 | 13.97 | 13.99(0.028) | 5.19E-112 | 14.21 |
| Haematocrit percentage | 40.76 | 40.72(0.083) | 1.42E-60 | 40.54 | 40.63(0.079) | 6.79E-227 | 41.16 |
| Red blood cell erythrocyte count | 4.48 | 4.47(0.016) | 1.67E-81 | 4.44 | 4.46(0.016) | 5.71E-226 | 4.51 |
| High light scatter reticulocyte percentage | 0.44 | 0.43(0.019) | 3.60E-120 | 0.40 | 0.41(0.02) | 2.33E-117 | 0.40 |
| Red blood cell erythrocyte distribution width | 13.62 | 13.6(0.018) | 3.32E-314 | 13.56 | 13.59(0.019) | 0 | 13.47 |
| White blood cell leukocyte count | 7.15 | 7.1(0.076) | 1.07E-88 | 6.98 | 7.05(0.079) | 1.23E-131 | 6.89 |
| High light scatter reticulocyte count | 0.02 | 0.02(0.001) | 1.16E-157 | 0.02 | 0.02(0.001) | 5.81E-99 | 0.02 |
| Neutrophil count | 4.45 | 4.41(0.046) | 2.68E-134 | 4.35 | 4.39(0.048) | 5.00E-112 | 4.24 |
| Reticulocyte count | 0.06 | 0.06(0.002) | 2.21E-172 | 0.06 | 0.06(0.002) | 5.46E-56 | 0.06 |
| Reticulocyte percentage | 1.42 | 1.38(0.043) | 3.86E-167 | 1.32 | 1.35(0.045) | 5.01E-57 | 1.35 |
| Mean corpuscular haemoglobin concentration | 34.46 | 34.46(0.01) | 1.84E-08 | 34.48 | 34.46(0.01) | 0 | 34.54 |
| Platelet crit | 0.24 | 0.24(0.001) | 5.28E-140 | 0.23 | 0.24(0.001) | 3.18E-64 | 0.23 |
| Basophil count | 0.04 | 0.04(0.001) | 9.52E-77 | 0.04 | 0.04(0.001) | 1.08E-163 | 0.03 |
| Monocyte percentage | 6.99 | 6.98(0.026) | 5.52E-78 | 6.97 | 6.98(0.026) | 1.07E-26 | 7.10 |
| Neutrophil percentage | 61.67 | 61.68(0.077) | 0.0004 | 61.67 | 61.72(0.079) | 1.20E-61 | 61.14 |
| Mean corpuscular volume | 91.09 | 91.2(0.148) | 3.37E-89 | 91.49 | 91.32(0.155) | 1.56E-210 | 91.34 |
| Platelet count | 256.95 | 256.41(0.845) | 2.72E-71 | 255.58 | 256.01(0.928) | 6.14E-39 | 253.21 |
| Lymphocyte percentage | 28.20 | 28.22(0.067) | 4.46E-07 | 28.25 | 28.19(0.07) | 2.15E-114 | 28.63 |
| Monocyte count | 0.49 | 0.48(0.005) | 7.09E-222 | 0.47 | 0.48(0.005) | 3.00E-225 | 0.48 |
| Basophil percentage | 0.59 | 0.59(0.006) | 3.44E-07 | 0.60 | 0.6(0.006) | 0.02 | 0.57 |
| Number of treatments medications taken | 3.08 | 2.9(0.132) | 0 | 2.70 | 2.81(0.135) | 3.64E-120 | 2.45 |
| Number of self reported non cancer illnesses | 2.24 | 2.12(0.087) | 0 | 1.98 | 2.07(0.088) | 4.23E-156 | 1.87 |
| Interpolated Age of participant when<br>non cancer illness first diagnosed | 41.89 | 41.65(0.256) | 8.57E-144 | 41.56 | 41.46(0.263) | 6.73E-26 | 42.32 |
| Townsend deprivation index at recruitment | -1.13 | -1.17(0.055) | 9.18E-71 | -1.26 | -1.18(0.057) | 1.54E-316 | -1.58 |
| Neuroticism score | 4.44 | 4.4(0.032) | 2.37E-322 | 4.38 | 4.41(0.033) | 1.51E-120 | 4.10 |
| Creatinine enzymatic in urine | 9077.69 | 8885.47(252.885) | 3.80E-101 | 8429.45 | 8677.5(254.514) | 5.26E-165 | 8806.93 |
| Sodium in urine | 79.56 | 77.86(1.986) | 7.51E-127 | 74.72 | 76.34(2.001) | 1.92E-114 | 76.21 |
| Sleep duration | 7.08 | 7.08(0.014) | 7.10E-21 | 7.08 | 7.08(0.014) | 2.82E-26 | 7.13 |
| Immature reticulocyte fraction | 0.30 | 0.3(0.004) | 2.20E-127 | 0.29 | 0.29(0.004) | 4.98E-141 | 0.29 |
| Mean corpuscular haemoglobin | 31.39 | 31.43(0.052) | 7.04E-99 | 31.55 | 31.46(0.055) | 0 | 31.55 |
| Lymphocyte count | 1.98 | 1.98(0.024) | 1.06E-16 | 1.93 | 1.96(0.025) | 1.32E-179 | 1.95 |
| Eosinophil count | 0.18 | 0.18(0.003) | 3.41E-43 | 0.17 | 0.18(0.003) | 6.94E-66 | 0.17 |
| Non cancer illness code self reported | 2405.55 | 2696.21(140.196) | 0 | 2779.19 | 2747.36(146.249) | 9.77E-10 | 2891.97 |
| Non cancer illness year age first occurred | 628.73 | 646.05(28.147) | 6.08E-67 | 703.82 | 665.47(29.619) | 1.36E-289 | 702.47 |
| Mean spheroid cell volume | 82.86 | 83(0.201) | 3.20E-90 | 83.37 | 83.18(0.206) | 1.21E-147 | 82.87 |
| Mean reticulocyte volume | 106.00 | 106.02(0.125) | 3.58E-05 | 106.22 | 106.13(0.13) | 3.50E-83 | 105.83 |
| Eosinophil percentage | 2.53 | 2.53(0.018) | 6.55E-10 | 2.52 | 2.52(0.019) | 0.71 | 2.56 |
| Platelet distribution width | 16.49 | 16.48(0.007) | 0 | 16.47 | 16.48(0.007) | 2.24E-205 | 16.49 |
| Potassium in urine | 63.61 | 62.84(0.851) | 1.64E-141 | 61.68 | 62.13(0.858) | 8.05E-48 | 63.47 |
| Birth weight | 3.30 | 3.3(0.009) | 0.0001 | 3.30 | 3.3(0.009) | 8.54E-06 | 3.33 |

**Table S18:** Summary of results from the analyses performed to characterize the tissue-specificity of phenotypic characteristics of the individuals assigned to a tissue-specific subtype of WHRadjBMI. Central tendency measures of the artificial tissue-specific mean of a quantitative trait (computed only in the individuals classified into artificial tissue-specific subtype of WHRadjBMI) across 500 artificial tissue-specific subtype groups identified based on 500 sets of artificial tissue-specific eQTLs (obtained by random exchange of eQTLs between the sets of adipose subcutaneous (AS) and muscle skeletal (MS) specific eQTLs) are provided. These quantitative traits were found primarily heterogeneous between at least one of real AS and MS specific subtype groups of individuals and the remaining population (Table S9). Here AS P denotes the p-value of testing whether the artificial AS tissue-specific trait mean across the artificial AS tissue-specific subtype groups is different from the original AS tissue-specific trait mean. Similarly MS P is defined. The real AS and MS tissue-specific mean and overall population mean of the traits are also provided.

| Trait | real<br>AS mean | shuffled<br>AS mean sd | AS P | real<br>MS mean | shuffled<br>MS mean sd | MS P | popln<br>mean |
| --- | --- | --- | --- | --- | --- | --- | --- |
| WHRadjBMI | -0.04 | -0.01(0.018) | 0 | 0.03 | -0.01(0.018) | 0 | 0.00 |
| WHR | 0.85 | 0.88(0.017) | 0 | 0.91 | 0.88(0.017) | 0 | 0.87 |
| Body mass index | 29.19 | 28.81(0.232) | 0 | 28.41 | 28.76(0.223) | 0 | 27.39 |
| Standing height | 166.91 | 167.14(0.152) | 0 | 167.48 | 167.17(0.15) | 0 | 168.84 |
| Sitting height | 88.68 | 88.6(0.105) | 7.55E-117 | 88.53 | 88.62(0.102) | 2.92E-129 | 89.41 |
| Weight | 81.27 | 80.53(0.513) | 0 | 79.83 | 80.42(0.488) | 2.76E-258 | 78.31 |
| Haemoglobin concentration | 14.01 | 14.06(0.029) | 0 | 14.12 | 14.06(0.028) | 0 | 14.21 |
| Haematocrit percentage | 40.67 | 40.78(0.075) | 0 | 40.96 | 40.78(0.074) | 0 | 41.16 |
| Red blood cell erythrocyte<br>distribution width | 13.57 | 13.56(0.012) | 4.15E-198 | 13.56 | 13.55(0.011) | 2.64E-86 | 13.47 |
| Red blood cell erythrocyte count | 4.47 | 4.48(0.007) | 2.12E-289 | 4.49 | 4.48(0.007) | 0 | 4.51 |
| High light scatter reticulocyte percentage | 0.41 | 0.42(0.007) | 0 | 0.43 | 0.42(0.007) | 0 | 0.40 |
| White blood cell leukocyte count | 6.99 | 7.05(0.058) | 1.29E-220 | 7.12 | 7.04(0.057) | 2.30E-300 | 6.89 |
| Immature reticulocyte fraction | 0.29 | 0.3(0.001) | 1.31E-202 | 0.30 | 0.3(0.001) | 1.62E-205 | 0.29 |
| Neutrophil count | 4.32 | 4.37(0.039) | 7.42E-258 | 4.41 | 4.36(0.038) | 7.97E-298 | 4.24 |
| Mean corpuscular haemoglobin<br>concentration | 34.47 | 34.48(0.014) | 3.47E-139 | 34.47 | 34.48(0.013) | 8.32E-125 | 34.54 |
| Monocyte percentage | 6.94 | 6.98(0.031) | 1.34E-285 | 6.98 | 6.99(0.03) | 3.47E-08 | 7.10 |
| Platelet crit | 0.24 | 0.24(0.001) | 1.54E-98 | 0.24 | 0.24(0.001) | 3.98E-12 | 0.23 |
| Lymphocyte count | 1.98 | 1.98(0.017) | 6.63E-15 | 2.00 | 1.98(0.016) | 1.18E-259 | 1.95 |
| Platelet count | 256.00 | 257.04(0.987) | 8.71E-192 | 257.00 | 256.89(0.957) | 0.001 | 253.21 |
| Number of treatments medications taken | 2.77 | 2.86(0.068) | 1.89E-288 | 2.97 | 2.85(0.067) | 0 | 2.45 |
| Number of self reported non cancer illnesses | 2.08 | 2.12(0.045) | 4.48E-154 | 2.15 | 2.11(0.044) | 1.61E-145 | 1.86 |
| Townsend deprivation index at recruitment | -1.41 | -1.33(0.06) | 0 | -1.26 | -1.34(0.059) | 4.50E-285 | -1.58 |
| Neuroticism score | 4.24 | 4.31(0.058) | 1.24E-223 | 4.35 | 4.3(0.056) | 7.90E-147 | 4.10 |
| Mean corpuscular haemoglobin | 31.43 | 31.46(0.026) | 0 | 31.50 | 31.47(0.026) | 4.64E-271 | 31.55 |
| Monocyte count | 0.47 | 0.48(0.004) | 0 | 0.49 | 0.48(0.005) | 4.75E-203 | 0.48 |
| Mean corpuscular volume | 91.18 | 91.25(0.056) | 3.11E-257 | 91.38 | 91.26(0.055) | 0 | 91.34 |
| Mean platelet thrombocyte volume | 9.36 | 9.35(0.016) | 6.32E-226 | 9.33 | 9.35(0.016) | 5.81E-136 | 9.32 |
| Eosinophil percentage | 2.49 | 2.52(0.018) | 0 | 2.53 | 2.52(0.019) | 3.38E-49 | 2.56 |
| Creatinine enzymatic in urine | 8633.55 | 8672.89(66.169) | 1.54E-62 | 8618.76 | 8670.39(67.216) | 4.07E-103 | 8806.64 |
| Waist circumference | 90.80 | 92.5(0.98) | 0 | 94.50 | 92.39(0.981) | 0 | 90.33 |
| High light scatter reticulocyte count | 0.02 | 0.02(0) | 0 | 0.02 | 0.02(0) | 0 | 0.02 |
| Reticulocyte percentage | 1.36 | 1.39(0.019) | 0 | 1.42 | 1.38(0.019) | 0 | 1.35 |
| Reticulocyte count | 0.06 | 0.06(0.001) | 0 | 0.06 | 0.06(0.001) | 0 | 0.06 |
| Basophil count | 0.03 | 0.04(0.001) | 4.27E-248 | 0.04 | 0.04(0.001) | 0 | 0.03 |
| Non cancer illness code self reported | 3071.24 | 2746.71(158.474) | 0 | 2623.36 | 2747.26(158.328) | 5.81E-107 | 2891.68 |
| Non cancer illness year age first occurred | 689.59 | 655.07(14.516) | 0 | 631.65 | 655.89(14.155) | 0 | 702.50 |
| Age completed full time education | 16.36 | 16.32(0.044) | 3.26E-150 | 16.25 | 16.33(0.042) | 0 | 16.45 |

**Table S19:**
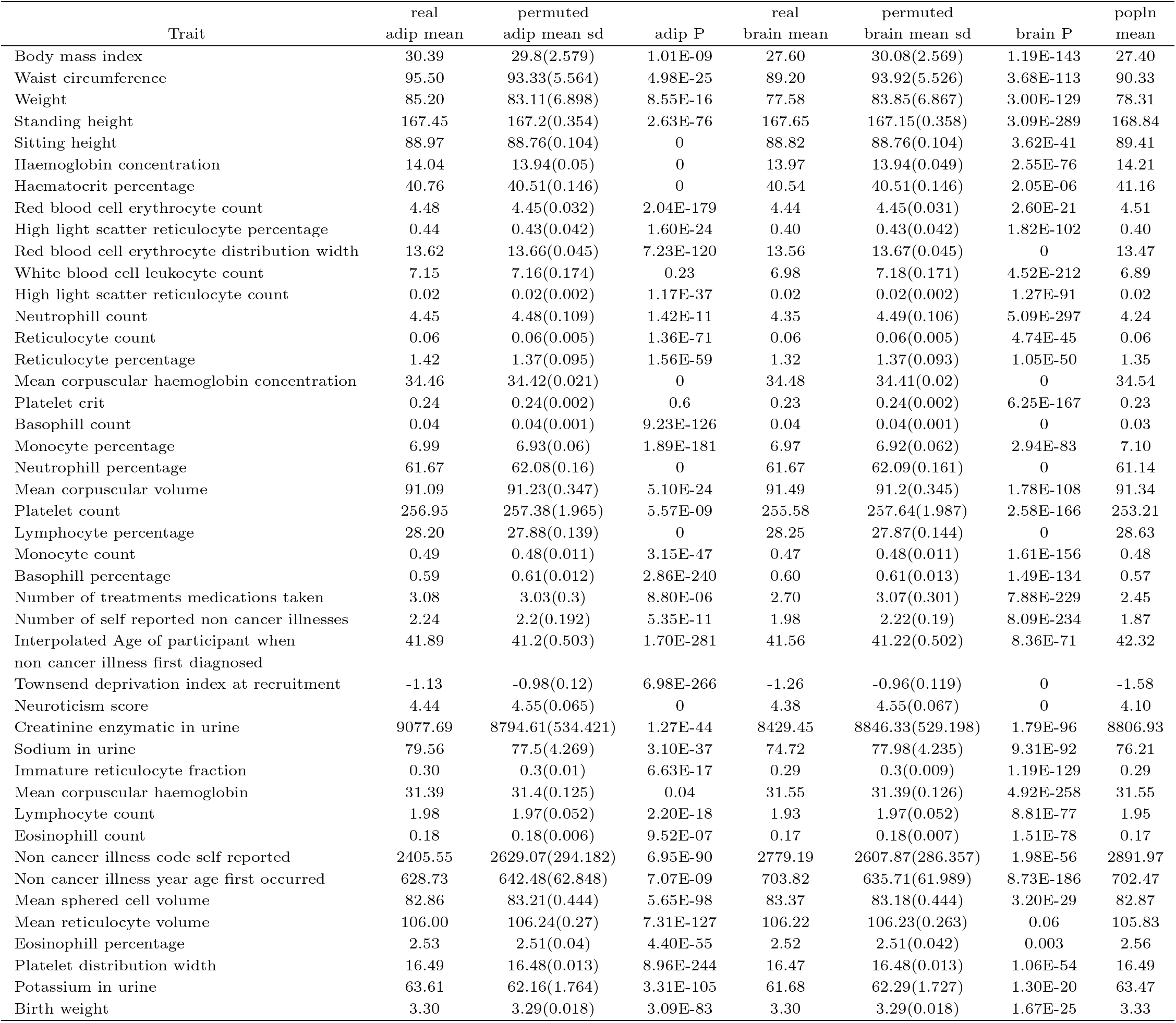
Summary of results on the phenotypic characteristics of the individuals assigned to tissue-specific subtype of BMI identified based on random permutations of the primary phenotype (BMI) across individuals. Central tendency measures of the tissue-specific mean of a quantitative trait (computed only in the individuals classified into tissue-specific subtype detected based on a randomly permuted BMI data) across 500 tissue-specific subtype groups identified based on 500 random permutations of BMI data are provided. These quantitative traits were found primarily heterogeneous between at least one of real adipose and brain specific subtype groups of individuals and the remaining population (Table S8). Here adipose P denotes the p-value of testing whether a trait mean computed only in the random adipose subtype group of individuals (detected based on random permutations of BMI) is different from the original adipose tissue-specific trait mean. Similarly Brain P is defined. The real adipose and brain tissue-specific mean and overall population mean of the traits are also provided.

**Table S20:**
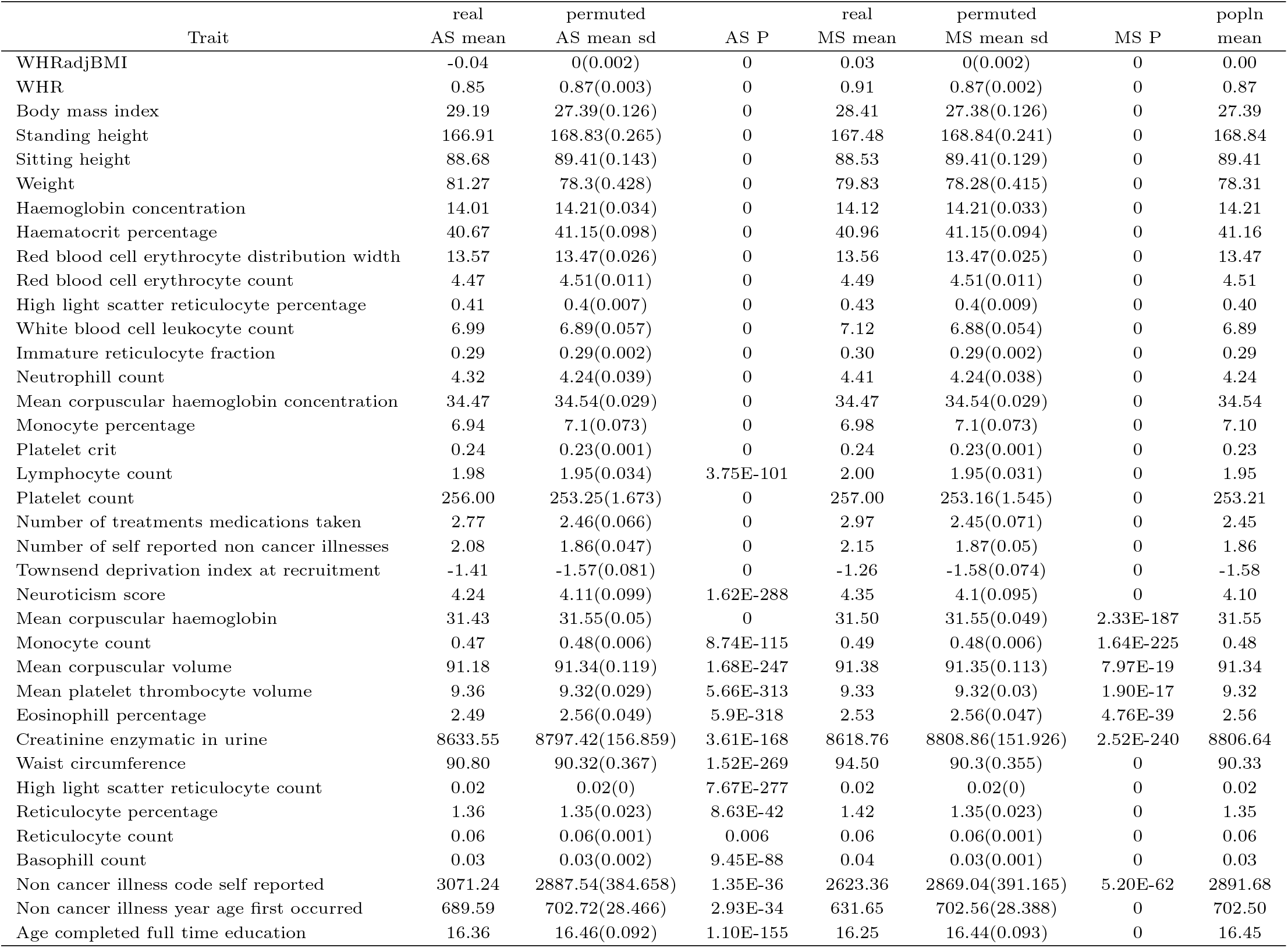
Summary of results on the phenotypic characteristics of the individuals assigned to tissue-specific subtype of WHRadjBMI identified based on random permutations of the primary phenotype (WHRadjBMI) across individuals. Central tendency measures of the tissue-specific mean of a quantitative trait (computed only in the individuals classified into a tissue-specific subtype detected based on a randomly permuted WHRadjBMI data) across 500 tissue-specific subtype groups identified based on 500 random permutations of WHRadjBMI data are provided. These quantitative traits were found primarily heterogeneous between at least one of real adipose subcutaneous (AS) and muscle skeletal (MS) specific subtype groups of individuals and the remaining population (Table S9). Here AS P denotes the p-value of testing whether a trait mean computed only in the random AS-specific subtype group of individuals (detected based on random permutations of WHRadjBMI) is different from the original AS tissue-specific trait mean. Similarly MS P is defined. The real AS and MS tissue-specific mean and overall population mean of the traits are also provided.

## References

[1] Jaana M Leppala, Jarmo Virtamo, Rainer Fogelholm, Demetrius Albanes, and Olli P Heinonen. Different risk factors for different stroke subtypes: association of blood pressure, cholesterol, and antioxidants. Stroke, 30(12):2535–2540, 1999.

[2] BM Schamberger, HC Geiss, MM Ritter, P Schwandt, and KG Parhofer. Influence of ldl apheresis on ldl subtypes in patients with coronary heart disease and severe hyperlipoproteinemia. Journal of lipid research, 41(5):727–733, 2000.

[3] John M Ringman, Alison Goate, Colin L Masters, Nigel J Cairns, Adrian Danek, Neill Graff-Radford, Bernardino Ghetti, John C Morris, Dominantly Inherited Alzheimer Network, et al. Genetic heterogeneity in alzheimer disease and implications for treatment strategies. Current neurology and neuroscience reports, 14(11):499, 2014.

[4] Paul Gibson, Yiai Tong, Giles Robinson, Margaret C Thompson, D Spencer Currle, Christopher Eden, Tanya A Kranenburg, Twala Hogg, Helen Poppleton, Julie Martin, et al. Subtypes of medulloblastoma have distinct developmental origins. Nature, 468(7327):1095, 2010.

[5] Judy H Cho and Marc Feldman. Heterogeneity of autoimmune diseases: pathophysiologic insights from genetics and implications for new therapies. Nature medicine, 21(7):730, 2015.

[6] Javaid Iqbal, Ophira Ginsburg, Paula A Rochon, Ping Sun, and Steven A Narod. Differences in breast cancer stage at diagnosis and cancer-specific survival by race and ethnicity in the united states. Jama, 313(2):165–173, 2015.

[7] Roger L Milne, Karoline B Kuchenbaecker, Kyriaki Michailidou, Jonathan Beesley, Siddhartha Kar, Sara Lindstrom, Shirley Hui, Audrey Lemaçon, Penny Soucy, Joe Dennis, et al. Identification of ten variants associated with risk of estrogen-receptor-negative breast cancer. Nature genetics, 49(12):1767, 2017.

[8] Montserrat Garcia-Closas, Fergus J Couch, Sara Lindstrom, Kyriaki Michailidou, Marjanka K Schmidt, Mark N Brook, Nick Orr, Suhn Kyong Rhie, Elio Riboli, Heather S Feigelson, et al. Genome-wide association studies identify four er negative-specific breast cancer risk loci. Nature genetics, 45(4):392, 2013.

[9] Mark Zimmerman, Theresa A Morgan, and Kasey Stanton. The severity of psychiatric disorders. World Psychiatry, 17(3):258–275, 2018.

[10] Raphael Bernier, Christelle Golzio, Bo Xiong, Holly A Stessman, Bradley P Coe, Osnat Penn, Kali Witherspoon, Jennifer Gerdts, Carl Baker, Anneke T Vulto-van Silfhout, et al. Disruptive chd8 mutations define a subtype of autism early in development. Cell, 158(2):263–276, 2014.

[11] Shafali S Jeste and Daniel H Geschwind. Disentangling the heterogeneity of autism spectrum disorder through genetic findings. Nature Reviews Neurology, 10(2):74, 2014.

[12] Holly A Stessman, Raphael Bernier, and Evan E Eichler. A genotype-first approach to defining the subtypes of a complex disease. Cell, 156(5):872–877, 2014.

[13] Miriam S Udler, Jaegil Kim, Marcin von Grotthuss, Sílvia Bonàs-Guarch, Joanne B Cole, Joshua Chiou, Michael Boehnke, Markku Laakso, Gil Atzmon, Benjamin Glaser, et al. Type 2 diabetes genetic loci informed by multi-trait associations point to disease mechanisms and subtypes: A soft clustering analysis. PLoS medicine, 15(9):e1002654, 2018.

[14] Thomas W Winkler, Felix Gunther, Simon Höllerer, Martina Zimmermann, Ruth JF Loos, Zoltán Kutalik, and Iris M Heid. A joint view on genetic variants for adiposity differentiates subtypes with distinct metabolic implications. Nature communications, 9(1):1946, 2018.

[15] Andy Dahl, Na Cai, Arthur Ko, Markku Laakso, Paivi Pajukanta, Jonathan Flint, and Noah Zaitlen. Reverse gwas: Using genetics to identify and model phenotypic subtypes. bioRxiv, page 446–492, 2018.

[16] GTEx Consortium et al. The genotype-tissue expression (gtex) pilot analysis: multitissue gene regulation in humans. Science, 348(6235):648–660, 2015.

[17] Alexander Gusev, Arthur Ko, Huwenbo Shi, Gaurav Bhatia, Wonil Chung, Brenda WJH Penninx, Rick Jansen, Eco JC De Geus, Dorret I Boomsma, Fred A Wright, et al. Integrative approaches for large-scale transcriptome-wide association studies. Nature genetics, 48(3):245, 2016.

[18] GTEx Consortium et al. Genetic effects on gene expression across human tissues. Nature, 550(7675):204, 2017.

[19] Jason M Torres, Eric R Gamazon, Esteban J Parra, Jennifer E Below, Adan Valladares-Salgado, Niels Wacher, Miguel Cruz, Craig L Hanis, and Nancy J Cox. Cross-tissue and tissue-specific eqtls: partitioning the heritability of a complex trait. The American Journal of Human Genetics, 95(5):521–534, 2014.

[20] Tune H Pers, Juha M Karjalainen, Yingleong Chan, Harm-Jan Westra, Andrew R Wood, Jian Yang, Julian C Lui, Sailaja Vedantam, Stefan Gustafsson, Tonu Esko, et al. Biological interpretation of genome-wide association studies using predicted gene functions. Nature communications, 6(1):1–9, 2015.

[21] Ruo-Han Hao, Tie-Lin Yang, Yu Rong, Shi Yao, Shan-Shan Dong, Hao Chen, and Yan Guo. Gene expression profiles indicate tissue-specific obesity regulation changes and strong obesity relevant tissues. International Journal of Obesity, 2017.

[22] Halit Ongen, Andrew A Brown, Olivier Delaneau, Nikolaos I Panousis, Alexandra C Nica, Emmanouil T Dermitzakis, GTEx Consortium, et al. Estimating the causal tissues for complex traits and diseases. Nature genetics, 49(12):1676, 2017.

[23] Hilary K Finucane, Yakir A Reshef, Verneri Anttila, Kamil Slowikowski, Alexander Gusev, Andrea Byrnes, Steven Gazal, Po-Ru Loh, Caleb Lareau, Noam Shoresh, et al. Heritability enrichment of specifically expressed genes identifies disease-relevant tissues and cell types. Nature genetics, 50(4):621, 2018.

[24] Diego Calderon, Anand Bhaskar, David A Knowles, David Golan, Towfique Raj, Audrey Q Fu, and Jonathan K Pritchard. Inferring relevant cell types for complex traits by using single-cell gene expression. The American Journal of Human Genetics, 101(5):686–699, 2017.

[25] Kamil Slowikowski, Xinli Hu, and Soumya Raychaudhuri. Snpsea: an algorithm to identify cell types, tissues and pathways affected by risk loci. Bioinformatics, 30(17):2496–2497, 2014.

[26] Dmitry Shungin, Thomas W Winkler, Damien C Croteau-Chonka, Teresa Ferreira, Adam E Locke, Reedik Möagi, Rona J Strawbridge, Tune H Pers, Krista Fischer, Anne E Justice, et al. New genetic loci link adipose and insulin biology to body fat distribution. Nature, 518(7538):187–196, 2015.

[27] Adam E Locke, Bratati Kahali, Sonja I Berndt, Anne E Justice, Tune H Pers, Felix R Day, Corey Powell, Sailaja Vedantam, Martin L Buchkovich, Jian Yang, et al. Genetic studies of body mass index yield new insights for obesity biology. Nature, 518(7538):197–206, 2015.

[28] Nicholas Mancuso, Simon Gayther, Alexander Gusev, Wei Zheng, Kathryn L Penney, Zsofia Kote-Jarai, Rosalind Eeles, Matthew Freedman, Christopher Haiman, and Bogdan Pasaniuc. Large-scale transcriptome-wide association study identifies new prostate cancer risk regions. Nature communications, 9(1):1—11, 2018.

[29] Yiming Hu, Mo Li, Qiongshi Lu, Haoyi Weng, Jiawei Wang, Seyedeh M Zekavat, Zhaolong Yu, Boyang Li, Jianlei Gu, Sydney Muchnik, et al. A statistical framework for cross-tissue transcriptome-wide association analysis. Nature genetics, 51(3):568–576, 2019.

[30] Alvaro N Barbeira, Milton Pividori, Jiamao Zheng, Heather E Wheeler, Dan L Nicolae, and Hae Kyung Im. Integrating predicted transcriptome from multiple tissues improves association detection. PLoS genetics, 15(1):e1007889, 2019.

[31] Liangying Yin, Carlos KL Chau, Pak-Chung Sham, and Hon-Cheong So. Integrating clinical data and imputed transcriptome from gwas to uncover complex disease subtypes: Applications in psychiatry and cardiology. The American Journal of Human Genetics, 105(6):1193–1212, 2019.

[32] Cathie Sudlow, John Gallacher, Naomi Allen, Valerie Beral, Paul Burton, John Danesh, Paul Downey, Paul Elliott, Jane Green, Martin Landray, et al. Uk biobank: an open access resource for identifying the causes of a wide range of complex diseases of middle and old age. PLoS medicine, 12(3):e1001779, 2015.

[33] Clare Bycroft, Colin Freeman, Desislava Petkova, Gavin Band, Lloyd T Elliott, Kevin Sharp, Allan Motyer, Damjan Vukcevic, Olivier Delaneau, Jared OConnell, et al. The uk biobank resource with deep phenotyping and genomic data. Nature, 562(7726):203, 2018.

[34] Eric R Gamazon, Heather E Wheeler, Kaanan P Shah, Sahar V Mozaffari, Keston Aquino-Michaels, Robert J Carroll, Anne E Eyler, Joshua C Denny, Dan L Nicolae, Nancy J Cox, et al. A gene-based association method for mapping traits using reference transcriptome data. Nature genetics, 47(9):1091, 2015.

[35] Rachel A Murphy, Ilse Reinders, Melissa E Garcia, Gudny Eiriksdottir, Lenore J Launer, Rafn Benedik-tsson, Vilmundur Gudnason, Palmi V Jonsson, and Tamara B Harris. Adipose tissue, muscle, and function: potential mediators of associations between body weight and mortality in older adults with type 2 diabetes. Diabetes Care, 37(12):3213–3219, 2014.

[36] DC Chan, Gerald F Watts, PHR Barrett, and Valerie Burke. Waist circumference, waist-to-hip ratio and body mass index as predictors of adipose tissue compartments in men. Qjm, 96(6):441–447, 2003.

[37] V Van Harmelen, T Skurk, K Röhrig, YM Lee, M Halbleib, I Aprath-Husmann, and H Hauner. Effect of bmi and age on adipose tissue cellularity and differentiation capacity in women. International journal of obesity, 27(8):889–895, 2003.

[38] Yiming Hu, Qiongshi Lu, Ryan Powles, Xinwei Yao, Can Yang, Fang Fang, Xinran Xu, and Hongyu Zhao. Leveraging functional annotations in genetic risk prediction for human complex diseases. PLoS computational biology, 13(6):e1005589, 2017.

[39] Farhad Hormozdiari, Steven Gazal, Bryce van de Geijn, Hilary K Finucane, Chelsea J-T Ju, Po-Ru Loh, Armin Schoech, Yakir Reshef, Xuanyao Liu, Luke OConnor, et al. Leveraging molecular quantitative trait loci to understand the genetic architecture of diseases and complex traits. Nature genetics, 50(7):1041, 2018.

[40] Wen Zhang, Georgios Voloudakis, Veera M Rajagopal, Ben Reahead, Joel T Dudley, Eric E Schadt, Johan LM Bjorkegren, Yungil Kim, John F Fullard, Gabriel E Hoffman, et al. Integrative transcriptome imputation reveals tissue-specific and shared biological mechanisms mediating susceptibility to complex traits. bioRxiv, page 532929, 2019.

[41] Matthew Stephens. Dealing with label switching in mixture models. Journal of the Royal Statistical Society: Series B (Statistical Methodology), 62(4):795–809, 2000.

[42] Geoffrey McLachlan and Thriyambakam Krishnan. The EM algorithm and extensions, volume 382. John Wiley & Sons, 2007.

[43] Maya R Gupta, Yihua Chen, et al. Theory and use of the em algorithm. Foundations and Trends@ in S’ignal Processing, 4(3):223–296, 2011.

